# Programmable design of functional proteins from natural language

**DOI:** 10.1101/2024.08.01.606258

**Authors:** Fengyuan Dai, Shiyang You, Yudian Zhu, Yuan Gao, Lihao Fu, Xibin Zhou, Jin Su, Chentong Wang, Yuliang Fan, Qingsong Tan, Xiaoxiao Ma, Xianjun Deng, Letong Yu, Hui Qian, Yan He, Yitao Ke, Zhe Zhang, Chenchen Han, Xing Chang, Liangzhen Zheng, Sheng Wang, Yuan Ji, Kexin Hao, Longxing Cao, Yajie Wang, Anping Zeng, Shunzhi Wang, Chunjun Zhan, Jing Zhao, Tong Si, Jianming Liu, Hongyuan Lu, Fajie Yuan

**Affiliations:** School of Engineering, Westlake University, Hangzhou, 310024, China; Zhejiang University, Hangzhou, 310058, China; Bioscience and Biomedical Engineering Thrust, Systems Hub, The Hong Kong University of Science and Technology (Guangzhou), Guangzhou, Guangdong 511400, China; Department of Biochemistry, University of Washington Seattle, WA 98195, USA; State Key Laboratory of Quantitative Synthetic Biology, Shenzhen Institute of Synthetic Biology, Shenzhen Institutes of Advanced Technology, Chinese Academy of Sciences Shenzhen, 518055, China; School of Life Sciences, Westlake University, Hangzhou, 310024, China; Shanghai Zelixir Biotech Co., Ltd., Shanghai, 201203, China; State Key Laboratory of Druggability Evaluation and Systematic Translational Medicine, Tianjin Institute of Pharmaceutical Research, 306 Huiren Road, Tianjin, 300301, China; Zhejiang Key Laboratory of Low-Carbon Intelligent Synthetic Biology, School of Engineering, Westlake University, Hangzhou, 310024, China

## Abstract

Programming biological function—designing bespoke proteins to perform specified molecular tasks—is a foundational goal of molecular engineering. However, current design paradigms remain fundamentally limited: they typically require either natural proteins as starting points for optimization or manual reformulation of functional goals as geometric and sequence-level constraints to guide candidate generation. Here we introduce Pinal, a 16-billion-parameter foundation model that designs candidate proteins from natural-language descriptions of desired function. Trained on 1.7 billion synthetically annotated protein–text pairs, Pinal links functional intent to protein sequence and structure. In computational evaluations, generated candidates combined high predicted foldability with functional-description alignment and sequence diversity, providing a basis for prioritizing experimentally testable designs. We applied Pinal to four distinct design tasks—a fluorescent protein, a polyethylene terephthalate hydrolase, an alcohol dehydrogenase and a metabolic H-protein—and experimentally observed the intended function in each case, including catalytic activity for both designed enzymes. Crucially, without iterative experimental optimization, a Pinal-designed H-protein increased product titer by 1.7-fold relative to the corresponding native *E. coli* H-protein in a multi-enzyme CO_2_ fixation pathway. These findings support natural language as a high-level interface for candidate generation in protein design, enabling programmable exploration with reduced reliance on manually specified structural or sequence constraints.

---

The ambition to program biological function—creating bespoke proteins for applications in medicine, sustainability and industry—is a longstanding goal of molecular engineering. For decades, this quest has been shaped by two complementary philosophies: mimicking natural selection through directed evolution ^1–6^, or applying first principles of physics and chemistry to design proteins *de novo* using computational tools like Rosetta ^7–13^. Recent advances in artificial intelligence (AI), particularly deep generative models, have enabled an emerging design paradigm in which models learn from protein sequences ^14–16^, structures ^17–24^, and functional annotations ^25,26^ to generate proteins under diverse design objectives.

Yet, despite this progress, a central bottleneck persists. The dialogue between the human scientist and the protein design algorithm often remains mediated through formalized technical specifications, including structural constraints ^18,27–29^, site-level mutational instructions ^30,31^, and family-specific parameters ^25,32,33^. Translating a high-level functional goal—such as a plastic-degrading enzyme or a fluorescent imaging reporter—into a precise, machine-readable design problem therefore often requires substantial computational effort and specialized design expertise. This technical barrier limits broad access to protein design and constrains the space of functions that can be explored.

Here, we introduce Pinal—a generative protein foundation model that provides a natural-language interface for protein design (http://www.denovo-pinal.com/, Fig. 1). Pinal translates high-level functional descriptions into candidate protein sequences, thereby reducing reliance on manually specified structural templates or geometric constraints. Powered by a 16-billion-parameter architecture trained on a synthetic corpus of 1.7 billion protein-text pairs, Pinal exhibits strong scaling trends, with performance improving with both model and data scale. Across computational benchmarks, Pinal-generated candidates exhibited predicted folding metrics comparable to those of reference natural proteins, while showing markedly improved alignment with intended functional descriptions compared with related text- or prompt-conditioned protein-generation baselines. Further analyses revealed sequence-level diversity across multiple enzyme-family contexts and identified candidates with predicted structural novelty, suggesting that natural-language guidance can support broad exploration of the protein design space.

**Fig. 1:**
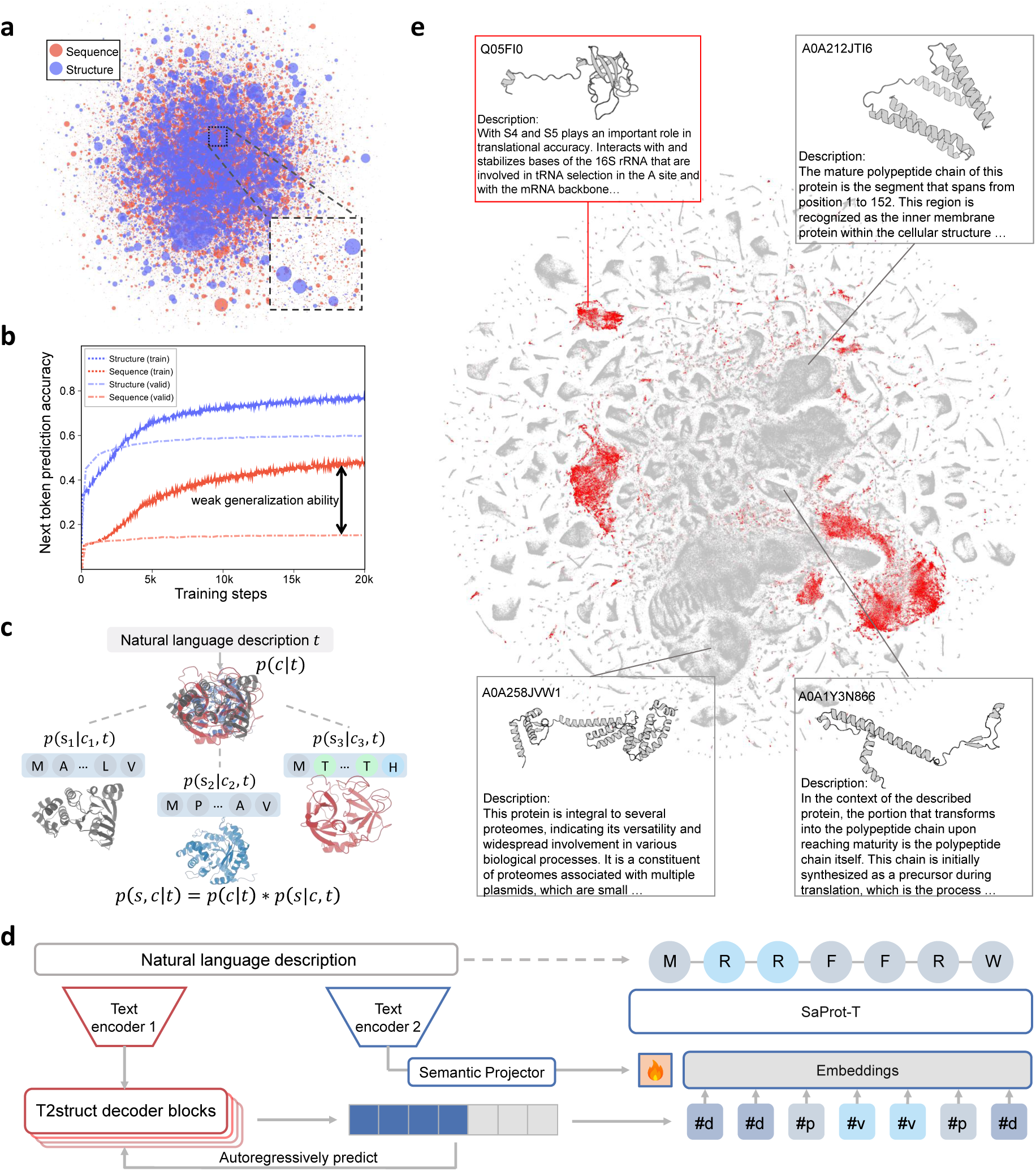
Pinal is a generative foundation model that uses a two-stage design process and large-scale training data to learn the text-to-protein link. **a:** Protein sequence versus structure space in the Swiss-Prot database. The 65,733 sequence clusters (30% identity) map to 21,368 structure clusters, demonstrating greater structural conservation. Each circle is a cluster; its size is proportional to the protein count. **b:** Next-token prediction accuracy on Swiss-Prot. Blue indicates text-to-structure generation and red text-to-sequence generation. The training and validation sets are split by 30% sequence identity. “train” and “valid” denote accuracy on each set. **c:** The chain rule decomposes text-to-protein design into two sequential stages mediated by an intermediate structural representation. **d:** Pinal model architecture. T2struct modules are outlined in red and SaProt-T modules in blue. Text Encoder 1 is initialized from pre-trained T5, and Text Encoder 2 from pre-trained PubMedBERT. **e:** Visualization of the distribution of human-reviewed Swiss-Prot data (545K proteins, 27M protein-text pairs, red) and our synthetic dataset (135M proteins, 1.7B protein-text pairs, gray) in a low-dimensional space, obtained by t-SNE reduction of the ProTrek embeddings. The synthetic dataset vastly expands protein design-space coverage, encompassing and extending beyond human-annotated data.

We evaluated Pinal across four protein design tasks: a green fluorescent protein (GFP), a polyethylene terephthalate hydrolase (PETase), an alcohol dehydrogenase (ADH), and a metabolic H-protein. Experimental validation identified functional designs across all four tasks, including catalytic activity for both designed enzymes. Notably, in the H-protein case, a candidate generated without any iterative experimental optimization achieved a higher product titer than the native *E. coli* H-protein when incorporated into a multi-enzyme CO_2_ fixation pathway. Together, these results establish Pinal as a practical route to protein design, lowering the barrier between functional intent and molecular implementation across diverse applications.

## 1 The Pinal network

The architecture of Pinal is motivated by the idea that protein structure can serve as a compact and functionally informative intermediate representation between natural-language specifications and amino acid sequences, as protein structures are often more conserved than their corresponding sequences ^34^. As an initial test of this representation choice, we performed a controlled preliminary comparison using SwissProt-scale protein-text pairs with high-quality, curated functional annotations (Fig. 1a,b, Methods 7.2). In this comparison, two identical encoder-decoder models were trained on the same protein-text pairs with the same optimization protocol and 30% sequence-identity split, differing only in the prediction target: amino-acid tokens or Foldseek 3Di structure tokens. The direct text-to-sequence model did not meaningfully exceed the uniform random-guessing baseline in validation next token prediction (NTP) accuracy (accuracy: ∼0.1 vs. random-guessing baseline: 0.05), whereas the text-to-structure model converged robustly (validation accuracy: ∼0.6) and, after translation to amino-acid sequences, yielded candidates with substantially higher predicted folding confidence as measured by ESMFold pLDDT (Supplementary Fig. 2). This contrast motivates a structure-mediated formulation of protein design, in which text first specifies a structural scaffold and the final sequence is then designed under both structural and textual constraints. Pinal is therefore formulated as a generative framework that models the conditional joint distribution of a sequence (*s*) and its structure (*c*) given text (*t*). We factorize the model distribution according to the probability chain rule: *p*(*s, c*|*t*) = *p*(*c*|*t*)*p*(*s*|*c, t*). This factorization naturally defines Pinal’s two-stage process (Fig. 1c, Extended Data Fig. 1): first, generating a structural scaffold (*c*) from the text-conditioned distribution *p*(*c*|*t*), and second, designing a sequence (*s*) conditioned on both the scaffold and the text, according to *p*(*s*|*c, t*).

The first stage is handled by T2struct, a text-to-structure module that translates a natural language prompt (*t*) into a structural blueprint (*c*). Critically, T2struct represents protein backbones not as continuous 3D coordinates, but as a 1D sequence of discrete 3Di structural tokens from Foldseek ^35^, where each token encodes the local tertiary interaction geometry of a residue with its nearest spatial neighbor. This representation reframes structure generation from a complex continuous coordinate prediction task into a discrete sequence-to-sequence learning paradigm. By doing so, it bypasses the architectural complexities associated with modeling 3D spatial symmetries (such as rotational and translational equivariance) and allows the framework to seamlessly benefit from the proven scalability of standard Transformer architectures ^36–38^. Furthermore, a key feature of this token-wise autoregressive approach is its ability to assign a computable log-probability, log *p*(*c*|*t*), to each generated sequence of structures—a capability that proves essential for the optimal sampling strategy described below. T2struct employs an encoder-decoder Transformer architecture (Methods 7.1): its encoder, a pre-trained T5 encoder model ^39^, is specialized for interpreting biological text, while its decoder, a GPT-2 style model trained from scratch, autoregressively predicts the 3Di token sequence (Fig. 1d), connected via a text-3Di cross-attention module.

With a structural blueprint in hand, the second stage is performed by SaProt-T, our text-conditioned sequence designer. This module receives the 3Di structure tokens generated by T2struct and outputs a viable amino acid sequence (*s*) conditioned on both the structure and the original textual prompt (Fig. 1d, right). SaProt-T is adapted from the original SaProt ^36^ model and incorporates a mechanism to ensure that the generated sequence satisfies the functional constraints specified in the text. To do so, we first encode the text using PubMedBERT ^40^. The resulting representation is projected to the model dimension and prepended to the input 3Di token embeddings. The combined representation is then processed by self-attention layers, allowing semantic cues from the language to be integrated with the geometric constraints of the structural scaffold. The model is trained with a masked language modeling objective to predict the correct amino acid at each structural token position (Extended Data Fig. 2, Methods 7.5.1).

The power of Pinal’s architecture lies not merely in the sequential execution of the two generation modules, but in their probabilistic unification to elegantly solve the global optimization problem defined in Eq. 1. This is because a naive, greedy approach—first finding the single most probable structure and then designing the best sequence for it—is mathematically suboptimal, as highlighted in Eq. 2. Such a myopic strategy risks overlooking a slightly less probable structure that, crucially, can accommodate a far superior functional sequence. Pinal circumvents this pitfall by employing a global sampling and re-ranking strategy, a process made possible by the fact that both of its two generative stages yield tractable log-probabilities 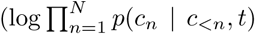 and log 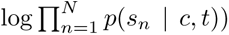. In practice, we first sample *K* structural scaffolds from T2struct’s distribution, *p*(*c*|*t*). For each of these *K* scaffolds, we then use SaProt-T to design its optimal corresponding sequence. Finally, the resulting (*K*) sequence–structure pairs are re-ranked using a per-position joint score derived from the two log-probability terms in Eq. 1. This allows Pinal to identify globally superior candidates that a purely sequential, greedy decision process would almost certainly miss (Methods 7.6, Extended Data Fig. 3).

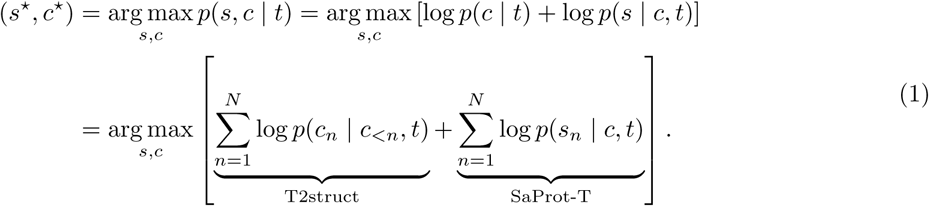

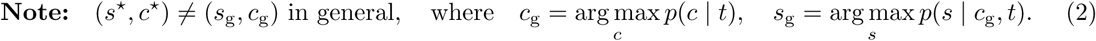

## 2 Scaling Pinal with massive synthetic data

While scaling laws have revolutionized general AI ^38,41^, their potential in natural language-guided protein generation is untapped due to critical data limitations. The premier human-annotated database, Swiss-Prot, contains merely 0.5 million proteins, far smaller than the billion-scale datasets powering modern language models. Moreover, its template-driven annotations do not reflect the complexity and diversity of queries common in biological research.

To overcome the constraints of scale and diversity, we developed an unprecedented synthetic training dataset containing 1.7 billion text-protein pairs. This comprehensive dataset encompasses over 100 billion word tokens across 135 million unique protein sequences and structures (Methods 7.3, Supplementary Table 2 and Table 4). The dataset construction involved a systematic, multi-faceted approach to generate high-quality natural language descriptions of proteins. For UniProt/TrEMBL entries, we employed the multimodal protein language model ProTrek ^42^ to predict diverse functional descriptions from multiple perspectives, creating the TrEMBL50-ProTrek subset. These multi-perspective descriptions were then synthesized using large language models (LLMs) to produce coherent, integrated functional annotations, resulting in the TrEMBL50-ProLLM dataset. For Swiss-Prot proteins, we applied a parallel strategy, utilizing LLMs to consolidate multiple functional descriptions into comprehensive annotations (SwissProt-LLM). Additionally, we preserved the original structured Swiss-Prot entries (Swiss-Annot) while generating LLM-paraphrased versions to enhance linguistic diversity (Swiss-Aug). To ensure robust coverage across varying description lengths and annotation types, we incorporated nearly 800 million keyword-protein pairs from the InterPro database. All protein structural information was sourced from the AlphaFold database (AFDB), providing a unified structural foundation for the entire dataset.

We present the first empirical demonstration of data scaling effects in natural language-guided protein design. Our scaling experiments revealed two key findings that illuminate the relationship between training data size and model performance. First, on challenging test sets with less than 30% sequence identity, model performance—as measured by ProTrek score (Methods 7.7.4, Extended Data Fig. 4) and GT-TMscore—exhibited substantial and consistent improvement with increasing training data size (Fig. 2b, right) (detailed evaluation settings and measures are described in the next section). Second, on standard test sets with less than 50% sequence identity, the ProTrek score demonstrates a steady upward trend throughout the scaling range, while GT-TMscore values plateau rapidly between 100 million and 1 billion word tokens before showing modest gains with further scaling (Fig. 2b, left). Furthermore, beyond these alignment metrics, increasing the scale of the synthetic training corpus expanded the accessible design space, enabling the generation of distinctly novel and diverse proteins across both sequence and structural dimensions (Supplementary Fig. 3, Supplementary Method 2). These findings demonstrated that synthetic data benefits manifested most prominently under demanding generalization scenarios, where the complexity of the task necessitated larger-scale training data to achieve meaningful performance improvements (see Supplementary Note 1 for further discussion of data scaling effects).

**Fig. 2:**
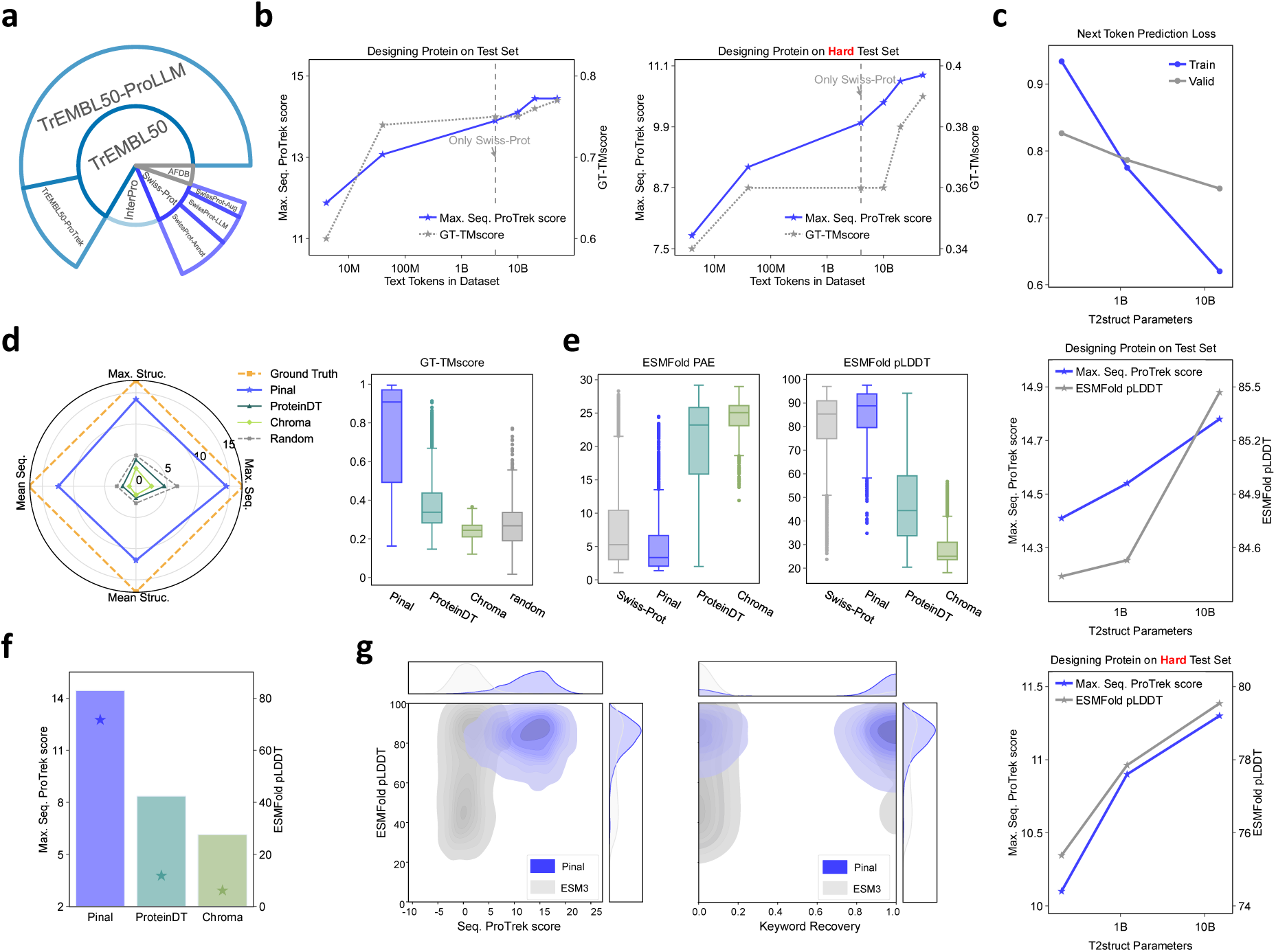
Scaling T2struct and benchmarking Pinal’s design performance. **a:** Training data sources for Pinal. **b:** Scaling of Pinal with training data size. Points left of the dashed line indicate training on Swiss-Prot only; points right indicate Swiss-Prot plus synthetic data. The left y-axis shows the best sequence- to-text ProTrek score from five designed candidates per functional prompt, and the right y-axis shows the average GT-TMscore of designed candidates relative to the target protein. The x-axis shows training text tokens on a log scale. Owing to computational cost, this ablation uses the medium-sized 1.2B-parameter T2struct model within Pinal. **c:** Scaling of the T2struct module within Pinal from 200M to 15B parameters, with log-scaled x-axis. The three plots are arranged vertically. **d:** Alignment between designed and target proteins in the test set. The left panel is a radar chart of four ProTrek-based alignment metrics. Max. Seq. and Mean Seq. denote the maximum and mean sequence-to-text scores across five designs per prompt; Max. Struc. and Mean Struc. denote the corresponding structure-to-text scores. Each method forms a polygon; values farther from the center and larger enclosed areas indicate better overall performance. The right panel shows the distribution of GT-TMscore values for *n* = 2, 500 proteins per method; higher values indicate better agreement with the target description or structure. **e:** Foldability of designed proteins evaluated using ESMFold PAE and pLDDT for *n* = 2, 500 proteins per method. In d and e, box plots show the median (centre line), 25th–75th percentiles (box), the most extreme values within 1.5× the interquartile range (whiskers), and outliers (points). **f** : Foldability and textual alignment of designed proteins conditioned on short sentences. Bars show ESMFold pLDDT foldability (right axis), and stars show Max. Seq. ProTrek textual alignment (left axis). **g:** Comparison with ESM3. Darker colors in the upper-right region indicate stronger model performance.

We further investigated Pinal’s model scaling trends. Recent studies on protein language models (PLMs) have observed that scaling model parameters may not yield substantial benefits for corresponding protein prediction tasks ^43,44^. Similarly, on the ProteinGym benchmark ^45^, ESM2’s 15 billion parameter version performs even worse than the 650 million parameter variant. We demonstrated Pinal’s model scaling capabilities by training T2struct at three distinct scales: 200 million, 1.2 billion, and 15.5 billion parameters (Supplementary Table 3). As model capacity increased, we observed substantial reductions in validation loss (Fig. 2c, upper). More critically, this improvement translated into superior practical performance: larger models generate proteins with markedly higher predicted foldability (pLDDT scores) and enhanced alignment with language instructions (ProTrek scores) across both standard and stringent test sets (Fig. 2c). Together with our data scaling findings, these experiments established that both model scaling and data scaling serve as fundamental drivers of Pinal’s performance achievements.

## 3 Computational benchmarks of Pinal-designed proteins

Here, we systematically evaluated proteins designed by Pinal, assessing both their structural correctness and functional relevance. Structural correctness was quantified using folding confidence (pLDDT), Predicted Aligned Error (PAE), and similarity to native structures (TM-score). To address the challenge of evaluating functional relevance—a critical gap in the nascent field of language-guided protein design—we employed the ProTrek protein-to-text matching score (Methods 7.7.4, Extended Data Fig. 4), which measured the semantic distance between protein sequence/structure and their functions.

We benchmarked Pinal against the limited number of existing approaches to language-guided protein design (Methods 7.7, Extended Data Fig. 5a, Supplementary Method 1). Among these, ProteinDT ^46^ was the only prior model explicitly developed for this task, but it remained an early-stage exploration, with both model scale and dataset size orders of magnitude smaller than those of Pinal. The other models also exhibited clear limitations: Chroma ^47^, according to the authors’ own report, showed only occasional alignment between designed proteins and intended prompts, whereas ESM3 ^38^, a concurrent work by preprint date, was limited to simple keyword inputs and lacked the ability to interpret complex natural-language instructions.

We designed a multi-faceted evaluation framework comprising three distinct settings: (1) long, descriptive sentences that mimic the complex queries of biologists; (2) short, function-specific phrases to test core generative accuracy; and (3) specialized keywords to enable a direct, rigorous comparison with ESM3 (see Methods 7.7.2). In tasks involving both long and short sentences, Pinal markedly outperformed ProteinDT and Chroma. Designs from ProteinDT and Chroma failed to align with their textual prompts, yielding ProTrek and TM-scores similar to those of a random baseline (Fig. 2d). Furthermore, a substantial fraction of their generated sequences failed to fold into plausible structures, as indicated by low pLDDT and high PAE values. In stark contrast, proteins designed by Pinal not only exhibited high textual & functional alignments, with ProTrek scores and TM-scores close to those of real natural proteins, but also possessed structural features (pLDDT, PAE) indicative of stable folding, comparable to those of well-characterized proteins in the Swiss-Prot database (Fig. 2e). Pinal’s superior performance can be attributed to three key factors: an advanced architecture, a substantially expanded parameter space (tens of times larger than those of previous models), and a much larger training dataset (thousands to tens of thousands of times larger than those used by ProteinDT and Chroma). This performance advantage was consistently observed across both complex, multi-aspect descriptions and concise, function-specific prompts (Fig. 2f, Extended Data Fig. 6).

To further assess Pinal’s capabilities, we compared it with ESM3 in a keyword-conditioned generation setting. To ensure a stringent and equitable comparison, we evaluated Pinal on ESM3’s “home ground”: generating proteins from InterPro database keywords, which were used as conditioning information in ESM3 training, and assessing performance using ESM3’s established keyword-recovery metric. Even in this ESM3-favorable setting, Pinal showed substantially better performance (Fig. 2g). Pinal-generated proteins exhibited higher predicted foldability overall, whereas ESM3 outputs showed greater variability. Most notably, Pinal achieved substantially better functional alignment, with a mean keyword recovery rate of 77%, compared with approximately 12% for ESM3.

In addition to these findings, a recent independent benchmark study ^48^ has confirmed Pinal’s effectiveness, showing it achieved the best performance on 11 out of 14 metrics focused on sequential plausibility, structural foldability, and language alignment. Compared to models like ProteinDT, Chroma, ESM3, and PAAG ^49^, Pinal demonstrated performance gains ranging from several-fold to tenfold on multiple metrics.

## 4 Pinal explores sequence diversity and predicted structural novelty

Having established that Pinal designs are predicted to be structurally plausible and aligned with their input prompts, we next evaluated whether the model could explore diverse and potentially novel regions of protein design space. We first examined whether Pinal could generate sequence-diverse proteins across established enzyme families. To this end, we prompted Pinal to generate sequences for four enzyme families: CoA ligase, glucokinase, aldehyde dehydrogenase, and carbonic anhydrase. These families span distinct catalytic mechanisms and functional contexts, providing a test of Pinal’s ability to generate sequence variation across different enzyme classes. Maximum-likelihood sequence-tree analysis showed that the analyzed Pinal-designed sequences were distributed across multiple branches in trees inferred jointly with curated natural homologues, rather than collapsing into a single compact cluster (Methods 7.8). In several cases, designed sequences were interspersed with natural homologues, suggesting overlap with multiple regions of the corresponding natural sequence space. Together, these results suggest that Pinal can generate sequence-level diversity across diverse enzyme-family contexts (Fig. 3a).

**Fig. 3:**
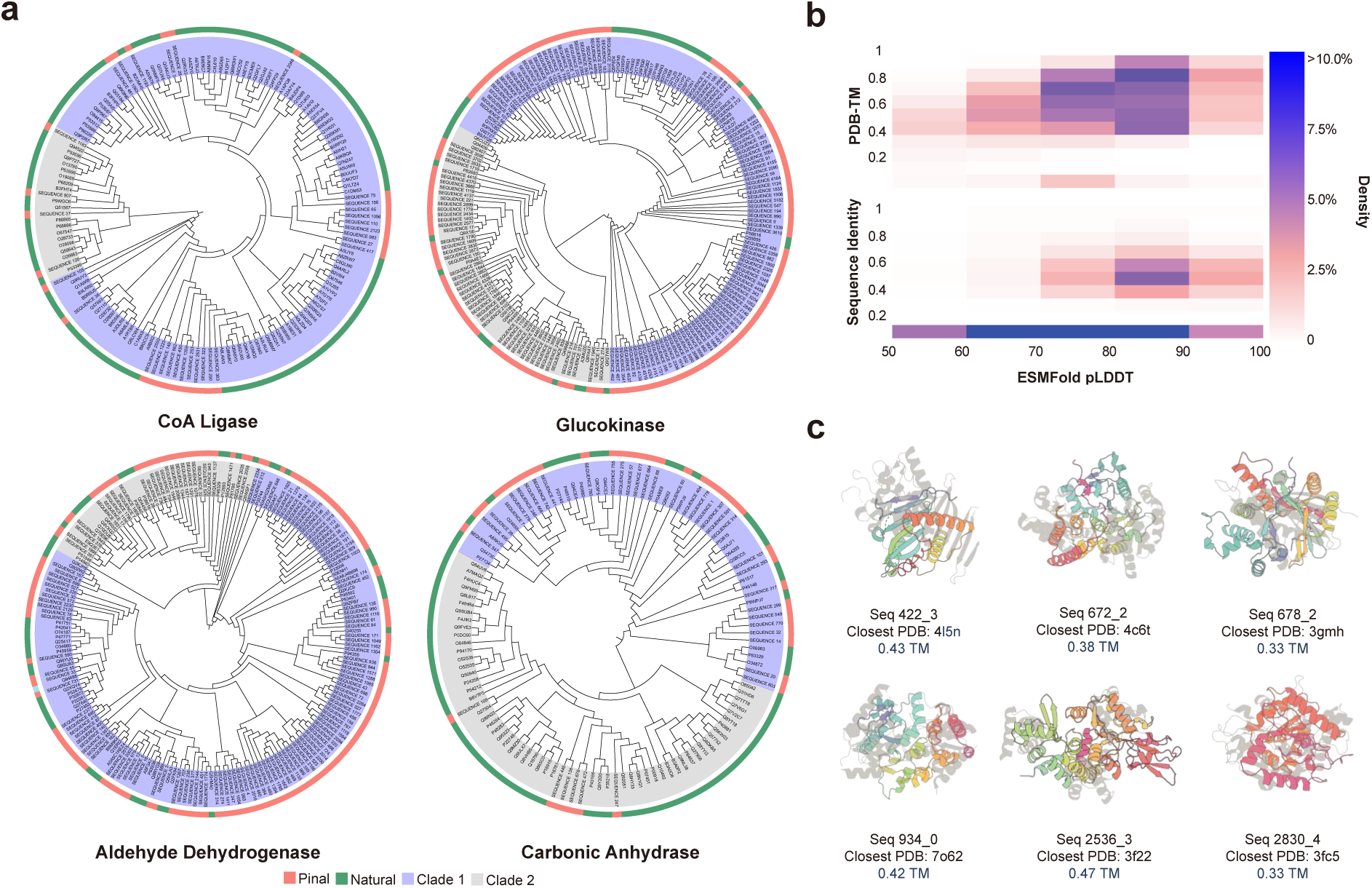
Pinal generates a diverse landscape of novel proteins. **a,** Circular maximum-likelihood sequence trees of Pinal-designed sequences and natural reference sequences from four enzyme families: CoA ligase, glucokinase, aldehyde dehydrogenase, and carbonic anhydrase. The outer ring indicates sequence origin (salmon, Pinal; green, natural reference), whereas the inner shading marks two major sequence clades (lavender, Clade 1; grey, Clade 2). **b,** Heatmaps illustrate the relationship between predicted structural confidence (ESMFold pLDDT, X-axis) and protein novelty. The top panel shows structural novelty (TM-score to known proteins in PDB, Y-axis), and the bottom panel shows sequence novelty (sequence identity to known proteins in UniRef100, Y-axis). Darker shades indicate higher density of designed proteins. **c,** Representative examples of novel protein structures designed by Pinal (rendered in rainbow). For each design, the closest structural homologue (rendered in grey) in the PDB is shown, along with a low TM-score, providing visual confirmation of Pinal’s ability to generate previously unknown protein folds.

We then assessed Pinal’s generative ability for structural novelty by designing proteins that are currently experimentally uncharacterized. To investigate this, we prompted the model with functional descriptions of proteins that lack any known structural homologue in the PDB (Methods 7.7.3). The results in Fig. 3b indicate that a subset of Pinal’s designs fall within the high-confidence foldability region (high pLDDT), while exhibiting low sequence identity (*<*30%) and low structural similarity (TM-score *<* 0.5) to known proteins. These results suggest that Pinal can generate candidate proteins that are predicted to be foldable while showing limited similarity to sequences and structures represented in current databases.

To test whether computationally selected novel designs could be experimentally realized, we selected 20 Pinal-designed proteins whose predicted structures showed low similarity to known PDB structures (maximum TM-score *<* 0.5) for experimental expression testing. Seventeen of these 20 designs (85%) were successfully expressed in soluble form in *E. coli* BL21(DE3), as assessed by SDS-PAGE analysis of the soluble fraction (Methods 7.10.1, Supplementary Fig. 5 and Supplementary Table 7). Although soluble expression does not establish biochemical function or experimentally confirm the predicted structures, it indicates that many of these low-similarity designs can be physically produced as soluble proteins under the tested conditions. These results suggest that Pinal can generate protein designs that are structurally distant from proteins currently represented in the PDB, supporting its capacity to explore novel regions of protein structure space (Fig. 3c).

## 5 Experimental realization of diverse, active proteins

We next asked whether Pinal-generated sequences could move beyond predicted foldability and soluble expression to exhibit measurable biochemical function. To this end, we designed a validation framework targeting four proteins of scientific and industrial importance (Fig. 4). Each target probed a distinct class of biochemical complexity: the intrinsic photophysics of a GFP; the enzymatic catalysis of a synthetic polymer by a PETase; the cofactor-dependent mechanism of an ADH; and the molecular coordination of an H-protein within a multi-protein system. For each target, Pinal generation was guided by a high-level functional text prompt (Supplementary Table 8), with no structural or sequence templates as input. Across the main validation set of these case studies, we experimentally validated 12 active Pinal-designed proteins; these designs shared 30–60% sequence identity with functionally annotated proteins in Swiss-Prot (Supplementary Table 6). Together, these results provide a rigorous test of Pinal’s potential as a general-purpose, programmable protein design engine.

**Fig. 4:**
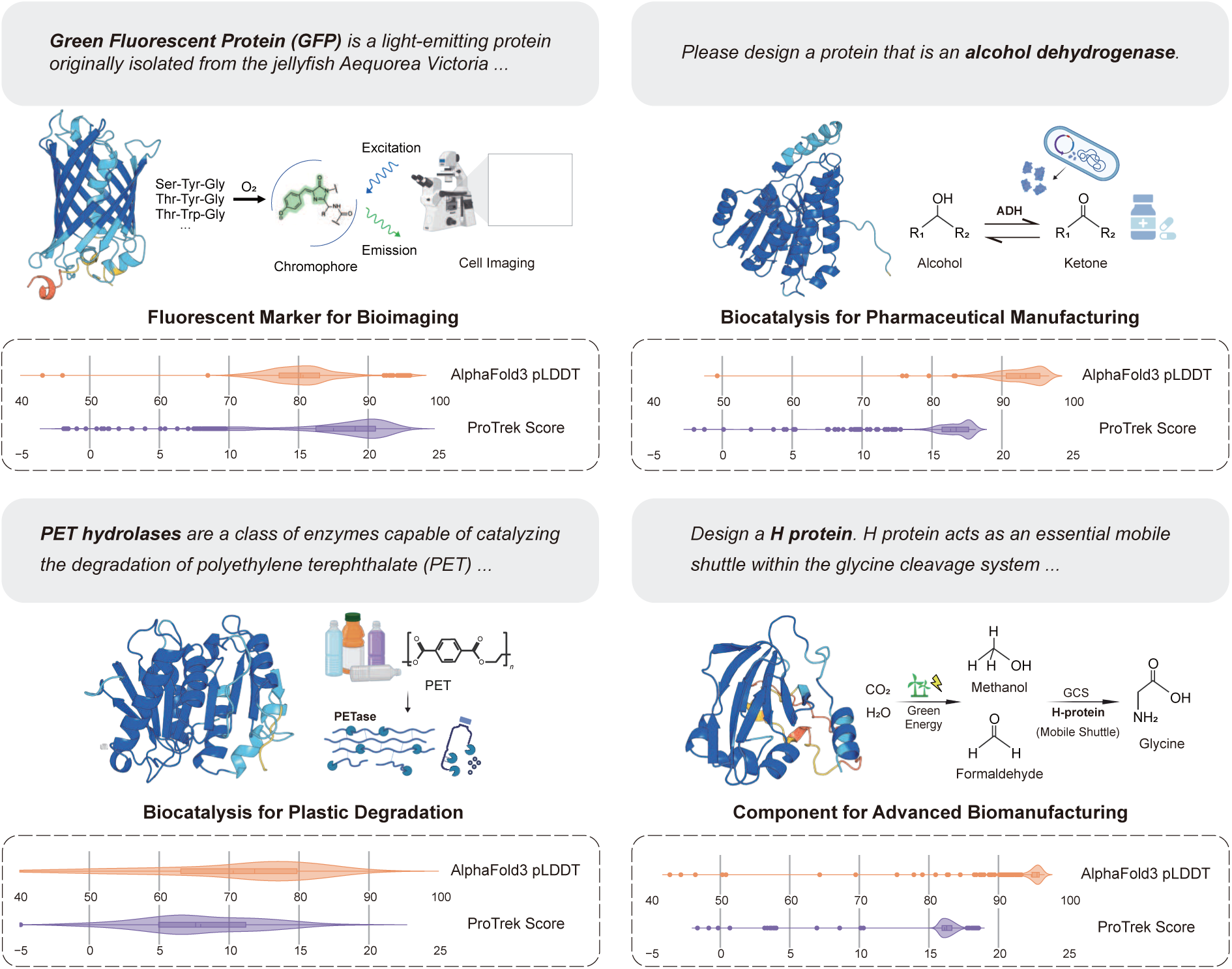
Pinal translates diverse functional prompts into high-quality protein candidates. The conceptual application, natural language prompt, and final design are presented for GFP, ADH, PETase, and H-protein. Violin plots show the high quality of the *n* = 1, 000 candidates per protein class, with high AlphaFold3 pLDDT scores indicating structural confidence and high ProTrek scores demonstrating close functional alignment with the prompt. Violin widths represent kernel-density estimates. Box plots show the median (centre line), 25th–75th percentiles (box), the most extreme values within 1.5× the interquartile range (whiskers), and outliers (points). Figure 4 was in part created with Biorender.com https://BioRender.com/gn9qk93.

Our first challenge for Pinal was the computational design of a GFP-like fluorescent protein, a demanding functional-design problem requiring correct *β*-barrel folding and maturation of an internal chromophore ^50^. Using only a high-level text prompt describing the desired GFP-like fluorescence as the initial design input, Pinal generated a diverse computational library of 50,000 sequences (Supplementary Table 8 and Table 13). To enable tractable experimental validation, we used an empirical, multi-criteria triage procedure to prioritize a small subset of 29 candidates containing putative chromophore-forming residues (Extended Data Fig. 7a, Methods 7.11.1), without treating the remaining untested designs as inactive. These prioritized designs shared *<*35% sequence identity with known fluorescent proteins. Screening this initial subset identified one active variant, pinalGFP, which produced green fluorescence (460 nm excitation, 510 nm emission) (Fig. 5a, Supplementary Table 6 and Table 9). Its predicted structure exhibited the characteristic GFP-like *β*-barrel architecture (Fig. 5b). This fluorescence was abolished upon mutation of the chromophore residues (Fig. 5c). As a reference point for an unoptimized first-pass GFP design, we included the previously reported ESM3-designed variant esmGFP-b8 ^38^, which, unlike the Pinal designs, was generated using explicit residue-level and structural conditioning. Under the same 24-h assay condition, pinalGFP exhibited detectable green fluorescence, whereas esmGFP-b8 showed little or no fluorescence within this assay window, consistent with its reported slow maturation (Fig. 5c). Purified pinalGFP displayed excitation and emission spectra analogous to those of natural avGFP (Extended Data Fig. 7c). However, analytical gel filtration revealed a more complex oligomeric state compared to the primarily dimeric avGFP as a natural variant (Extended Data Fig. 7d), highlighting a clear avenue for future structural optimization.

**Fig. 5:**
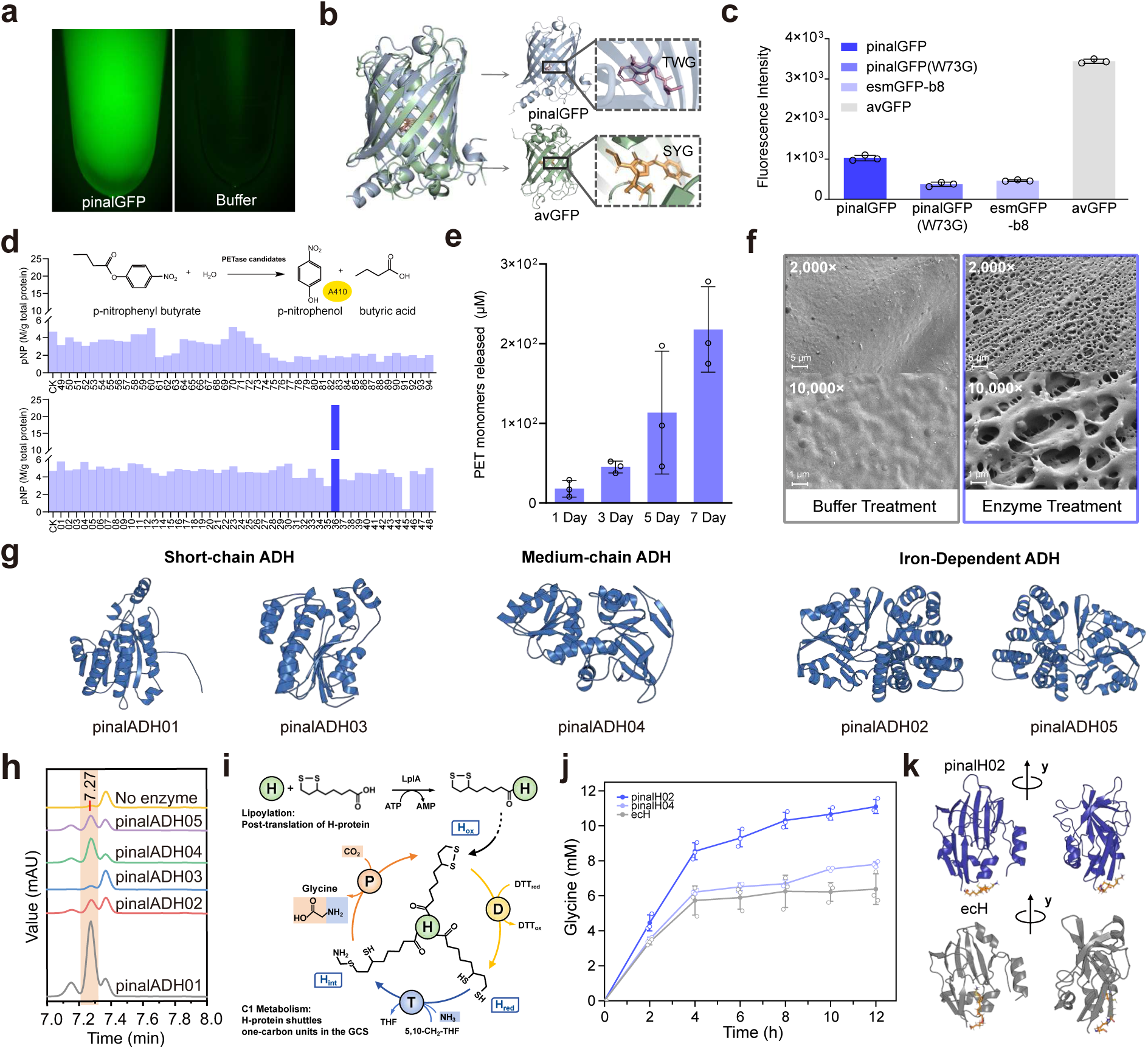
Experimental validation of Pinal-designed proteins across four distinct functional classes. **a,** Fluorescence microscopy image of purified pinalGFP (20 mg ml^−1^) acquired using the 488/520 filter. This experiment was independently repeated three times with similar results. **b,** AlphaFold3-predicted pinalGFP (blue) aligned with avGFP (green; PDB 2WUR), showing the conserved *β*-barrel architecture, with chromophore residues T72–W73–G74 and S65–Y66–G67, respectively, highlighted. **c,** Fluorescence of purified pinalGFP, the W73G chromophore mutant, esmGFP-b8 and avGFP. Synthetic variants were measured at 1 mg ml^−1^ and avGFP at 1 *µ*g ml^−1^. Data are mean ± s.d. (n = 3 independent reactions). **d,** Screening of 94 Pinal-designed PETase candidates for p-nitrophenyl butyrate (pNPB) esterase activity, with pNP concentration normalized to total protein amount; pinalPETase36 is highlighted as the candidate with the highest activity. **e,** High-performance liquid chromatography (HPLC) quantification of terephthalic acid (TPA) and mono-(2-hydroxyethyl) terephthalate (MHET) released from pretreated PET by purified pinalPETase36 over 7 days. Data are mean ± s.d. (n = 3 independent reactions). **f,** Scanning electron microscopy (SEM) images of pretreated PET films treated with pinalPETase36 or buffer. This experiment was independently repeated three times with similar results. **g,** Predicted structures of five experimentally validated alcohol dehydrogenase (ADH) designs spanning short-chain, medium-chain and iron-dependent long-chain subfamilies. **h**, HPLC chromatograms showing substrate conversion by the five purified ADHs relative to a no-enzyme control. **i,** H-protein lipoylation and the reverse glycine cleavage system (GCS). H protein accepts the aminomethyl group from T protein and transfers it to P protein during glycine synthesis from CO_2_. LplA, lipoate-protein ligase A; P, glycine decarboxylase; T, aminomethyltransferase; DTT*_ox_* and DTT*_red_*, oxidized and reduced dithiothreitol; 5,10-CH_2_-THF, N^5^,N^10^-methylene-tetrahydrofolate; H*_int_*, aminomethylated lipoylated H protein; H*_ox_* and H*_red_*, oxidized and reduced lipoylated H protein, respectively. **j,** Glycine production with pinalH02, pinalH04 and native *E. coli* H protein (ecH). Data are mean ± s.d. (n = 3 biologically independent reactions). **k,** pinalH02 and ecH conformations at 200 ns in molecular dynamics (MD) simulations; the aminomethyl lipoate arm of ecH is locked in the hydrophobic cavity.

Next, we escalated the challenge from designing a functional protein to creating a novel enzyme for catalysis. To this end, we targeted a PETase to degrade the recalcitrant plastic polyethylene terephthalate (PET). Guided by a text prompt specifying this catalytic function, Pinal generated 200,000 candidate sequences, which were then filtered for predicted foldability and semantic alignment with the prompt, yielding 2,416 candidates (Extended Data Fig. 8a). From this refined set, we selected the top 94 proteins for gene synthesis, ranked by their text-protein semantic alignment (ProTrek scores). Following expression in *E. coli*, we screened the crude cell lysates of these 94 candidates using a p-nitrophenyl butyrate (pNPB) esterase assay. This primary screening identified one standout variant, pinalPETase36, that exhibited notable catalytic activity toward pNPB (Fig. 5d, Supplementary Table 6 and Table 11, Methods 7.12.2). Purified pinalPETase36 displayed a melting temperature (Tm) of 56.84 ± 0.29 ^◦^C. To verify its capacity for actual polymer degradation, we evaluated its activity on solid PET substrates at 40 ^◦^C and pH 8.0. High-performance liquid chromatography (HPLC) analysis over a 7-day period (sampled on days 1, 3, 5, and 7) revealed a progressive accumulation of the degradation products, terephthalic acid (TPA) and mono-(2-hydroxyethyl) terephthalate (MHET)^51^ (Fig. 5e, Methods 7.12.3). This depolymerization was corroborated by scanning electron microscopy (SEM), which revealed distinct micro-pores and pitting on the pinalPETase36-treated PET substrates compared to the smooth surface of buffer-treated controls (Fig. 5f, Methods 7.12.4). Collectively, these chemical and physical analyses demonstrate that pinalPETase36 functions as an active PET-degrading enzyme.

Having demonstrated the design of a novel PETase, we next tested Pinal’s ability to navigate the complexity of a large enzyme family and generate diverse solutions across its distinct subfamilies. For this, we targeted ADH, a versatile class of biocatalysts for producing chiral alcohols that are essential building blocks for pharmaceuticals ^52^. Pinal was provided with five paraphrased general prompts encoding the same ADH-design request (Supplementary Table 8), imposing no constraints on metal dependence, subfamily or chain length. In total, 2,500 initial candidates were generated across these prompts. Feasibility filtering and prioritization yielded a pool of 1,000 high-confidence sequences, from which eight were selected for experimental validation (Extended Data Fig. 9a, Methods 7.13.1, Supplementary Table 10 and Table 13). Of these, five were successfully expressed and purified (Supplementary Table 6). These five purified designs spanned three distinct ADH subfamilies: Fe^2+^-dependent long-chain (pinalADH02, pinalADH05), medium-chain (pinalADH04), and short-chain variants (pinalADH01, pinalADH03) (Fig. 5g, Extended Data Fig. 9b). In silico analysis confirmed that the predicted folds, cofactor-binding features and, where applicable, metal-coordination geometries were consistent with their assigned subfamilies. HPLC-based activity assays confirmed that all five purified designs showed detectable ADH activity at 37 ^◦^C (Fig. 5h). This result demonstrates Pinal’s ability to capture the core functional principles of distinct enzyme subfamilies, allowing it to generate active catalysts across different protein architectures.

As the culminating test of Pinal’s capabilities, we challenged it to move beyond single-enzyme function and design a protein requiring precise coordination within a complex multi-protein system. We targeted the H-protein, a key shuttle protein within the glycine cleavage system ^53,54^, whose function requires both correct post-translational lipoylation and dynamic interactions with its partner enzymes to become active (Fig. 5i). From a single functional prompt, Pinal generated 50,000 initial candidates; feasibility filtering and prioritization reduced this pool to around 1,000 sequences, from which 20 were selected for gene synthesis and experimental validation (Extended Data Fig. 10a, Supplementary Table 6, Table 12 and Table 13). The designs yielded a high success rate, with 14 of the 20 candidates (70%) showing soluble expression. When tested for CO_2_ fixation in a reconstituted assay, five of the designed proteins were active (Extended Data Fig. 10b, Supplementary Table 6 and Table 12, Methods 7.14.1). The top-performing design, pinalH02, drove glycine synthesis to a final titer of 11.1 mM, a 1.7-fold increase over the native *E. coli* H-protein (ecH) (Fig. 5j). Molecular dynamics simulations suggested a possible structural basis for this improved performance. While the native ecH locks its reactive lipoamide arm in a protective cavity for regulatory purposes, the arm of pinalH02 is largely liberated. This key architectural difference may allow pinalH02 to bypass the native protein’s rate-limiting step, facilitating more efficient intermediate shuttling and leading to enhanced carbon fixation (Fig. 5k, Extended Data Fig. 10c-f). Together, these data indicate that Pinal-designed proteins can be incorporated into reconstituted multi-protein enzymatic pathways, with some designs matching or exceeding the activity of native components. Additional follow-up H-protein validation data are provided in the Supplementary Information (Supplementary Fig. 7).

## 6 Discussion

Pinal’s significance lies not only in its performance, but also in reframing protein design as a problem that can be specified through functional language. For decades, the field has largely required specialized expertise to translate desired functions into low-level structural or sequence constraints and to navigate workflows grounded in computation and biophysics. By interpreting high-level functional descriptions within a generative framework, Pinal reduces reliance on manual constraint engineering and points toward a more programmable paradigm for protein engineering.

Our study further highlights two principles that appear to be important for this setting. First, directly mapping the combinatorial space of language onto protein sequence is highly challenging. Our decoupled two-stage framework, in which protein structure serves as an intermediate semantic and geometric representation (t → c → s), appears to make this problem more tractable in practice. Second, scale matters. The performance of our 16-billion-parameter model, trained on 1.7 billion synthetic protein–text pairs, suggests that the scaling behaviours observed in general AI extend to language-guided protein design.

The successful experimental validation of four distinct protein classes supports the breadth of this approach, including the generation of catalytically active enzymes directly from textual descriptions. Moreover, the observation that a Pinal-designed H-protein outperformed the native *E. coli* H-protein in a multi-enzyme CO_2_ fixation pathway suggests that language-guided generation can access alternative functional solutions that surpass a native counterpart in a defined application context. Together, these results indicate that high-level functional descriptions can serve as a practical interface for specifying protein-design objectives and obtaining experimentally testable candidates.

Several limitations remain. Pinal’s capabilities are constrained by the granularity and coverage of available biological annotations, which currently limit fine-grained quantitative control over specified properties. For example, prompting for a precise emission wavelength in a fluorescent protein remains challenging. In addition, although our experiments validate the efficacy of the approach, the candidates advanced to experimental testing were prioritized conservatively using available in silico proxies, including AlphaFold confidence, ProTrek scores, and, where appropriate, motif-based criteria. These heuristics can improve practical hit rates, but they do not imply that candidates outside such filters are non-functional. Rather, they reflect the current lack of universally reliable computational metrics for evaluating protein function *a priori*. As a result, such prioritization may restrict exploration to a limited region of the design space. Coupling language-guided generation to high-throughput functional screening will therefore be important for more comprehensively probing the accessible design space and the limits of Pinal.

More broadly, our work suggests a path towards protein design systems specified increasingly in terms of function rather than manual constraint engineering. As biological datasets become richer and AI systems become more capable of integrating mechanistic reasoning with generative design, language-guided models may support increasingly open-ended exploration of protein function. In this sense, Pinal represents a step toward a programmable interface for biology, in which high-level intent can be translated into experimentally testable molecular candidates.

## 7 Methods

### 7.1 Detailed model architecture

Pinal’s generative capability is realized through a two-stage network composed of two main modules: T2struct and SaProt-T.

T2struct is an encoder-decoder Transformer that translates natural language descriptions into structural (3Di) token sequences (Extended Data Fig. 1a). The text encoder, initialized from a pre-trained T5 model ^39^, processes the input prompt to extract a high-dimensional text representation. This representation subsequently guides an autoregressive decoder, which is built upon a GPT-2-style architecture ^55^ but augmented with cross-attention mechanisms. Specifically, the decoder consists of *N* stacked blocks, each integrating three sequential sub-modules: (1) a structural context module employing multi-head self-attention to capture dependencies within the preceding structural tokens; (2) a text-conditioning module utilizing multi-head cross-attention to inject guidance from the encoded text representation; and (3) a feature refinement network consisting of a pointwise feed-forward layer. Each sub-module is followed by layer normalization. During training, T2struct is optimized using a teacher-forcing strategy: when predicting the next 3Di structural token, the autoregressive decoder is conditioned on the actual preceding tokens from the training data (rather than its own past predictions), while simultaneously being guided by the text representation. During inference, the model operates autoregressively, predicting each subsequent 3Di structural token based on its own previously generated sequence and the corresponding text representation.

The second stage of the Pinal network, SaProt-T, is an architectural adaptation of the SaProt model designed to generate a protein sequence from a structural scaffold and a text prompt (Extended Data Fig. 1b). To condition the generation on language, a separate text encoder (a pretrained PubMedBERT ^40^ model) produces a text representation, which is then passed to a semantic projector—a trainable linear layer—to map the text embedding into the protein embedding space. This projected text vector is then prepended to the sequence of input structural token embeddings to form a text-protein joint representation. This joint representation is subsequently processed by a stack of *N* SaProt-T blocks, each composed of self-attention and feed-forward modules, to predict the final amino acid sequence in a single forward pass.

### 7.2 Preliminary experiments of text-to-sequence and text-to-structure generation

In our approach, we represent protein structures as one-dimensional sequences of Foldseek 3Di tokens ^35^, framing structure generation as a discrete sequence-modeling task. Conveniently, because this 20-state structural alphabet matches the size of the standard amino acid vocabulary, it provides a unique opportunity to contrast text-to-structure and direct text-to-sequence generation under an identical architectural framework. Leveraging this shared formulation, we trained two identical encoder-decoder Transformer models (comprising a pre-trained T5-base encoder and a randomly initialized GPT-2 decoder, totaling 224M parameters) on the SwissProt-Annot dataset (Methods 7.3.1). Both models were trained for 20,000 steps under identical optimization settings, with training and validation sets rigorously split at a 30% sequence identity threshold. The sole distinction between the two setups was the prediction target: one model directly predicted amino acid sequences, while the other predicted 3Di structural tokens.

Given the 20-token vocabulary for both tasks, the theoretical uniform-random baseline accuracy is 0.05. During training, the direct text-to-sequence model failed to meaningfully surpass this random baseline on the validation set, indicating that mapping natural language directly to amino acid sequences is prohibitively difficult under the current data-scale setting. In contrast, the text-to-structure model converged robustly (Fig. 1b), reaching a validation accuracy of approximately 0.6. To move beyond the next token prediction (NTP) accuracy, we further evaluated the structural plausibility of the generated candidates across training checkpoints using ESMFold pLDDT (Supplementary Fig. 2). Because 3Di token sequences cannot be directly evaluated for structural plausibility by standard folding algorithms, they needed to be translated into amino acid sequences using SaProt ^36^. Consistent with the NTP accuracy results, the structure-mediated approach generated candidates with higher predicted folding confidence, whereas the direct text-to-sequence baseline produced candidates with substantially lower predicted folding confidence.

It is important to note that these results do not establish the intrinsic infeasibility of direct text-to-sequence generation. Rather, they show that, in this controlled SwissProt-scale paired-data setting, text-to-3Di-token generation is substantially more learnable than direct text-to-amino-acid generation under the same architecture and optimization setup. Recent studies, including ProLLaMA ^56^, ProtDAT ^57^ and ProDVa ^58^, have also reported promising progress in direct annotation- or prompt-conditioned amino-acid sequence generation, using strategies such as large-scale protein language model pretraining followed by instruction tuning, specialized cross-attention mechanisms, or dynamic protein vocabularies based on retrieved sequence fragments. These advances indicate that direct text-to-amino-acid generation can benefit from substantially larger pretraining corpora or tailored model designs. At the same time, whether the modality gap observed here would persist after scaling the paired text–sequence data and compute within our framework by orders of magnitude (e.g., 1, 000× or 10, 000×) remains an open question. Such a full-scale ablation would require resources comparable to training Pinal itself and therefore falls outside the scope of this preliminary study.

### 7.3 Dataset construction

Training a large-scale, text-guided protein design model requires extensive and diverse datasets. However, a fundamental challenge exists: high-quality, manually curated databases such as Swiss-Prot are limited in scale (containing 0.5 million proteins), making them insufficient for training billion-parameter models. Conversely, large sequence databases like UniProt/TrEMBL ^59^, while containing hundreds of millions of proteins, lack reliable text annotations.

To overcome this data gap, we devised a comprehensive, multi-pronged strategy to construct a hybrid dataset. Our approach encompasses two primary components: (1) augmenting the high-quality Swiss-Prot annotations using large language models (LLMs) to improve their diversity and comprehensiveness, and (2) generating massive-scale synthetic text-protein pairs from extensive, less-annotated resources—such as UniProtKB/TrEMBL and the AlphaFold Database (AFDB)—by leveraging the retrieval capabilities of ProTrek. This strategy yields a final dataset of over 1.7 billion text-protein pairs, the composition of which is detailed in the following subsections and summarized in Supplementary Table 2.

#### 7.3.1 SwissProt-Annot

Following the procedure in ProTrek ^42^, we construct the SwissProt-Annot dataset by extracting protein function information from 56 distinct subsections, including protein-level (*e.g*. Function, Miscellaneous, Caution) and residue-level (*e.g*. Active site, Binding site, DNA binding), from Swiss-Prot database (Release 2023 03). We employ GPT-4 to enhance the linguistic diversity of our training data, using distinct strategies tailored to the format of the Swiss-Prot annotations. For each protein-level description, we generate 10 paraphrases. For the residue-level annotations, we prompt GPT-4 to generate a diverse pool of sentence templates, which are then populated with specific annotations to form a variety of complete sentences (see Supplementary Table 5 for the generation prompt and Supplementary Table 4 for an illustrative example). To provide the model with natural language descriptions of enzymatic function, rather than just symbolic chemical equations from Swiss-Prot, we incorporated the comprehensive textual annotations for each EC number from the ENZYME database ^60^. Specifically, we download data from the official website (https://ftp.expasy.org/databases/enzyme/enzyme.dat) and extract the descriptions and comments (*i.e*., the “DE” and “CC” lines). This process results in an additional 253,279 protein-text pair data for SwissProt-Annot.

#### 7.3.2 SwissProt-Aug

While the SwissProt-Annot dataset is highly structured, it typically presents each piece of functional information as isolated sentences. To create more comprehensive and coherent functional descriptions, we integrate these individual sentences into unified narratives. We first establish 56 canonical questions, each corresponding to one of the 56 subsections of a Swiss-Prot entry. To ensure linguistic diversity, we then use GPT-4 to generate 100 distinct paraphrases for each canonical question. Subsequently, we prompt the *glm-4-flash* model ^61^ with a prompt constructed from two components: a domain-specific question (selected from the paraphrases) and the collection of existing descriptions for a given protein from SwissProt-Annot. The language model synthesizes the provided descriptions to generate a comprehensive answer, which serves as the new functional description for the protein. The complete prompts used in this process are detailed in Supplementary Table 5.

#### 7.3.3 TrEMBL50-ProTrek

To address the scale limitations of the high-quality Swiss-Prot database, we generate a massive-scale dataset by pairing proteins in TrEMBL with descriptions from SwissProt-Annot. Specifically, we first cluster the TrEMBL database at 50% sequence identity, yielding a set of representative proteins (hereafter referred to as TrEMBL50). Next, we construct a comprehensive text pool by aggregating all functional descriptions from SwissProt-Annot. Finally, for each protein in TrEMBL50, we employed the ProTrek model to calculate the protein-text matching scores. By selecting the top 10 descriptions for each protein, this process yielded the TrEMBL50-ProTrek dataset, containing approximately 412 million synthetic protein-text pairs. Additionally, we made targeted additions for protein classes with limited functional annotations in this training set. We curated 46 protein classes (Supplementary Table 1) and used their names as textual queries for ProTrek-based retrieval, primarily from TrEMBL50 with source details provided in Supplementary Table 1, generating approximately 22,000 additional synthetic protein-text pairs.

#### 7.3.4 SwissProt-LLM and TrEMBL50-ProLLM

We incorporated two datasets introduced by Evolla ^62^, which contain LLM-generated question-answer pairs for proteins from Swiss-Prot and TrEMBL50, respectively. We extracted only the answers from each pair to serve as functional descriptions, creating two datasets: SwissProt-LLM and TrEMBL50-ProLLM.

#### 7.3.5 Keyword-protein pairs from InterPro database

To further expand the diversity of textual prompts, we incorporate keyword-protein pairs from the InterPro database (version 101.0; https://www.ebi.ac.uk/interpro/download/InterPro/). We collect these pairs from InterProScan annotations, using the functional keywords directly as a concise form of textual input. This dataset ensures the model is exposed to a wide range of short, function-centric descriptions.

#### 7.3.6 Unannotated structures from the AlphaFold database

To enhance the structural diversity of our training set, we include a large corpus of unannotated proteins from the AlphaFold Database (AFDB). As these proteins lack descriptive text, we pair them with a randomly selected prompt from a predefined set of 50 non-informative sentences (*e.g*. “design a protein randomly”). This strategy of using a various set of generic prompts encourages the model to explore the vastness of the protein fold space independently of any specific functional constraints.

### 7.4 Limitations of data synthesis

While our ProTrek-based augmentation strategy effectively expands the sequence and structural diversity for functions represented in Swiss-Prot, we note an inherent limitation: functions that are entirely undocumented in curated databases cannot be explicitly modeled. Because TrEMBL annotations are often computationally inferred, sparse, or redundant, anchoring to Swiss-Prot is necessary to reduce noise in text supervision. Consequently, our approach is designed to broaden the coverage of known functions rather than to discover entirely new functional categories.

Furthermore, the use of ProTrek for large-scale retrieval inevitably introduces a degree of noise, as its matching scores serve as empirical signals rather than perfectly calibrated estimates of semantic correctness. To mitigate the impact of inaccurate matches, we employed conservative quality-control thresholds to retain only high-confidence pairs. Although some residual noise remains, our empirical scaling results demonstrate that the model effectively tolerates this noise at scale, extracting a net positive training signal from the massive data expansion.

### 7.5 Training regimen

We implemented the training using PyTorch. To optimize GPU memory usage, we employed activation checkpointing ^63^ and applied DeepSpeed ZeRO-2 ^64^ for gradient partitioning and optimizer partitioning. All models were trained using BFloat16 mixed-precision to enhance efficiency.

The models were trained with the standard AdamW optimizer, using hyperparameters of *β*_1_ = 0.9, *β*_2_ = 0.98 and *ɛ* = 1e-8. We used a global batch size of 512 and applied a weight decay of 0.01 to prevent overfitting. The learning rate was warmed up over the first 10% of training steps, followed by a cosine decay schedule. Additional details on other hyperparameters are provided in Supplementary Table 3.

To ensure a balanced representation of protein space and mitigate distributional biases, our training regimen leverages the AFDB cluster ^65^ for sampling proteins from the InterPro and AFDB datasets. This is implemented through a hierarchical, two-stage sampling strategy: a cluster is first selected uniformly at random, and a protein is then sampled uniformly from within the chosen cluster. For the remaining datasets, proteins are sampled directly, without this cluster-based approach. Furthermore, each dataset is assigned a specific sampling weight, as detailed in Supplementary Table 2.

#### 7.5.1 SaProt-T/O training details and rationale

To generate amino acid sequences in the second stage of the Pinal pipeline, we develop two specialized models, SaProt-T and SaProt-O, based on the SaProt architecture.

SaProt-T(ext) is specifically designed for the task of protein reverse folding: predicting amino acid sequences given both protein backbone structures (as 3Di tokens) and textual descriptions.

SaProt-O(mni) is trained for maximum flexibility to better align with diverse, real-world protein (re)design workflows. It is trained on various combinations of inputs, which may include or exclude textual prompts, structural information, or portions of the amino acid sequence (see Extended Data Fig. 2b for data combination details). This “omni-modal” training strategy enables SaProt-O to effectively handle a wide range of design scenarios, such as infilling, redesign, or sequence generation from structure and text alone. To validate our core hypothesis—that conditioning on text is crucial for generating high-quality, functionally relevant sequences—we conducted an ablation study comparing our models against the original SaProt and the ProstT5 ^66^ baseline. While existing models like baseline SaProt ^36^ excel at predicting sequences from structural tokens alone, we posit that text provides essential fine-grained cues not present in the coarse-grained 3Di tokens. To test this hypothesis, we compared our text-conditioned models SaProt-T and SaProt-O (760M) with the original SaProt (650M) and a variant, SaProt-T-T5xxl encoder (the same text encoder used in the 15.5B T2struct), to assess the impact of text encoder capacity.

The findings (Extended Data Fig. 2a) demonstrate that sequences generated by SaProt-T and SaProt-O achieve stronger textual alignment while maintaining higher predicted foldability than those generated by the baseline models. In contrast, although ProstT5 shows good textual alignment, its generated sequences have lower predicted foldability than those from our models. Furthermore, simply scaling the text component, for example by using the pre-trained T5 encoder in SaProt-T-T5xxl encoder, did not improve the performance compared to SaProt-T. This result underscores the significance of our text-conditioning strategy and justifies the development of SaProt-T/O for high-fidelity, language-guided sequence design.

In this paper, we use SaProt-T for all Pinal experiments. By contrast, SaProt-O is more suited for protein editing tasks where partial sequences or structures are provided. For example, after proteins are designed by Pinal, one can use SaProt-O to re-engineer the protein by mutating multiple amino acids. We leave the systematic exploration of SaProt-O applications as future work.

#### 7.5.2 The stability issue of large-scale model training

Training the 15.5B T2struct model presented a challenge due to its size. We observed that both training and validation losses exhibited spikes and failed to converge after the learning rate warmup phase. Inspired by prior work ^67^ suggesting that learning rates should be adapted based on model parameters, we reduced the learning rate to 1e-5, which is 10 times smaller than the initial value. This adjustment effectively resolved the training stability issues, enabling the model to converge reliably and maintain a consistent loss trajectory throughout training.

#### 7.5.3 Training phase for T2struct

The training protocol for the 15.5B T2struct model was executed in three sequential phases. The initial phase consisted of 320,000 training steps on the complete data corpus, utilizing an experimental sampling distribution specified in Supplementary Table 2. A key feature of this stage was the stratified sampling of the TrEMBL-ProTrek dataset, wherein pairs with a ProTrek score of 12 or higher were sampled with 80% probability, versus 20% for pairs with scores below 12. This was followed by a 305,000-step training phase on a curated corpus. For this second phase, the TrEMBL-ProTrek data component was wholly substituted with a high-quality subset, which was derived by filtering for pairs with a ProTrek score exceeding 12, thereby reducing its volume from 412 million to approximately 130 million pairs. The sampling weights for all datasets held constant from the initial phase. While the two phases training have already established Pinal’s superior performance over baselines like ESM3, Chroma, and ProteinDT, we identified a tendency in all models to generate repetitive amino acid sequences (e.g., “AAAAAAA”) ^68^. Our investigation revealed that the root cause was not the sequence design module (SaProt-T), but rather the generation of repetitive Foldseek structural tokens—a tendency correlated with training data derived from lower-quality predicted structures in the AFDB (Extended Data Fig. 5b).

Based on these findings, we introduce a third training phase to specifically address this repetition issue. This phase involves fine-tuning the model for an additional 115,000 steps on a further refined dataset, from which we exclude protein structures with a pLDDT score below 70. The effectiveness of this refined training strategy is confirmed by evaluations on the hard test set (Extended Data Fig. 5a). Specifically, this third phase improves the model’s performance on foldability metrics (pLDDT and PAE). Furthermore, as shown by a comparative analysis of the model before and after this phase (Phase 2 *vs*. Phase 3), training with this higher-quality structural data substantially decreases the percentage of designed proteins containing excessive amino acid repeats (Extended Data Fig. 5b), validating our approach.

The total training across all phases cost approximately 50,000 H800 Nvidia GPU hours.

### 7.6 Initial candidate generation and screening

Protein generation with Pinal follows a two-stage procedure. First, T2struct generates structure-token sequences from the input natural-language description using multinomial sampling, enabling diverse structural candidates. Second, SaProt-T decodes each structure-token sequence into an amino-acid sequence conditioned on both the input description and the generated structure tokens.

For large-scale in silico evaluations, we used deterministic SaProt-T decoding to obtain one reproducible, high-probability amino-acid sequence for each generated structure-token sequence. Specifically, SaProt-T was decoded by argmax sampling, selecting the amino acid with the highest predicted probability at each position. We sampled *K* = 50 structure-token sequences for each text prompt, yielding 50 initial candidates per text prompt. These candidates were ranked by their joint log-probability of the generated structure-token and amino-acid sequences under Pinal, normalized by sequence length, and the top five candidates were retained for downstream computational evaluation.

To determine an appropriate value of *K*, we conducted an empirical study on the hard test set. We varied *K* from 5 to 100 and found that performance, measured by the ProTrek score and pLDDT of the top-ranked designs, improved notably up to *K* = 50, after which the gains became marginal (Extended Data Fig. 3). We therefore used *K* = 50 for computational evaluation, balancing performance and computational cost.

For wet-lab validation, we used the same two-stage procedure but larger task-specific sampling scales, because each experiment focused on a single target protein. We generated 50,000 structure-token sequences for GFP and H-protein, 2,500 for ADH, and 200,000 for PETase. The SaProt-T decoding strategy, multinomial sampling settings, and initial candidate-pool sizes are summarized in Supplementary Table 13.

Inference is performed on NVIDIA H800 GPUs, which require at least 40GB of memory to run the 15.5B T2struct model. On a single GPU, Pinal generates sequences at a rate of approximately 300 residues per second. The model’s maximum generation length is 1022 residues, with inference time increasing with sequence length.

### 7.7 Benchmarking and evaluation protocol

#### 7.7.1 Baseline setup

The baseline models evaluated in this study were run using their official implementations and publicly released pretrained checkpoints. For ProteinDT, we followed the official text-to-protein pipeline using the pretrained model, Gaussian facilitator and T5-based autoregressive protein decoder. Sequences were generated by stochastic decoding with top-*k* = 40, top-*p* = 0.9 and a temperature of 1.0. For Chroma, we used ProCap conditioning with the guidance weight of 10, generated protein backbones using 50 diffusion steps with a reverse-time SDE and Euler–Maruyama integration, and performed Potts-based sequence co-design using the official default settings. For ESM3, we evaluated the open 1.4-billion-parameter `biohub/esm3-sm-open-v1` checkpoint, setting the number of generation steps to the target sequence length and sampling sequences at a temperature of 0.7.

#### 7.7.2 Construction of evaluation sets

Our evaluation protocol assesses two distinct capabilities of Pinal: generalization from complex functional descriptions and translation of specific functional prompts into proteins.

##### Designing protein from complex text descriptions (long sentences evaluation)

To evaluate the generalization capability of Pinal on unseen text-protein pairs, we constructed two distinct evaluation sets from a held-out portion of the Swiss-Prot dataset. First, we established a standard test set comprising 3,304 proteins. This set was achieved by clustering the Swiss-Prot database at a 50% sequence identity threshold and selecting protein clusters that were disjoint from those used in training. For each protein in this set, we generated comprehensive descriptions by simply concatenating all available functional annotations. These descriptions were subsequently paraphrased using GPT-4 to ensure the resulting prompts were textually novel to our model. From this pool of unique prompts, we randomly sampled 500 for the long-sentence-based design evaluation. Second, we created a more stringent ‘hard’ test set containing 465 proteins. These proteins were derived by filtering the standard test set, retaining only proteins that share less than 30% sequence identity with any protein in the training clusters. We applied the prompt generation procedures used in both the standard test set and SwissProt-Aug, and randomly sampled another 500 prompts to constitute the final ‘hard’ evaluation set. For each description, we then generated five protein candidates, resulting in a total of 2,500 designed proteins per setting for our analysis.

##### Translating specific functional prompts (short sentences & keyword-based evaluation)

To assess the model’s ability to accurately translate functional concepts, we created two evaluation sets by sampling prompts directly from the entire datasets. The short sentence set was constructed to cover a broad range of functional commands while simultaneously testing the model’s robustness to textual variation. To achieve this, we first systematically sampled 50 descriptions from each of the 17 distinct protein-level subsections within SwissProt-Annot. These 850 descriptions were then processed using the same paraphrasing protocol as the long sentences to ensure their phrasing was novel to the model. The keyword-based set was designed for a direct, head-to-head comparison with ESM3. This set was constructed by randomly sampling 500 entry names from the InterPro database to serve as textual inputs for both models, ensuring a fair benchmark of their keyword-conditioned generative capabilities. Similar to the long-sentence evaluation, we then generated five protein candidates for each prompt in these sets.

#### 7.7.3 Prompt generation for structurally uncharacterized protein

The construction of these prompts follows a multi-step pipeline. First, we identify a pool of structurally uncharacterized proteins by comparing AlphaFold-predicted structures against the entire PDB database using Foldseek. We retain 6,000 proteins whose predicted structures show no substantial structural similarity (TM-score *<* 0.5) to any known experimental structure. For this set of novel proteins, we first use ProTrek to predict functional text for them, and then we use a large language model ^61^ to paraphrase this retrieved text to ensure linguistic diversity. These rewritten descriptions serve as the final input prompts for the novelty generation task.

#### 7.7.4 Metrics

Evaluation metrics are grouped into three primary categories: textual and structural alignment, foldability, and novelty.

##### Textual and structural alignment

We use three metrics to assess alignment with the input prompt. For designs with a known native structure, we report the GT-TMscore, defined as the TM-score (https://zhanggroup.org/TM-score/) between the design’s predicted structure and the ground-truth structure. For keyword-based prompts, we additionally report keyword recovery, calculated with InterProScan (https://www.ebi.ac.uk/interpro/download/InterProScan/) to measure functional consistency, strictly following the protocol of ESM3 ^38^. For a more general assessment, we use the ProTrek score ^42^, which measures the cosine similarity between the prompt’s text embedding and the embeddings of the designed sequence (Seq. ProTrek) or structure (Struc. ProTrek).

For both sequence and structure modalities, ProTrek scores computed using native protein-text pairs are highly correlated with those computed using semantically equivalent paraphrased descriptions, as measured by Spearman correlation. This indicates that the metric is robust to variation in wording. Furthermore, the scores of correctly matched pairs are higher than those of randomly shuffled mismatched pairs, indicating that the method has a strong discriminative ability (see Extended Data Fig. 4 for full analysis). For evaluations in which multiple designs were considered for the same text prompt (i.e., Fig. 2b,c,d,e,f,g), Pinal first generated multiple candidates (*K* = 50) and then retained the top five designs per prompt. Metrics were computed for each of these five retained designs. Unless otherwise specified, reported values correspond to the mean across the retained designs. When both mean and maximum ProTrek values are shown, Mean Seq. and Max. Seq. denote the mean and maximum sequence-to-text ProTrek scores across the five retained designs, respectively, whereas Mean Struc. and Max. Struc. denote the corresponding mean and maximum structure-to-text ProTrek scores.

##### Foldability

The structural plausibility of each designed sequence is predicted with ESMFold ^69^ or AlphaFold3 ^70^. We report two standard confidence metrics: the predicted Local Distance Difference Test (pLDDT), a per-residue confidence score, and the Predicted Aligned Error (PAE), which measures the confidence in relative domain positions. Higher pLDDT and lower PAE values indicate a more confident, well-folded structure.

##### Novelty

The novelty of a design is determined by evaluating both sequence and structural homology. Sequence novelty is assessed by sequence identity against the UniRef100 database, while structural novelty is determined by comparing it against all entries in the Protein Data Bank (PDB), reporting the highest query TM-score as the PDB-TM.

##### Diversity

To quantify the generative diversity, we clustered the generated proteins in both sequence and structural spaces following the DISCO methodology ^71^, tracking the total number of distinct clusters.

##### Evaluation limitations and discussion

Pinal-generated proteins were evaluated using complementary metrics capturing prompt alignment, structural plausibility and annotation-level functional consistency. Prompt-sequence semantic alignment was quantified using the ProTrek score, defined by the similarity between the input text representation and the generated protein representation in the ProTrek embedding space. Because ProTrek was also used during synthetic data construction, this metric is not fully independent of the training-data generation procedure. We therefore complemented ProTrek-based evaluation with additional alignment metrics. Structural quality was assessed using TM-score, which compares predicted structures independently of ProTrek. Functional consistency was further evaluated by keyword recovery, in which annotations inferred from generated sequences using InterProScan were compared with the keywords specified in the input prompts. Because text-guided protein generation is an emerging task and no universally accepted metric exists for evaluating prompt-level functional consistency, we report these complementary metrics together rather than relying on any single score.

### 7.8 Phylogenetic analysis of Pinal-generated sequence diversity

To evaluate the sequence diversity of Pinal-designed proteins, we retrieved natural homologous sequences for each target protein family from the Swiss-Prot database. For each family, 5,000 sequences were generated using Pinal and filtered to retain high-confidence designs with a ProTrek score *>* 15. These designs also exhibited high predicted structural confidence (average ESMFold pLDDT *>* 85, Supplementary Fig. 4). To reduce redundancy, the natural and filtered Pinal-designed datasets were independently clustered using CD-HIT in a two-step procedure, first at 90% sequence identity (-c 0.9, -aS 0.8) and then at 70% sequence identity (-c 0.7, -aS 0.8). Representative sequences from the natural and designed sets were combined and aligned using Clustal Omega. Maximum-likelihood trees were inferred from the amino acid sequence alignments using IQ-TREE 2 with ModelFinder-based automatic model selection (-m MFP). Branch support was estimated using 1,000 ultrafast bootstrap replicates (-bb 1000), and the number of CPU threads was selected automatically (-nt AUTO). The resulting trees were midpoint-rooted for visualization and used to compare the distribution of high-confidence Pinal-designed sequences with that of curated natural homologues. Midpoint rooting was used only for visualization.

### 7.9 Materials, strains, and cultivation

Glycine, nicotinamide adenine dinucleotide (NAD^+^, NADH), and tetrahydrofolic acid (THF) were obtained from Sigma-Aldrich (St. Louis, MO, USA). Dithiothreitol (DTT), pyridoxal 5’-phosphate monohydrate (PLP), Ethanol, Acetaldehyde, 2,4-Dinitrophenylhydrazine (DNPH), Acetonitrile (ACN), Trifluoroacetic acid (TFA), Imidazole, NaOH, HCl, KH_2_PO_4_, 1,1,1,3,3,3-Hexafluoro-2-propanol (HFIP), 4-nitrophenyl butyrate (pNPB), standards of bis-hydroxyethyl terephthalate (BHET), terephthalic acid (TPA) and mono(2-hydroxyethyl) terephthalate (MHET) were obtained from Aladdin (Shanghai, China). *E. coli* BL21(DE3) (Tsingke Biotechnology, Beijing, China) was used as the primary host for recombinant protein expression. Cultures were routinely grown in Luria-Bertani (LB) medium at 37 ^◦^C with shaking at 200–225 rpm.

### 7.10 Recombinant protein expression and purification

#### 7.10.1 Soluble-expression analysis of structurally novel designs

Twenty computationally selected structurally novel protein designs were tested for soluble expression in *E. coli* BL21(DE3). Recombinant proteins carrying an N-terminal SUMO tag and Strep-tag II were expressed from three independent colonies for each design. Cultures were grown to an OD_600_ of approximately 0.6 and induced with 0.1 mM IPTG at either 22 ^◦^C or 37 ^◦^C for 20 h. Cells were harvested and lysed in PBS, and whole-cell lysates and soluble fractions were analysed by SDS–PAGE (Supplementary Fig. 5). Uncropped and unprocessed SDS–PAGE images underlying Supplementary Fig. 5 are provided in the Source Data file.

#### 7.10.2 Expression and purification of pinalGFP

The codon-optimized gene sequence for pinalGFP, along with benchmark proteins, was synthesized and cloned into a pET28a(+) vector containing an N-terminal 6×His tag (GENEWIZ, Suzhou, China). The construct was transformed into *E. coli* BL21(DE3) cells (Tsingke, Beijing, China). For large-scale protein production, transformed BL21(DE3) cells were cultured in 4 L of LB medium at 37 ^◦^C until the OD_600_ reached 0.6–0.8. Protein expression was induced with 0.5 mM IPTG, followed by incubation at 18 ^◦^C for 16– 20 h. Cells were harvested by centrifugation, resuspended in lysis buffer (50 mM Tris-HCl, 500 mM NaCl, 10 mM imidazole, pH 8.0), and lysed by sonication on ice. The cleared lysate was obtained by centrifugation (4,000 × g, 30 min, 4 ^◦^C). The supernatant was first purified using Ni-NTA agarose affinity chromatography (Qiagen). The bound protein was eluted with a buffer containing 250 mM imidazole. Following concentration using Amicon Ultra-15 centrifugal filters (30 kDa MWCO, Millipore), the protein was further purified by size exclusion chromatography on a Superdex 200 pg 16/600 column using an Ä KTA prime PLUS system (GE Healthcare), equilibrated with 25 mM Tris-HCl, 150 mM NaCl, pH 8.0.

#### 7.10.3 Expression and purification of pinalADH and pinalPETase

Candidate gene sequences for pinalADH and pinalPETase, codon-optimized for *Escherichia coli*, were commercially synthesized (Guangzhou Aiji Biotechnology Co., Ltd.). The synthesized genes were cloned into the pET28a(+) expression vector containing a C-terminal 6×His tag and transformed into *E. coli* BL21(DE3) competent cells. For protein expression, transformed cells were cultured in 100 mL of LB medium at 37 ^◦^C with shaking at 225 rpm until the OD_600_ reached 0.6. Expression was induced with a final concentration of 0.1 mM IPTG, and the cultures were subsequently incubated at 16 ^◦^C and 120 rpm for 24 h. Cells were harvested by centrifugation (4,500 × g, 30 min), and the pellet was washed twice with 10 mM phosphate buffer. Cell disruption was performed using a high-pressure homogenizer (KS-2000M, Kaibai Nano Technology). The lysate was cleared by centrifugation at 12,000 × g for 30 min at 4 ^◦^C. The clarified supernatant was loaded onto a 5 mL HISTRAP FF CRUDE column (Cytiva) using an Ä KTA pure 25 system (Cytiva). The column was washed with 50 mM phosphate buffer (pH 7.8) containing 60 mM imidazole and 500 mM NaCl. The protein was then eluted with 50 mM phosphate buffer (pH 7.8) containing 250 mM imidazole and 500 mM NaCl. Protein concentration was determined using a BCA Protein Assay Kit (Biogene), and purity was assessed by SDS-PAGE.

#### 7.10.4 Expression and purification of pinalH

The selected H-protein genes, codon-optimized for *E. coli*, were synthesized with C-terminal 6×His-tags and cloned into the pET28a vector (Tsingke Biotechnology). The constructs were transformed into *E. coli* BL21(DE3) for expression. Transformants were cultured in LB medium with 100 mg L^−1^ ampicillin at 37 ^◦^C until an OD_600_ of 0.6–0.8 was reached. Expression was induced with 0.1 mM IPTG, and cultures were incubated at 16 ^◦^C for 16 h. Cells were harvested by centrifugation (8,000 rpm, 10 min, 4 ^◦^C) and washed with 50 mM phosphate buffer (pH 7.0). For purification, cell pellets were resuspended in 50 mM Tris buffer (pH 7.0) and lysed by ultrasonication. The lysate was cleared by centrifugation (10,000 × g, 5 min, 4 ^◦^C). The supernatant was applied to a gravity-flow column packed with Ni^2+^-charged resin (High Affinity Ni-charged Resin FF, GenScript) pre-equilibrated with lysis buffer (50 mM Tris, 10 mM imidazole, 300 mM NaCl, pH 7.5). The column was washed with a wash buffer (50 mM Tris, 30 mM imidazole, 300 mM NaCl), and the purified H-proteins were eluted using an elution buffer (50 mM Tris, 300 mM imidazole, 300 mM NaCl, pH 7.5). Excessive imidazole was removed by ultracentrifugation. Purity was confirmed by SDS-PAGE.

### 7.11 Characterization of pinalGFP

#### 7.11.1 Computational screening and design refinement of pinalGFP

An initial pool of 50,000 GFP candidate sequences generated by Pinal was processed through a multi-tiered computational pipeline. Quality filtering retained 16,843 sequences with a ProTrek score *>* 15 and high AlphaFold3-predicted structural quality (pLDDT *>* 80, PAE *<* 7). The remaining candidates were then filtered by similarity to known fluorescent proteins, retaining sequences with *<*35% sequence identity. To identify candidates with autonomous chromophore-forming potential, we further applied a sequence-based motif screen around the expected GFP chromophore-forming region, using the canonical Ser65–Tyr66–Gly67 site in avGFP ^72^ (PDB: 2WUR) as the reference. Candidates were retained if this region contained a known chromophore-forming triplet, specifically SYG, TYG, SHG, TWG, QYG, or HYG, based on motifs observed in characterized fluorescent proteins ^73^. Together, the sequence-identity and motif filters yielded 29 diverse candidates for experimental validation. We note that this conservative motif filter considered only six chromophore-forming triplets, a limited subset of the broader repertoire reported in characterized fluorescent proteins. It was therefore used for experimental prioritization rather than as an exhaustive assessment of fluorescence potential, and sequences excluded by this step were not interpreted as non-functional.

#### 7.11.2 Fluorescence characterization

Fluorescence microscopy images were acquired using an Olympus MVX10 Macro Zoom Fluorescence Microscope System. Fluorescence intensity was quantified on an Agilent Microplate Reader (excitation 460 nm, emission 510 nm). To bring all signals within the linear detection range of the instrument, synthetic variants (pinalGFP, pinalGFP chromophore mutant W73G, and esmGFP-b8) were measured at 1 mg mL^−1^, while avGFP was measured at 1 *µ*g mL^−1^. Fluorescence excitation and emission spectra were recorded using the same instrument from 300–520 nm and 460–650 nm, respectively.

### 7.12 Characterization of pinalPETase

#### 7.12.1 Thermal shift assay by nanoDSF (PR Panta)

Protein thermal stability was measured on a PR Panta instrument (NanoTemper Technologies) using nano-differential scanning fluorimetry (nanoDSF) in chemically inert glass capillaries (NanoTemper, Cat. #K003). Each sample (10 µL, 5–250 µg mL^−1^ in the final formulation buffer) was loaded by capillary action into the standard chip. The temperature was ramped from 25 ^◦^C to 95 ^◦^C at a constant rate of 5 ^◦^C min^−1^, while intrinsic fluorescence was recorded at 330 nm and 350 nm every 0.2 ^◦^C (≥20 data points min^−1^). Melting temperature (*T*_m_) was derived from the first derivative of the ratio *F*_350 nm_*/F*_330 nm_ versus temperature.

#### 7.12.2 Esterase activity test

The enzymatic activity of the expressed proteins was measured using p-nitrophenyl butyrate (pNPB) as the substrate. *E. coli* BL21(DE3) cells were cultured in LB medium supplemented with kanamycin (50 *µ*g mL^−1^). Protein expression was induced with 0.2 mM IPTG, and the cells were incubated at 30 ^◦^C and 200 rpm for 36 h. Following incubation, cells were harvested by centrifugation at 4000 rpm for 30 min. The cell pellets were resuspended in 400 *µ*L Bugbuster Protein Extraction Reagent and incubated at 30 ^◦^C with shaking (200 rpm) for 40 min. After lysis, the samples were centrifuged at 4000 rpm for 30 min, and the supernatants (90 *µ*L) were mixed with 10 *µ*L of pNPB substrate. The reaction mixture was incubated at 37 ^◦^C in the dark for 10 min. The reaction was terminated by adding 100 *µ*L of anhydrous ethanol, and the absorbance at 410 nm was measured immediately. Total protein concentration in the supernatant was determined using the Bradford assay. pNP concentration was determined from a pNP standard curve (Extended Data Fig. 8b). The resulting pNP concentration was normalized to the total protein amount in the 90 *µ*L soluble fraction and expressed as M g^−1^ total protein.

#### 7.12.3 Plastic hydrolysis activity test

Amorphous PET film (Goodfellow, ES30-FM-000145; thickness, 0.25 mm) was cut into small pieces and completely dissolved to form a homogeneous polymer solution. Pretreated PET films were prepared following a previously reported PET membrane preparation method ^74^. The resulting PET films were collected, washed, dried, and used as the pretreated PET substrate in subsequent experiments. Plastic hydrolysis activity was assessed by incubating 1 mg of pretreated PET substrate with 1 *µ*M enzyme in 600 *µ*L of 100 mM KH_2_PO_4_–NaOH buffer (pH 8.0) at 40 ^◦^C. Samples were collected on days 1, 3, 5 and 7 for HPLC analysis. Soluble degradation products (MHET and TPA) were quantified by HPLC using a Shim-pack GIST C18-AQ column (4.6 × 150 mm, 3 *µ*m; Shimadzu) maintained at 25 ^◦^C. Mobile phase A consisted of water containing 0.1% (v/v) TFA, and mobile phase B consisted of acetonitrile containing 0.1% (v/v) TFA. Aliquots (10 *µ*L) were injected for analysis. Separation was performed at a flow rate of 0.8 mL min^−1^ using the following gradient: 5% B from 0 to 3 min, 5–20% B from 3 to 6 min, 20–25% B from 6 to 11 min, 25–95% B from 11 to 19 min, 95% B from 19 to 23 min, and 95–5% B from 23 to 25 min, followed by re-equilibration at 5% B. The total run time was 32 min, and detection was performed at 260 nm.

#### 7.12.4 Scanning electron microscopy

PET films were mounted on SEM stubs using conductive adhesive tape and sputter-coated with gold prior to imaging. The surface morphology was examined using a field-emission scanning electron microscope (FE-SEM; Sigma 300, Zeiss). Images were acquired using secondary-electron detection at an accelerating voltage of 10–15 kV and at magnifications of 2,000× and 10,000×.

### 7.13 Characterization of pinalADH

#### 7.13.1 Preliminary screening of pinalADH

Pinal generated an initial library of 2,500 ADH candidates from five paraphrased general ADH-design prompts, with 500 candidates sampled from each prompt. After removing redundant sequences and applying computational quality filters (ESMFold pLDDT *>* 80 and PAE *<* 7), 1,639 non-redundant candidates were retained. From this set, 1,000 sequences were selected for downstream consideration. To promote coverage of diverse ADH-like architectures, the retained sequences were grouped by protein length into three approximate ranges centered around 250, 350 and 380 amino acids, broadly corresponding to short-chain, medium-chain and longer ADH-like designs, respectively. For experimental validation, eight candidates were selected from these length-defined groups, including four short-chain, two medium-chain and two long-chain ADH designs, with ESMFold pLDDT considered during prioritization.

#### 7.13.2 Derivatization and HPLC analysis of acetaldehyde

To detect acetaldehyde formation from ethanol, we employed a 2,4-dinitrophenylhydrazine (DNPH) derivatization strategy. Briefly, 100 *µ*L reaction mixtures were quenched with 100 *µ*L of DNPH solution (1 mg mL^−1^ in acetonitrile) and 4 *µ*L of phosphoric acid solution (85% phosphoric acid/water, 1:4, v/v). Samples were incubated at room temperature for 20 min in the dark to allow formation of the corresponding hydrazone derivative. Following centrifugation at 12,000 r.p.m. for 10 min, the supernatants were collected and filtered through 0.22 *µ*m membranes. Aliquots (5 *µ*L) were injected onto an HPLC system equipped with a Shim-pack GIST C18-AQ column (4.6 × 150 mm, 3 *µ*m; Shimadzu) maintained at 40 ^◦^C. Mobile phase A consisted of water containing 0.1% (v/v) TFA, and mobile phase B consisted of acetonitrile containing 0.1% (v/v) TFA. Separation was performed at a flow rate of 1.0 mL min^−1^ using the following gradient: 50% B from 0 to 3.5 min, 50–98% B from 3.5 to 5.5 min, 98% B from 5.5 to 10.0 min, 98–50% B from 10.0 to 10.5 min, and 50% B from 10.5 to 15.0 min. Detection was performed at 360 nm.

#### 7.13.3 PinalADH activity test

The activity of designed ADHs was determined by detecting the production of acetaldehyde from ethanol. The 100 *µ*L reaction mixture contained 100 mM ethanol, 500 *µ*M NAD^+^, 0.5 mg mL^−1^ purified enzyme, and 20 mM Tris-HCl buffer (pH 8.0). Reactions were initiated by addition of the purified enzyme and incubated at 37 ^◦^C for 120 min. The resulting acetaldehyde was derivatized and analysed by HPLC as described above (Methods 7.13.2), with the acetaldehyde–DNPH derivative eluting at approximately 7.3 min (Extended Data Fig. 9c).

### 7.14 Characterization of pinalH

#### 7.14.1 Reconstituted Glycine Cleavage System assay

The function of designed H-proteins was assessed in a reconstituted GCS. The reaction mixture (50 mM Tris-HCl, pH 7.5) contained 200 mM NH_4_HCO_3_, 60 mM DTT, 1 mM THF, 30 mM formaldehyde, 5 *µ*M P-protein, 3 *µ*M T-protein, 1 *µ*M LplA, 0.5 mM lipoic acid, 1 mM ATP, 1 mM MgCl_2_, and was initiated by adding 40 *µ*M of the respective designed H-protein. The reaction was carried out at 42 ^◦^C for 12 h, with samples taken periodically for analysis.

#### 7.14.2 Quantification of glycine

Glycine concentration was determined by HPLC following pre-column derivatization with dansyl chloride. 40 *µ*L of the reaction mixture was mixed with NaHCO_3_ and dansyl chloride solution, incubated at 30 ^◦^C for 30 min, and then acidified. The dansyl-glycine derivative was quantified by HPLC on a Shim-pack GIST C18 column, with detection at 254 nm. Results were cross-verified using an alternative OPA derivatization method on an Agilent InfinityLab Poroshell HPH-C18 column.

## 8 Data availability

Protein sequences were obtained from UniProtKB/Swiss-Prot and UniProtKB/TrEMBL (https://www.uniprot.org/, release 2023 03). Predicted protein structures were obtained from the AlphaFold Protein Structure Database (https://alphafold.ebi.ac.uk/). Functional annotations and keywords were obtained from InterPro (version 101.0; https://www.ebi.ac.uk/interpro/), and enzyme classification information was obtained from the ENZYME nomenclature database (https://enzyme.expasy.org/). Experimentally determined protein structures used for structural comparisons were obtained from the Protein Data Bank (PDB; https://www.rcsb.org/). UniRef100 (https://www.uniprot.org/help/uniref) was used for sequence-similarity analyses. Fluorescent-protein reference information was obtained from FPbase (https://www.fpbase.org/). The pre-trained Pinal weights can be downloaded from https://huggingface.co/westlake-repl/Pinal.

## 9 Code availability

Pinal is open-sourced under the Apache 2.0 license. The code repository is available at https://github.com/westlake-repl/Denovo-Pinal. The Pinal web server is located at http://www.denovo-pinal.com/. The SaProt-T/O server, corresponding to the second component of Pinal, is linked from the Pinal web server and can also be accessed directly at http://113.45.254.183:9527/.

## 10 Training data and code availability

The release of large-scale training resources for generative protein design requires careful consideration of potential dual-use biosecurity risks. In developing Pinal, the large-scale structural component of the training corpus was derived from the AlphaFold Database, whose released protein-structure collection does not include viral proteomes. Accordingly, Pinal was not trained or intended for viral protein design. Nevertheless, unrestricted public release of the full pre-training corpus together with the complete training pipeline could lower the barrier for adapting similar systems to sensitive biological datasets. To mitigate this risk, these fullscale training resources were provided only under confidential access for editorial and peer review assessment. Because they will not be publicly released in full, and to support transparency and reproducibility, we instead provide public access to the inference code, evaluation code, model usage instructions, trained model weights, and the web server, in line with common release practices for large-scale protein foundation models such as AlphaFold ^70,75^, ESM3 ^38^ and ESMFold ^69^. Additional information about the composition, sources, filtering strategy, and construction of the training corpus is provided in the Methods and Supplementary Information.

## Acknowledgements

We thank Nan Li and the Westlake HPC Center for providing computing resources and technical support. We also appreciate the valuable suggestions from Anthony Gitter, Haiyan Liu, and Sergey Ovchinnikov, which helped improve the manuscript. We acknowledge the support of the Biosciences Central Research Facility (BioCRF (GZ)) and the Bioresearch Lab at the Hong Kong University of Science and Technology (Guangzhou). Their assistance with equipment and facility access, experimental procedures, and material characterization was essential to this study. Finally, we thank Yongjun Mao for assistance with the H-protein-R2 validation experiments.

## Author contributions

F.Y., F.D., and H.L. conceived the study. F.Y. led the overall project design and computational framework development. H.L., J.L., and T.S. led and supervised the biological experiments. F.D. and S.Y. carried out the main research. F.Y. and F.D. designed the Pinal framework and data-scaling strategy. F.D. implemented and trained the Pinal model. Y.F. and Q.T. performed some model evaluation. Shunzhi Wang, X.C., and Y.W. contributed to discussions and provided insightful feedback. Y.Z., A.Z., and J.L. contributed to the H-protein validation experiments. Y.G., L.F., and T.S. contributed to the GFP validation experiments. X.M., L.Y., S.Y., Z.Z., C.Z., and H.L. contributed to the PETase validation experiments. X.D., S.Y., Y.J., K.H., J.Z., and H.L. contributed to the ADH validation experiments. S.Y., H.Q., Y.K., C.W., Y.W., and L.C. performed the soluble protein expression experiments. Y.H., L.Z., and Sheng Wang contributed to early-stage exploratory experiments. X.Z., J.S., and C.H. collected and curated the datasets used in this study. F.Y., F.D., H.L., S.Y., J.L., and T.S. wrote the paper and all authors read the paper.

---

**Extended Data Fig. 1:**
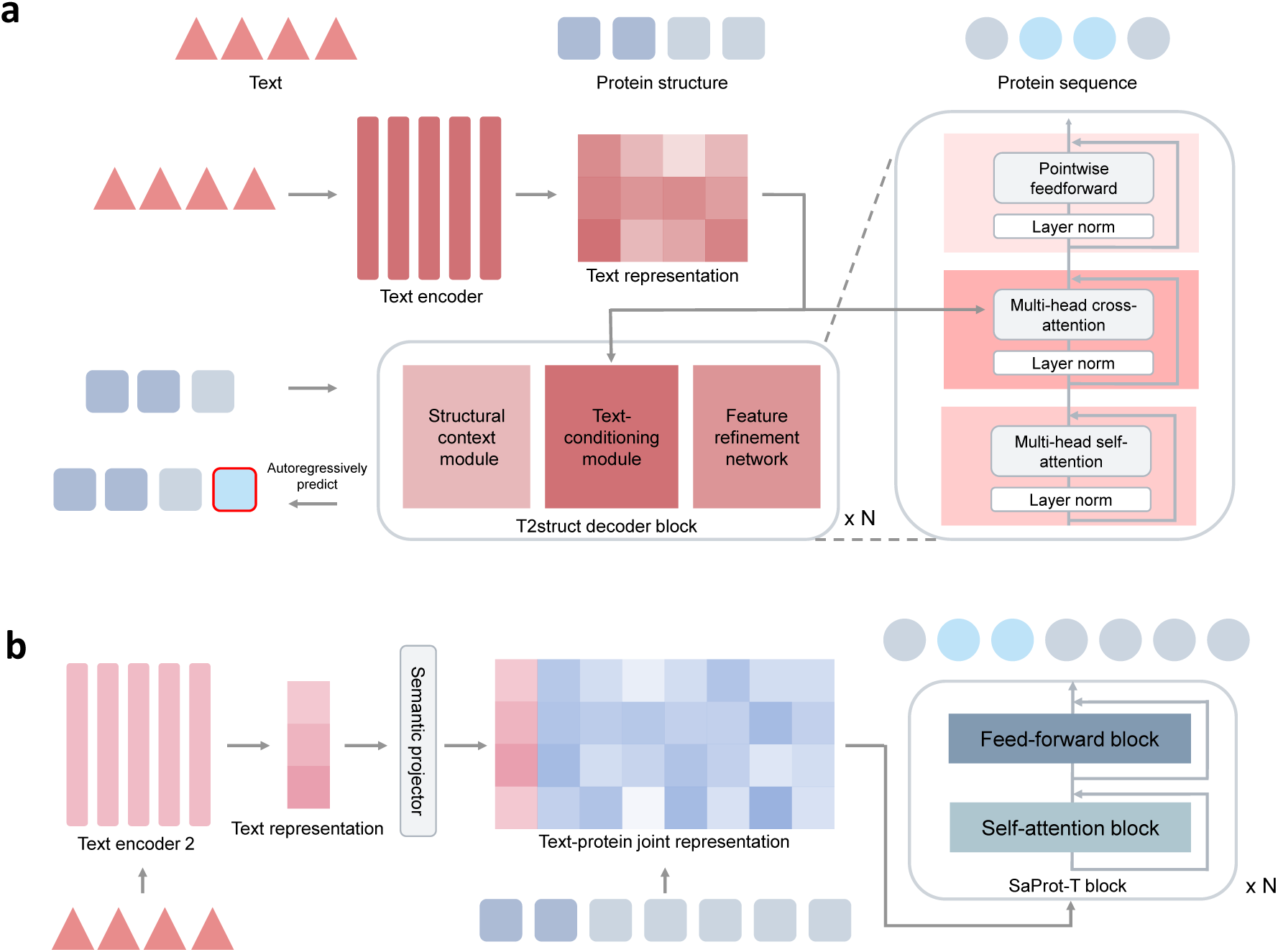
Detailed architectures of the Pinal generative modules. **a:** T2struct translates natural language descriptions into structural (3Di) token sequences. A text encoder first generates the text representation according to the function prompt. This representation is then used to guide the T2struct decoder block, which autoregressively predicts the sequence of 3Di structural tokens. The detailed view (right) illustrates the internal composition of each decoder block (repeated *N* times), consisting of three sequential sub-modules: a structural context module (implemented via multi-head self-attention) to process the generated structural sequence, a text-conditioning module (employing multi-head cross-attention) to integrate guidance from the text representation, and a feature refinement network (a pointwise feed-forward layer) for further feature processing. Layer normalization is applied within each sub-module. **b:** The SaProt-T module, which designs an amino acid sequence based on both the structural input and the text prompt. A semantic projector maps the text representation into the same embedding space as the structural tokens. This projected text vector is prepended to the structural token embeddings to form a text-protein joint representation. This joint representation is then processed by the SaProt-T block, repeated *N* times, to predict the final amino acid sequence.

**Extended Data Fig. 2:**
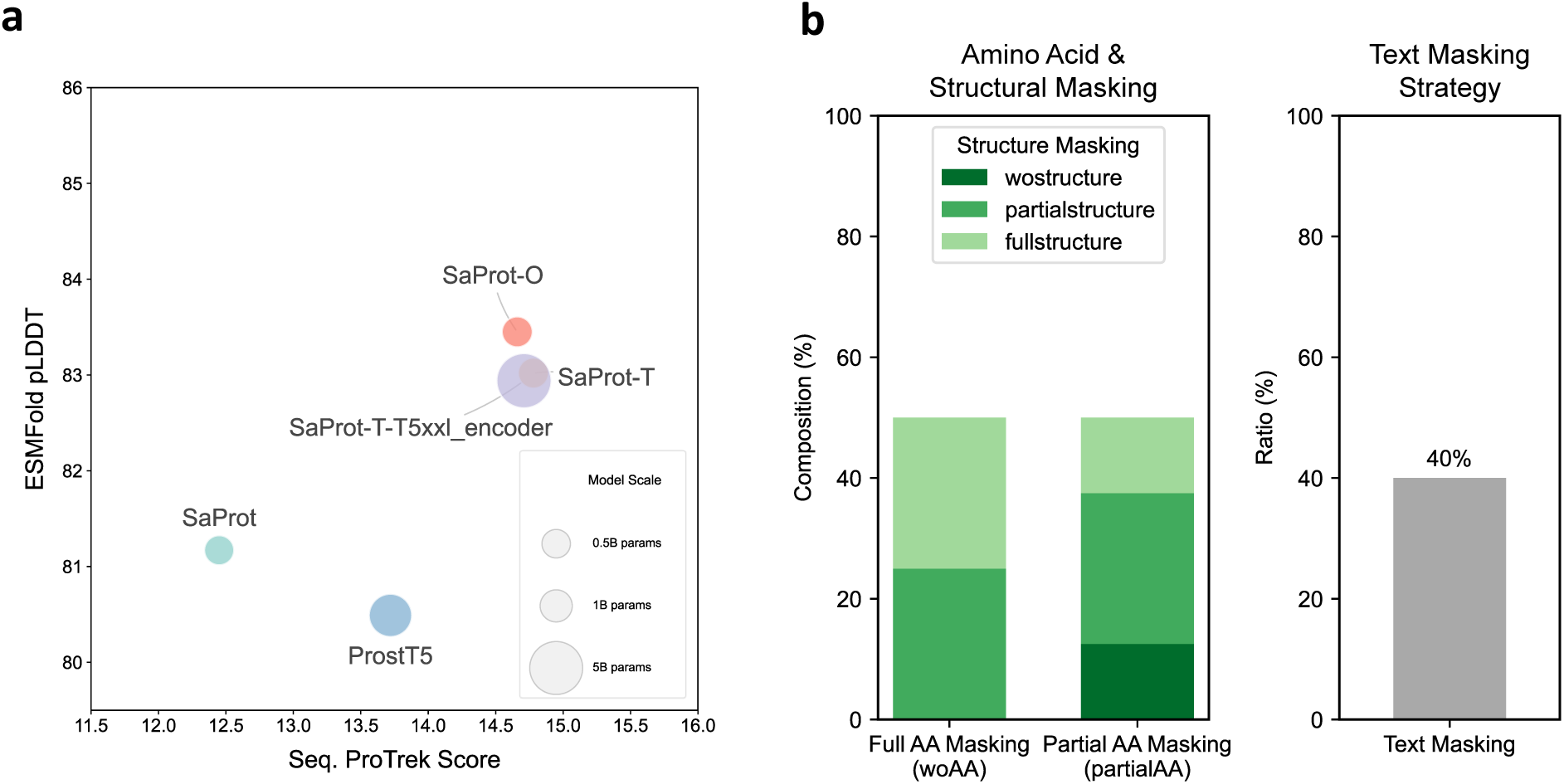
Text conditioning is crucial for the sequence design module. **a:** Ablation study comparing the performance of SaProt-T and SaProt-O (760M parameters) against the original SaProt (650M), the ProstT5 baseline, and the scaled SaProt-T-T5xxl encoder (∼5B). Performance is evaluated based on textual alignment (x-axis) and predicted foldability (y-axis). The results show that our text-conditioned models (SaProt-T/O) outperform the original SaProt and the ProstT5 baseline, particularly in maintaining structural foldability. The size of each bubble corresponds to the number of model parameters, as indicated in the legend. **b:** Composition of the omni-modal training strategy for the SaProt-O model. This strategy involves a varied combination of masking applied to the amino acid (AA) sequence, the structural tokens, and the text prompt to train a flexible, multi-purpose design model. The left panel details the distribution of structural token masking (no, partial, or full masking) under two different AA masking conditions. The right panel shows that the text prompt is masked in 40% of the training examples.

**Extended Data Fig. 3:**
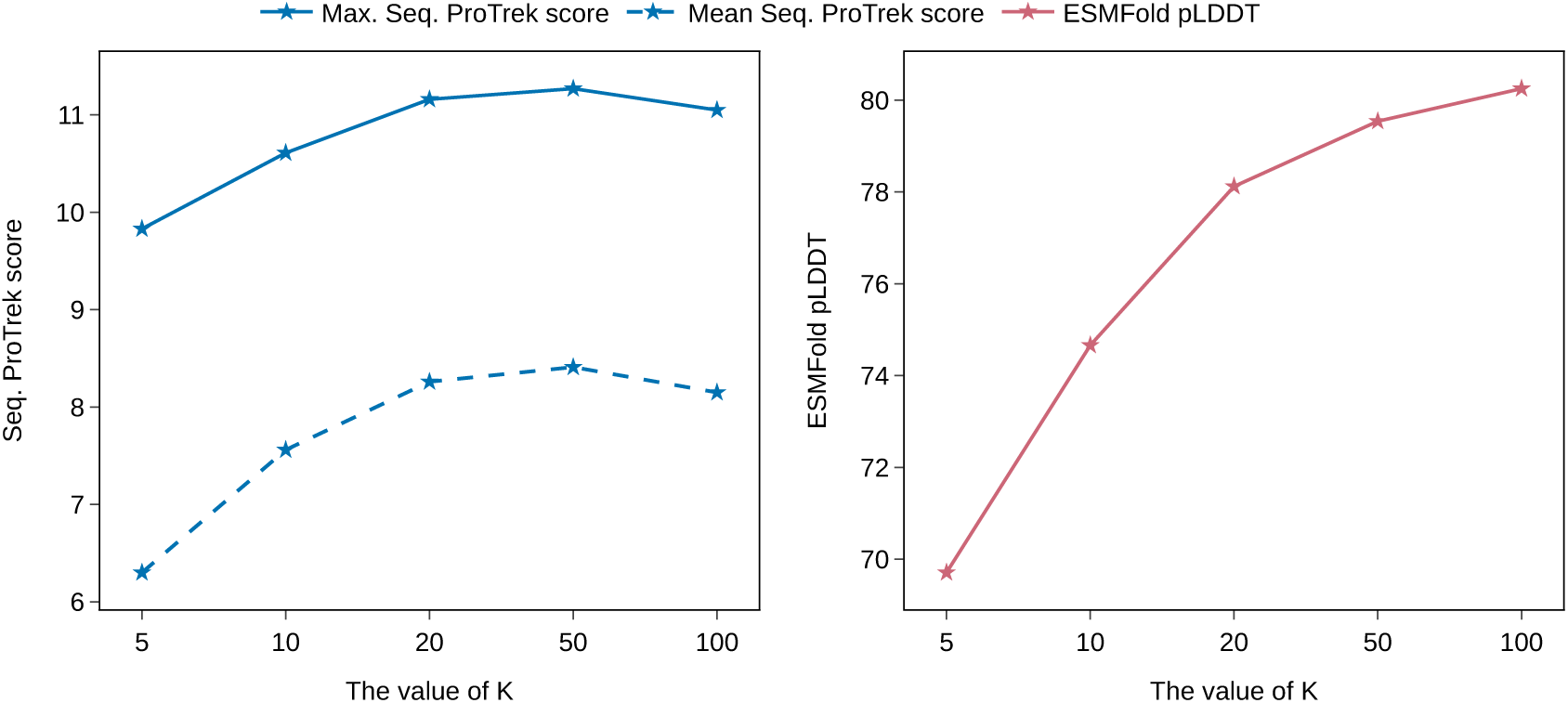
Determining the optimal number of sampling candidates (K). Performance of the top-ranked Pinal designs on the “hard” test set as a function of the number of sampled candidates (*K*). For each value of *K*, we generate *K* candidates, rank them by their joint log-probability, and then evaluate the top-ranked design. Left panel: textual alignment, measured by Max. and Mean Seq. ProTrek score. Right panel: predicted structural plausibility, measured by ESMFold pLDDT. Both textual alignment and foldability improve as *K* increases from 5 to 50. Further increasing *K* to 100 yields only marginal gains (or a slight decrease in ProTrek score) while substantially increasing computational cost. Based on this analysis, *K*=50 is used as the standard setting for all experiments, offering a robust balance between design performance and inference efficiency.

**Extended Data Fig. 4:**
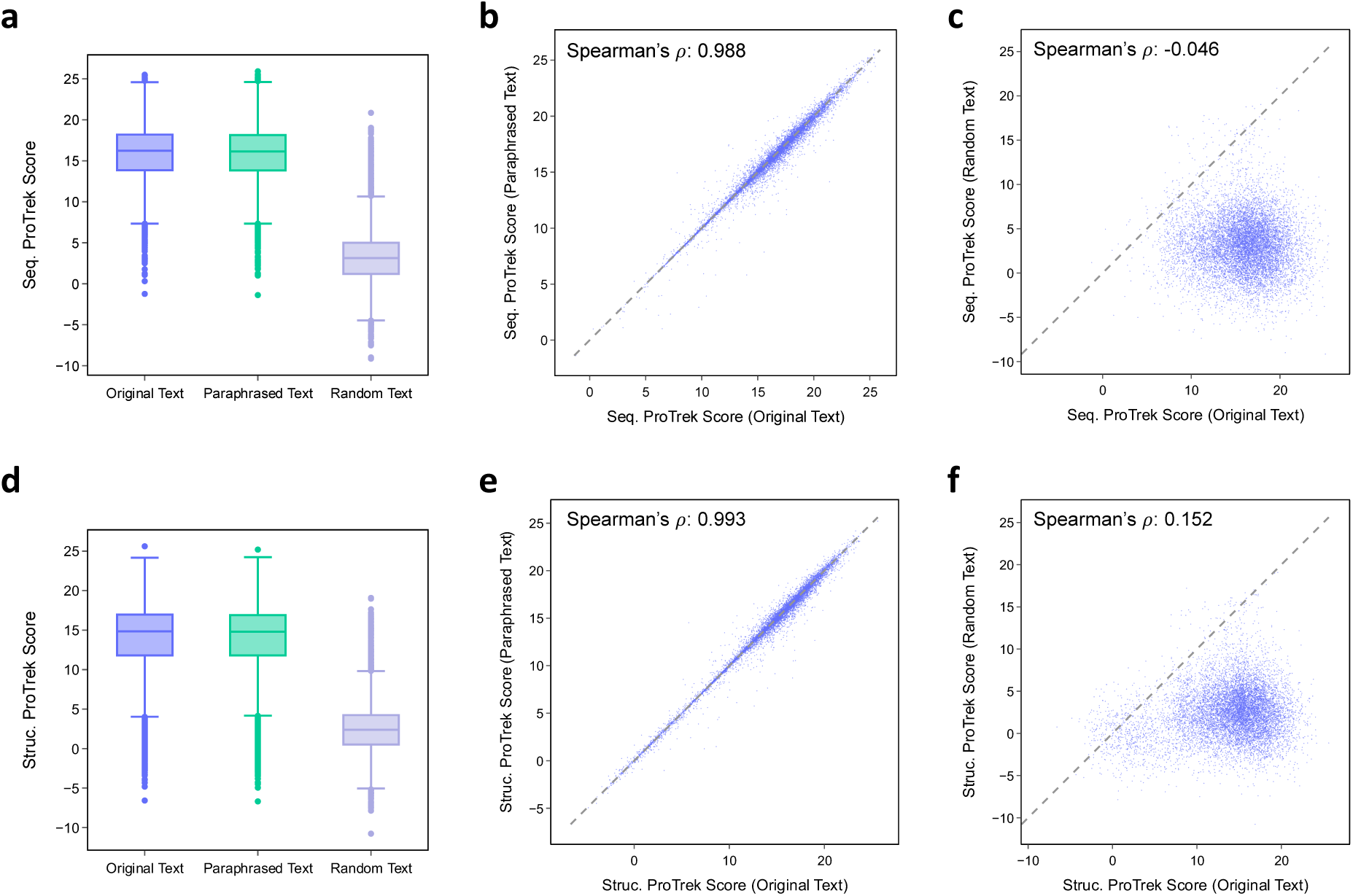
ProTrek score is a robust and discriminative metric for both text-sequence and text-structure alignment. The figure validates the ProTrek score’s performance using a test set of 1,000 protein-text pairs from Swiss-Prot. The top row (a-c) evaluates the text-to-sequence score (Seq. ProTrek), while the bottom row (d-f) evaluates the text-to-structure score (Struc. ProTrek). (a, d) Box plots show the distribution of ProTrek scores for proteins paired with their original descriptions, LLM-paraphrased descriptions, and randomly assigned (mismatched) descriptions. For both sequence and structure, the scores for original and paraphrased text are high and similar, while scores for random text are lower, demonstrating the metric’s discriminative power. Box plots show the median (centre line), 25th–75^th^ percentiles (box), the most extreme values within 1.5× the interquartile range (whiskers), and outliers (points). (b, e) Scatter plots comparing the scores of original versus paraphrased descriptions for each protein. The strong linear relationship, confirmed by a Spearman’s rank correlation coefficient (*ρ*) of *>*0.98 for both sequence and structure, demonstrates that the metric is robust to variations in wording and captures core semantic meaning. (c, f) Scatter plots comparing the scores of original versus randomly assigned descriptions. The near-zero correlations (*ρ* ≈ 0) further confirm that the ProTrek score is a valid metric that reliably distinguishes between relevant and irrelevant protein-text pairings. The original ProTrek study also demonstrated that its score selectively responds to biologically significant keywords while ignoring non-functional terms. Building on this capability, a recent benchmark paper ^48^ adopted ProTrek as a key metric for evaluating the alignment between protein sequences and their textual descriptions.

**Extended Data Fig. 5:**
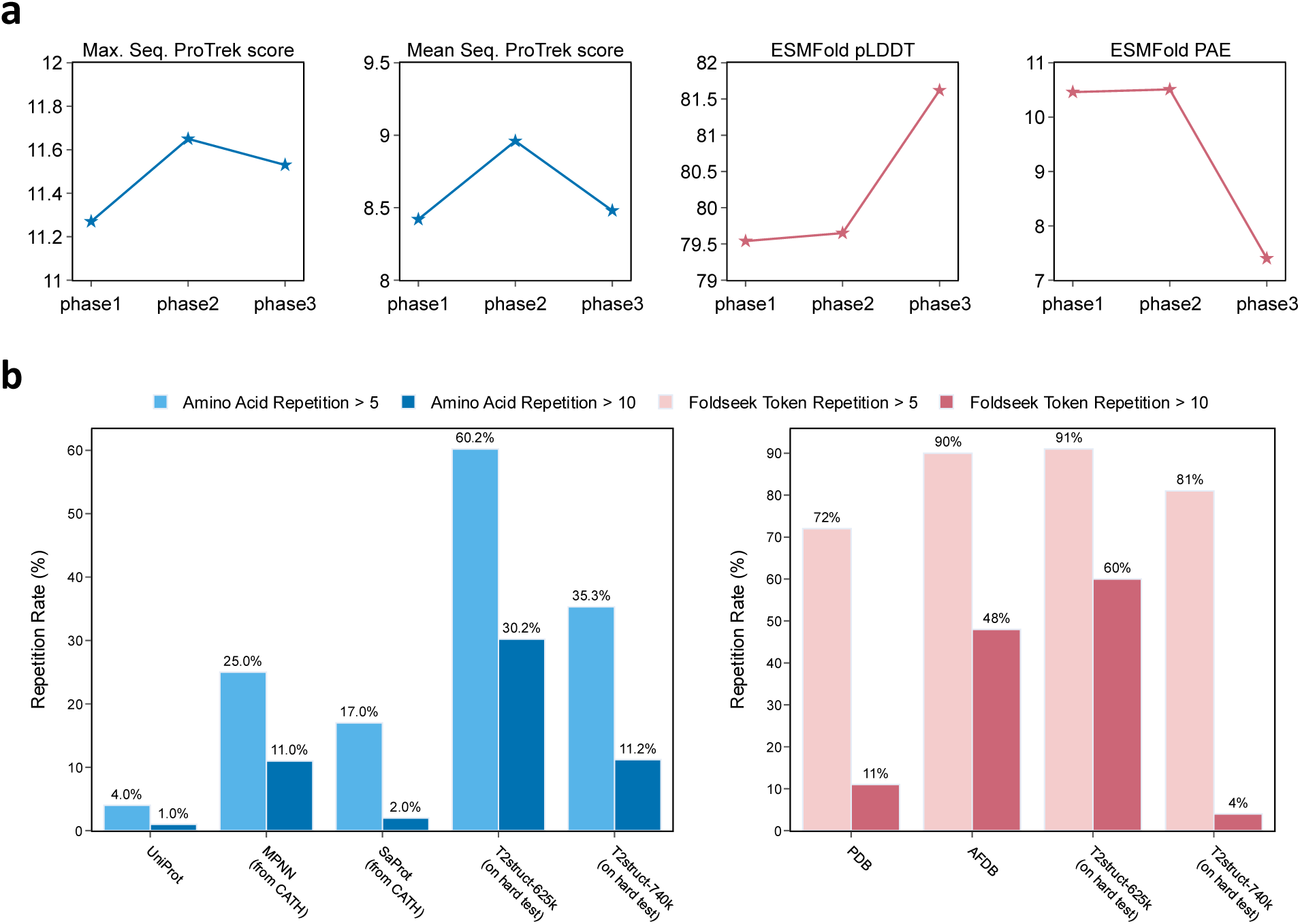
The three-phase training improves model performance and mitigates sequence repetition. **a,** Performance of the 15.5B T2struct model on the “hard” test set across three distinct training phases. Phase 1 (320k steps) represents initial training; Phase 2 (625k steps) incorporates data filtering by ProTrek score; Phase 3 (740k steps) is a final fine-tuning step on higher-quality data filtered to exclude low-pLDDT scores and high token repetition. The plots demonstrate that the third phase improves protein foldability, while incurring only a minor, acceptable trade-off in textual alignment (ProTrek score). **b,** Analysis of the sequence repetition phenomenon that motivated the third training phase. The left panel shows that the model after Phase 2 exhibited a high rate of amino acid repetition (≥5) (e.g.,“AAAAA”) at 60.2%, far exceeding that of natural proteins (UniProt, 4.0%). The final model after Phase 3 substantially reduces this rate to 35.3%. The right panel illustrates the effectiveness of the Phase 3 training data-filtering strategy. The high rate of Foldseek token repetition in the Phase 2 model’s outputs (91%) correlates with the high rate found in the AFDB training data (90%). The refined training strategy after Phase 3 successfully reduces the rate of token repetition greater than 10 to just 4%, thereby substantially resolving the Foldseek structure token repetition issue.

**Extended Data Fig. 6:**
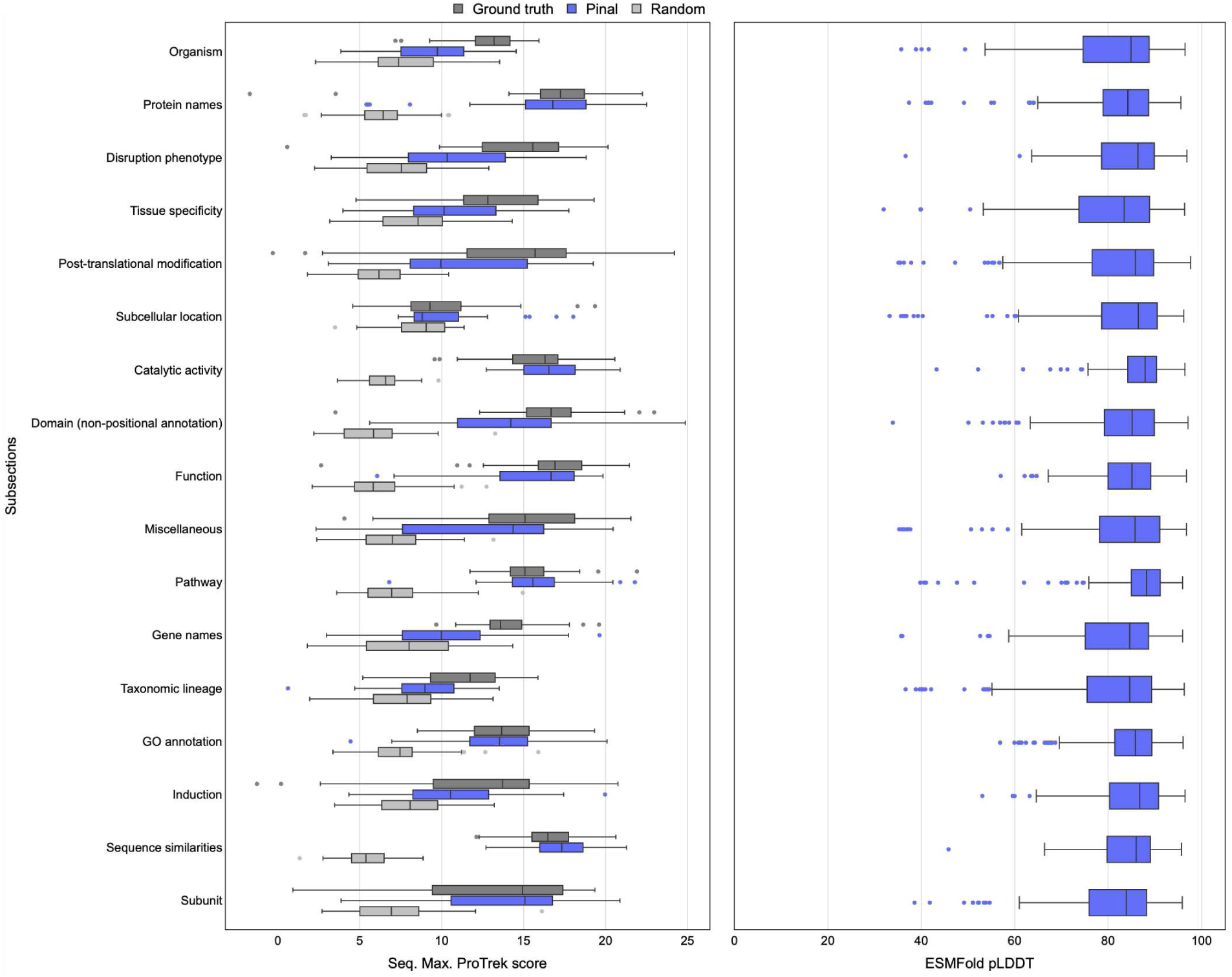
Detailed performance analysis across diverse protein function descriptions. The figure evaluates Pinal’s design performance across 17 distinct functional categories curated from Swiss-Prot. Left panel: Analysis of textual alignment, measured by the maximum sequence ProTrek score (*n* = 50 functional prompts per subsection). The performance of Pinal’s top-ranked design (blue) is compared against a random baseline (light grey) and the ground-truth protein (dark grey) for each category. Pinal consistently outperforms the random baseline across all categories and approaches ground-truth performance, demonstrating robust language understanding. Right panel: Predicted structural plausibility of the corresponding Pinal-designed proteins, measured by ESMFold pLDDT (*n* = 250 Pinal-designed proteins per subsection). The results indicate that Pinal generates proteins with high predicted foldability (median pLDDT is typically *>* 80) across all functional description types, from concrete categories like “Protein names” to more abstract ones like “Disruption phenotype”. Box plots show the median (centre line), 25th–75th percentiles (box), the most extreme values within 1.5× the interquartile range (whiskers), and outliers (points).

**Extended Data Fig. 7:**
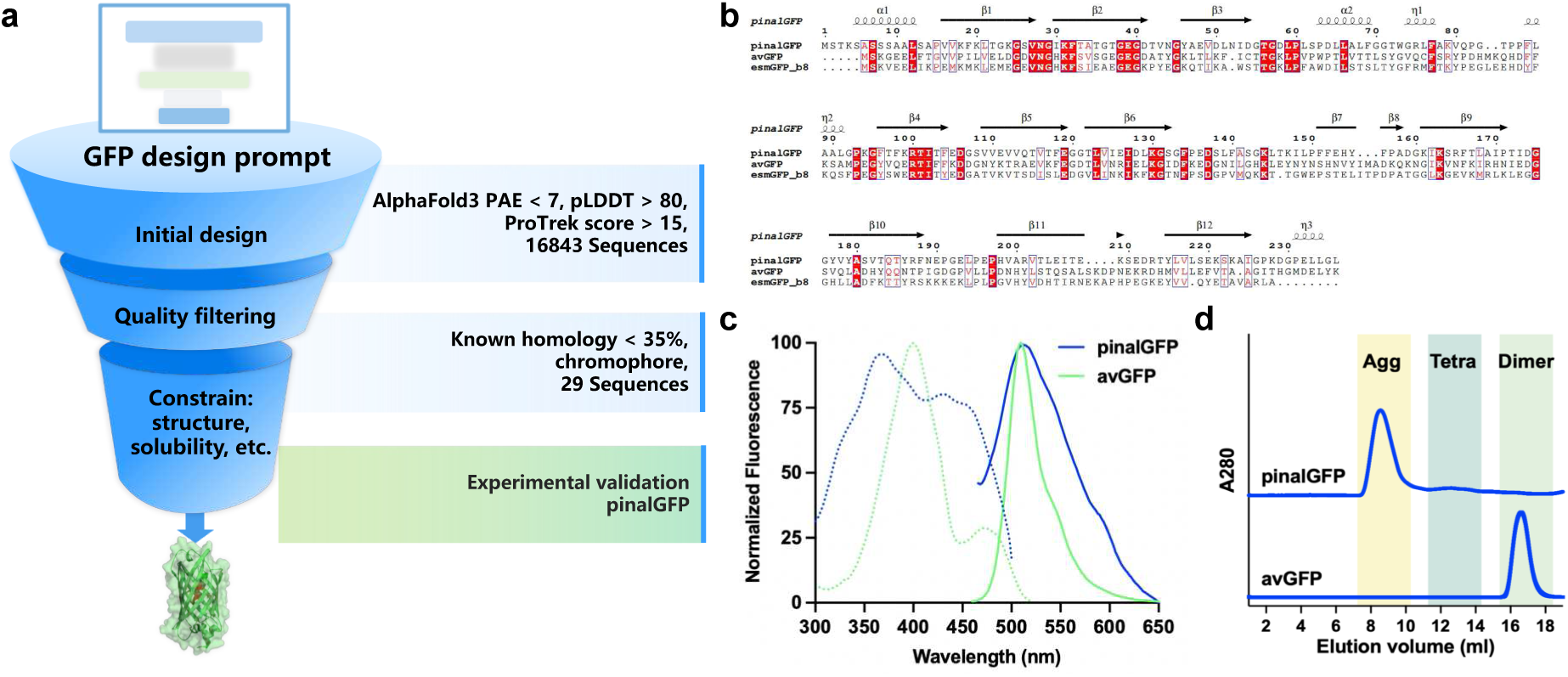
Computational design, screening, and characterization of pinalGFP. **a,** Flowchart illustrating the multi-tiered computational filtering strategy applied to the initial Pinal-generated sequences. This pipeline culminated in the selection of 29 candidates for experimental validation. **b,** Multiple sequence alignment of pinalGFP, avGFP, and esmGFP-b8. The alignment highlights the substantial sequence divergence of pinalGFP from both the natural protein and the other synthetic variant. **c,** Normalized fluorescence excitation (dashed lines, *λ*_em_ = 510 nm) and emission (solid lines, *λ*_ex_ = 460 nm) spectra of pinalGFP (blue) and avGFP (green). **d,** Size-exclusion chromatography profiles of purified GFP variants on a Superdex 200 column. Elution volumes reflected their oligomeric states in solution: avGFP was predominantly dimeric, whereas pinalGFP exhibited a more complex profile dominated by higher-order multimers.

**Extended Data Fig. 8:**
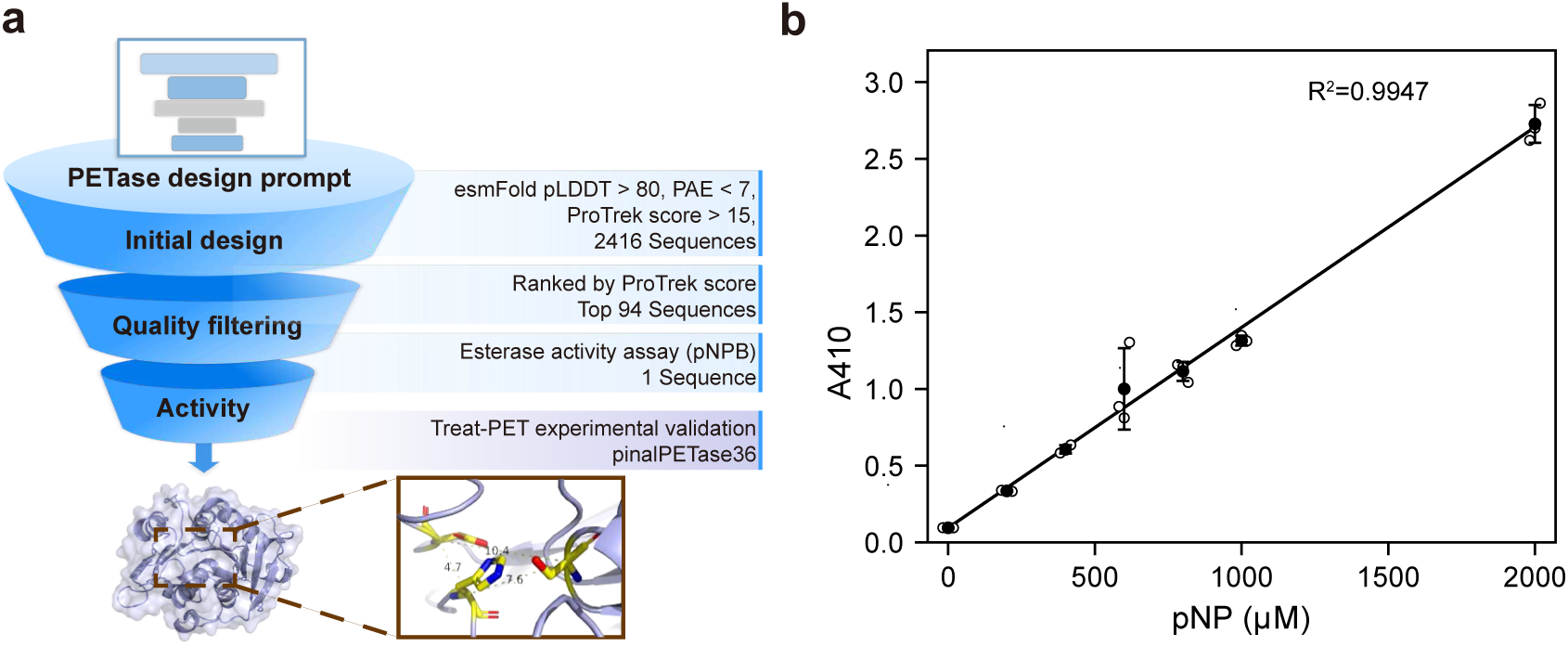
Computational screening pipeline and pNP standard curve for pinalPETase identification. **a,** Flowchart illustrating the multi-tiered screening strategy used to identify the active PETase candidate from the initial pool of Pinal-generated sequences. Candidates were first filtered on the basis of predicted structural quality (ESMFold pLDDT *>* 80, PAE *<* 7) and text–protein alignment (ProTrek score *>* 15), yielding 2,416 sequences. The top 94 candidates, ranked by ProTrek score, were then subjected to esterase activity screening using the *p*-nitrophenyl butyrate (pNPB) assay. This workflow led to the identification of a single standout variant, pinalPETase36, which was subsequently advanced to PET hydrolysis validation. The inset shows the predicted active-site architecture of pinalPETase36. **b,** Standard curve for *p*-nitrophenol (pNP) used for quantification of esterase activity in the pNPB assay. Absorbance at 410 nm was plotted against pNP concentration, showing a strong linear relationship (*R*^2^ = 0.9947). Data are shown as mean ± s.d. (n = 3 independent measurements).

**Extended Data Fig. 9:**
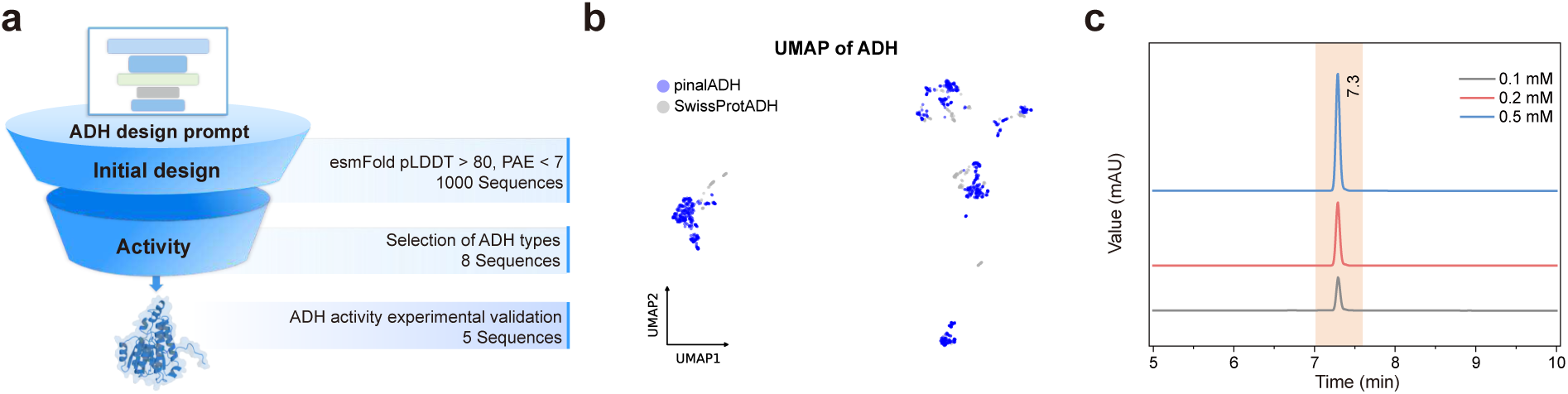
Computational design pipeline and analytical method for Pinal-designed ADHs. **a,** Flowchart of the computational screening and selection process for ADH candidates. From an initial library of 2,500 generated sequences, 1,639 candidates passing the feasibility filters based on structural confidence (ESMFold pLDDT *>* 80, PAE *<* 7) were identified, from which 1,000 sequences were randomly retained for downstream prioritization. From these, eight diverse candidates representing different ADH subfamilies were selected, leading to five successfully validated enzymes. **b,** UMAP of alcohol dehydrogenase (ADH) sequences. A total of 1,000 Pinal-designed ADH sequences (blue) were analysed together with natural ADH sequences from Swiss-Prot (gray; EC 1.1.1.1). Sequence embeddings were generated using ESM2-650M and projected into two dimensions by UMAP. The Pinal-designed sequences span a substantial region of the natural ADH sequence space. c, Representative HPLC chromatograms of derivatized acetaldehyde standards, corresponding to the product of the ADH-catalyzed reaction. Acetaldehyde standards at 0.1, 0.2, and 0.5 mM showed a consistent retention time of approximately 7.3 min.

**Extended Data Fig. 10:**
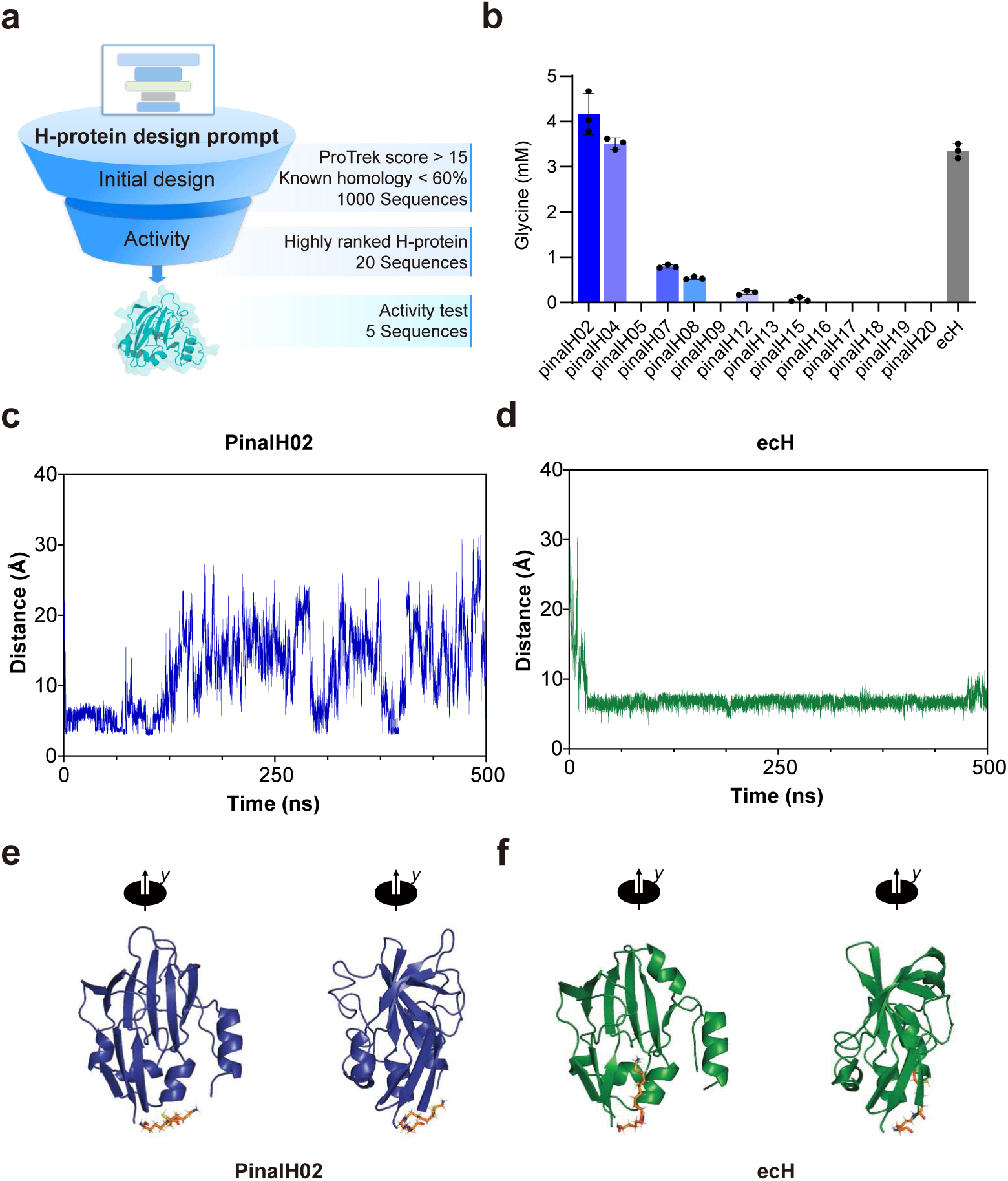
Computational design, screening, and mechanistic analysis of Pinal-designed H-proteins. **a,** Flowchart illustrating the computational design and screening pipeline for H-protein candidates. The initial generation was filtered to yield approximately 1,000 sequences that simultaneously satisfied the protein-text alignment criterion (ProTrek score *>* 15) and the sequence-divergence criterion (*<*60% sequence identity to known homologs). From this pool, 20 highly ranked candidates were selected, identifying five sequences with experimentally confirmed activity. **b,** Initial activity screening of 14 soluble Pinal-designed H-proteins compared to the native *E. coli* H-protein (‘ecH’). Glycine synthesis was measured after a 2 h reaction in a reconstituted Glycine Cleavage System (GCS). Data are shown as mean ± s.d. (n = 3 biologically independent reactions). **c-f,** Molecular dynamics (MD) simulations comparing the conformational dynamics of the lipoamide arm in the top-performing ‘pinalH02’ and the native ‘ecH’. **c,** Trajectory of the distance between the C*δ* atom of Glu27 in ‘pinalH02’ and the nitrogen atom of the lipoamide arm over 500 ns. The plot shows that the arm frequently samples extended conformations far from the protein core. **d,** Corresponding distance trajectory for ‘ecH’, plotting the distance between the C*δ* atom of Glu12 and the arm’s nitrogen atom over 500 ns. The arm remains stably sequestered close to the protein body. **e,** A representative snapshot of the ‘pinalH02’ conformation at 200 ns, illustrating the solvent-exposed lipoamide arm. **f,** A representative snapshot of ‘ecH’ at 200 ns, showing the aminomethyl lipoate arm locked within the protein’s hydrophobic cavity.

