## Supplementary Information for "Programmable design of functional proteins from natural language"

#### Supplementary Materials Guide:

Supplementary Method 1: Benchmarking against concurrent direct text-to-sequence models

Supplementary Method 2: Computational evaluation of generative diversity and novelty across data scales

Supplementary Method 3: Design and experimental validation of pinalGFP-g

Supplementary Method 4: Design and experimental validation of pinalH-R2

Supplementary Note 1: Interpretation of data scaling effects on foldability, novelty, and diversity

Supplementary Note 2: Target-dependent generation profiles across four experimentally validated protein targets

Supplementary Fig. 1-8

Supplementary Table 1-15

### 1 Benchmarking against concurrent direct text-to-sequence models

We included ProDVa [81], a recent text-to-sequence protein design model, as an additional baseline in the Supplementary Information. Although ProDVa appeared on arXiv approximately nine months after the Pinal bioRxiv preprint, we included this comparison because it provides an informative recent baseline. ProDVa represents a distinct design paradigm from token-by-token text-to-sequence generation. Specifically, ProDVa dynamically retrieves biologically meaningful protein fragments of varying lengths, including domains, protein families, and functional sites annotated by InterPro, conditioned on the input textual description. It therefore incorporates structural and evolutionary priors from naturally occurring protein fragments during generation.

We benchmarked our full 16B-parameter Pinal system against two ProDVa variants, ProDVa-CAMEO and ProDVa-Molinst, using the same comprehensive evaluation framework as in our main analysis. As shown in Supplementary Fig. 1, Pinal consistently outperforms both ProDVa variants across the major evaluation metrics for both short and long text prompts. These include functional alignment, measured by ProTrek score; structural accuracy, measured by GT-TMscore; and generative foldability, assessed by ESMFold pLDDT and PAE. In fact, we note that ProDVa performs relatively well on prompts resembling its training distribution, such as Swiss-Prot-style functional descriptions, but shows reduced performance under more open-ended or compositionally complex prompts. This trend is consistent with its more restricted training regime, which uses approximately 712K text-protein pairs, primarily from Swiss-Prot and Mol-Instructions, compared with the substantially larger training corpus used for Pinal. Overall, this comparison supports the conclusion that Pinal's larger and more diverse training corpus, spanning protein sequence, structure and functional descriptions, enables stronger functional alignment and foldable sequence generation across diverse natural-language design prompts.

We used the official implementation and publicly released ProDVa-CAMEO and ProDVa-Molinst checkpoints. Protein fragments were sampled using the 16 highest-ranked retrieved documents. Sequences were sampled with top-$k = 950$, a temperature of 0.7 and a maximum output length of 256 tokens.

### 2 Computational evaluation of generative diversity and novelty across data scales

To systematically assess how increasing the scale of training data influences the generative landscape of Pinal, we evaluated the sequence diversity, structural diversity, and novelty of the generated proteins on the standard test set (Supplementary Fig. 3). We first applied a rigorous quality filter to ensure that our diversity and novelty analyses were not artificially inflated by poorly folded or functionally irrelevant sequences. Only generated proteins that exhibited high structural confidence (ESMFold pLDDT $> 70$) and strong functional alignment with the text prompts (ProTrek score $> 10$) were retained and subsequently analyzed following a strategy similar to the DISCO methodology [82]. For each training scale, the same standard test prompts, generation settings, and number of initial candidate designs were used, and all downstream diversity and novelty analyses were performed on designs passing the same quality filters.

Sequence diversity was quantified by clustering the generated sequences with MMseqs2 using the following command:

```
mmseqs easy-cluster input.fasta clust tmp --min-seq-id 0.3 -c 0.8 --cov-mode 0
```

Structural diversity was measured by clustering the corresponding backbones with Foldseek using TM-score-based alignment:

```
foldseek easy-cluster input_dir clust tmp --alignment-type 1 --cov-mode 0 \
--min-seq-id 0 --tmscore-threshold 0.5
```

The total number of distinct clusters was recorded as a count-based measure of diversity yield among quality-filtered designs.

For the novelty assessment, we evaluated the sequence and structural similarity of the quality-filtered designs against UniRef100 and experimentally determined PDB structures, respectively. Sequence novelty was determined by searching the generated sequences against the comprehensive UniRef100 database:

```
mmseqs easy-search input.fasta uniref100 result.m8 tmp --gpu 1
```

Structural novelty was assessed by searching the predicted structures against all experimental entries in the Protein Data Bank (PDB):

```
foldseek easy-search input_dir pdb result.m8 tmp --alignment-type 1 --tmscore-threshold 0.0
```

We reported the highest query-normalized TM-score as the PDB-TM. Designs were classified as sequence-novel if their highest sequence identity to UniRef100 was below 0.3, and as structurally novel if their highest query-normalized TM-score against PDB structures was below 0.5. In Supplementary Fig. 3, sequence and structural novelty were reported separately as absolute counts. All unlisted MMseqs2 and Foldseek parameters were kept at their default settings.

### 3 Design and experimental validation of pinalGFP-g

#### 3.1 Pinal-generated $\beta$-barrel scaffolds support fluorescence after chromophore grafting

Although the main text demonstrates that Pinal can generate functional GFP candidates directly from natural-language descriptions, practical protein design often benefits from combining generative models with domain knowledge and minimal expert intervention. To illustrate this use case, we report an early experiment in which Pinal-generated $\beta$-barrel scaffolds were functionalized by manual grafting of a canonical chromophore motif and subsequently subjected to experimental validation.

An initial pool of 1,000 GFP candidate sequences generated by Pinal had already passed basic preliminary quality filters, including a ProTrek text-to-sequence matching score above 15 and high structural confidence from AlphaFold3 (average pLDDT $>$ 80 and PAE $<$ 7). This filtered set was further refined to 80 high-confidence candidates based on optimal physicochemical properties, such as sequence length (215–245 amino acids), predicted solubility (Protein-Sol $>$ 0.7), and thermodynamic stability (PyRosetta energy score $<$ -50 REU). To ensure structural diversity, we performed K-means clustering on PCA-reduced biophysical features to select 20 representative designs. For targeted functionalization, these representatives were structurally aligned to avGFP (PDB: 2WUR), and the essential Thr65–Tyr66–Gly67 (TYG) motif was manually engineered into the homologous positions using PyMOL. Following a final assessment of structural integrity (MolProbity clashscore $< 21$) and stability, 15 distinct candidates were ultimately selected for experimental validation (Supplementary Fig. 6a).

These 15 sequences shared $< 30\%$ identity with known fluorescent proteins but were all confidently predicted to adopt the characteristic $\beta$-barrel architecture (Supplementary Fig. 6b,d). Remarkably, 11 of the 15 proteins (73%) expressed in a soluble form. Screening this pool of soluble proteins for function revealed one highly active variant, hereafter referred to as pinalGFP-g (grafted), which produced green fluorescence (460 nm, 510 nm) visible to the naked eye (Supplementary Fig. 6c,f). Whole-cell fluorescence measurements showed that pinalGFP-g achieved 50% of the fluorescence intensity of the highly optimized esmGFP and 20% of avGFP within 24 h. Notably, this performance under minimal constraints surpassed the equivalent design, esmGFP-b8 from ESM3, which was non-fluorescent at the same 24-h time point due to slow maturation (Supplementary Fig. 6e). Analytical gel filtration showed that while avGFP and esmGFP are primarily dimeric, pinalGFP-g exists in a more complex oligomeric state, with fluorescence confined to the dimeric fraction exhibiting a quantum yield of 0.44 (Supplementary Fig. 6g). This oligomeric behavior, also observed for pinalGFP, highlights a clear avenue for future optimization.

These results demonstrate that Pinal generates $\beta$-barrel scaffolds capable of supporting chromophore maturation upon introduction of the canonical TYG motif, complementing the autonomous design described in the main text.

##### 3.2 Whole-cell fluorescence screening and quantum yield determination of pinalGFP-g

Whole-cell fluorescence was screened in 24-well plates. After 20 h of induction at 30°C, cells were pelleted, resuspended in PBS (pH 7.4), and transferred to 96-well plates for measurement on an Agilent Microplate Reader (excitation 460 nm, emission 510 nm). Fluorescence intensity was normalized to cell density ($OD_{600}$). For quantum yield determination of purified proteins, excitation and emission spectra were recorded from 300–520 nm and 460–650 nm, respectively. Quantum yields were determined relative to the avGFP standard (QY = 0.79) using the equation:

$$QY_i = \frac{\int_{460\,\mathrm{nm}}^{650\,\mathrm{nm}} (F_i)\, d\lambda}{A280_i} \cdot \frac{A280_{\mathrm{avGFP}}}{\int_{460\,\mathrm{nm}}^{650\,\mathrm{nm}} (F_{\mathrm{avGFP}})\, d\lambda} QY_{\mathrm{avGFP}}$$

#### 4 Design and experimental validation of H-protein-R2

To further assess the experimental validity of Pinal-designed H-proteins, our experimental collaborators performed an additional round of H-protein design and characterization, referred to as pinalH-R2. This follow-up experiment used the same overall Pinal-based generation procedure and a similar in silico filtering workflow as those used for the H-protein designs described in the main text.

Briefly, candidate H-protein sequences were generated using the two-stage Pinal design procedure described in the Methods section. Given the H-protein design prompt, Pinal first generated structure-token sequences using T2struct with multinomial sampling at a temperature of 2.0. Each generated structure-token sequence was subsequently decoded by SaProt-T into an amino-acid sequence using multinomial sampling at a temperature of 0.3. In total, 6,250 new H-protein candidates were generated in this round.

The generated candidates were then evaluated using similar downstream in silico criteria. Specifically, candidates were filtered based on predicted structural confidence, text-protein alignment and redundancy relative to the model-training corpus, requiring AlphaFold3 predicted PAE $< 7$, predicted pLDDT $> 80$, and ProTrek score $> 15$. In contrast to the main-text H-protein validation set, where an additional sequence-identity threshold was applied to enrich for more divergent designs, this follow-up round did not impose a sequence-identity cutoff during candidate selection. The resulting candidates therefore provide a complementary validation set with a slightly broader sequence-identity distribution, as summarized in Supplementary Table 14. After in silico filtering, our experimental collaborators selected 98 candidate genes for synthesis and experimental characterization, prioritizing candidates with higher ProTrek scores.

Experimental characterization was performed using the same reconstituted glycine-production assay described for the H-protein validation. Glycine concentrations were measured after 6 h. Among the 98 synthesized candidates, 86 showed soluble expression and were further evaluated for activity. 49 exhibited measurable glycine-production activity, and with over 20 outperforming the native E. coli H-protein (Supplementary Table 15, Supplementary Fig. 7).

### Supplementary Note 1: Interpretation of data scaling effects on foldability, novelty, and diversity

LLM-based rewriting was used primarily to increase the linguistic diversity and coverage of protein descriptions, rather than to directly optimize the foldability or novelty of generated proteins. The major source of sequence, structure, and functional diversity in the synthetic corpus comes from ProTrek-based retrieval, which expands the paired training data from the curated Swiss-Prot space to the much broader TrEMBL/AFDB protein space. Therefore, the ProTrek+LLM data augmentation strategy does not simply paraphrase existing annotations, but exposes the model to a substantially broader sequence–structure–function landscape.

The effects of this data scaling are expected to be most direct for text–protein alignment, novelty, and diversity. Indeed, increasing the scale of the synthetic corpus improves alignment-related metrics, including ProTrek score and GT-TMscore, and increases the number of quality-filtered generated proteins that are diverse or novel in sequence and structure space (Supplementary Fig. 3). These trends indicate that large-scale synthetic data enable the model to explore a broader accessible design space while maintaining structural plausibility and functional relevance.

In contrast, the relationship between data scale and foldability-related metrics, such as ESMFold pLDDT and PAE, is more complex. Foldability is influenced not only by the number of training examples, but also by the structural confidence and noise level of those examples. The curated Swiss-Prot subset has substantially higher average AF2 structural confidence than the much larger TrEMBL-derived synthetic subset. Thus, adding more ProTrek-augmented data substantially increases biological coverage, but may also introduce training examples with lower average structural confidence. This can attenuate or mask simple monotonic improvements in pLDDT- or PAE-based foldability metrics.

In addition, Pinal contains two core modules, T2struct and SaProt-T. SaProt-T is initialized from the pretrained SaProt model and therefore provides a strong prior for generating foldable amino acid sequences conditioned on structures. As a result, pLDDT and PAE reflect the combined effects of T2struct, SaProt-T, the SaProt pretraining data, the SaProt-T fine-tuning data, and the T2struct synthetic training data. A fully controlled scaling analysis of pLDDT and PAE would therefore require jointly scaling multiple data sources and model components, which is computationally prohibitive. Consistent with the importance of structural-data quality, fine-tuning T2struct on a higher-confidence structural subset improved the foldability of generated proteins (Extended Data Fig. 4a).

Together, these analyses suggest that synthetic-data scaling improves Pinal primarily by expanding the accessible sequence–structure–function space and strengthening natural-language alignment. Foldability-related metrics, by contrast, should be interpreted as outcomes of the complete Pinal framework, including data scale, structural-data quality, and pretrained sequence-generation priors, rather than as direct consequences of LLM rewriting or dataset size alone.

### Supplementary Note 2: Target-dependent generation profiles across four experimentally validated protein targets

Protein targets may differ substantially in their conditional generation profiles and in the sampling depth required to recover sequences similar to a specific design. We therefore examined the positions of experimentally validated designs within their corresponding PPL distributions and compared target-dependent differences in generation difficulty among the four protein targets.

In Supplementary Fig. 8, we show the target-specific PPL distributions, the corresponding ProTrek scores and predicted pLDDT values across PPL intervals, and the positions of the experimentally validated active designs. The validated active ADH and H-protein designs were located in relatively low-PPL regions, while still showing favourable ProTrek scores and predicted structural confidence, indicating that these sequences remain within the learned protein manifold. More broadly, Supplementary Fig. 8 illustrates that ProTrek score, predicted pLDDT and PPL provide complementary but partially correlated measures for evaluating designed sequences. Among the four targets, ADH, H-protein and GFP showed more favourable overall ProTrek-score and PPL profiles than PETase, potentially reflecting the greater generation difficulty of PETase.

This difference was particularly evident for PETase. When equally sized pools of 50,000 candidates generated under the same inference settings were clustered at 30% sequence identity, the ADH pool formed 743 clusters, whereas the PETase pool formed 20,196 clusters. This indicates that PETase candidates were distributed across many more sequence neighbourhoods, with fewer sequences per neighbourhood on average. Therefore, independently sampled PETase candidates are less likely to fall into the same local region as a particular validated design, such as pinalPETase36, explaining why a larger candidate pool may be needed to recover moderately similar variants.

One possible contributing factor is the limited representation of PETase- or PET-hydrolase-related functions in curated protein databases. For example, relatively few entries can be retrieved from Swiss-Prot using PETase- or PET hydrolase-related keywords. Such limited curated coverage may lead to a sparser and more diffuse conditional generation space for the PETase task, consistent with the broader PPL distribution in Supplementary Fig. 8 and the wider range of ProTrek scores observed for PETase candidates. Although this observation does not establish a direct causal relationship, it provides a possible explanation for the greater diversity of generated PETase candidates and the larger sampling depth required to recover sequences close to a specific validated PETase design.

### Software and web resources used in this study

| **Software or resource** | **URL** |
|---|---|
| Pinal | http://www.denovo-pinal.com/ |
| SaProt-T/O (Pinal) | http://113.45.254.183:9527/ |
| Matplotlib (v3.7.5) | https://matplotlib.org/ |
| AlphaFold3 | https://alphafoldserver.com/ |
| Foldseek | https://github.com/steineggerlab/foldseek |
| MMseqs2 | https://github.com/soedinglab/MMseqs2 |
| ProTrek | https://github.com/westlake-repl/ProTrek |
| Protein-Sol | https://www.novopro.cn/tools/prot-sol.html |
| PyRosetta | https://www.pyrosetta.org/ |
| BLAST | https://blast.ncbi.nlm.nih.gov/Blast.cgi |
| TM-align | https://zhanggroup.org/TM-align/ |
| PyMOL (v3.1.1) | https://pymol.org/ |
| MolProbity | https://molprobity.biochem.duke.edu/ |
| ChimeraX (v1.10) | https://www.cgl.ucsf.edu/chimerax/ |
| IQtree2 | https://github.com/iqtree/iqtree2/ |
| CD-HIT (v4.8.1) | https://github.com/weizhongli/cdhit/ |
| Clustal Omega (v1.2.4) | http://www.clustal.org/omega/ |
| iTOL (v7.2) | https://itol.embl.de/ |
| Autodock Vina (v2.0) | https://github.com/ccsb-scripps/AutoDock-Vina |
| BioRender | https://www.biorender.com/ |
| Chromeleon 7 | https://www.thermofisher.com/order/catalog/product/CHROMELEON7 |

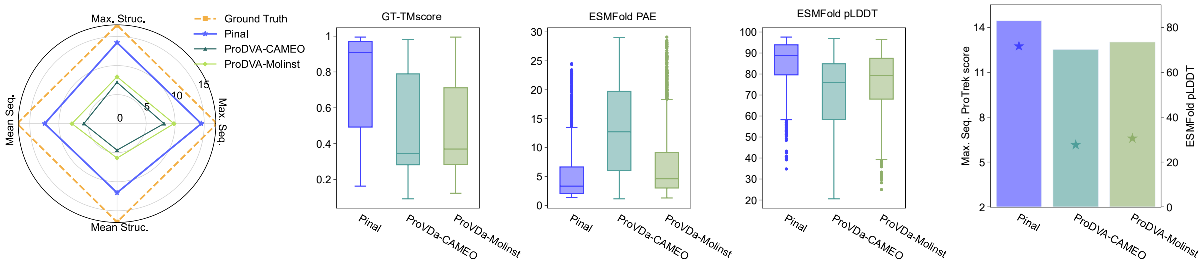


**Supplementary Fig. 1**: **Benchmarking Pinal against the recent text-to-sequence model ProDVa.** Following the evaluation framework established in the main text, we compared the 16B-parameter Pinal model with ProDVa-CAMEO and ProDVa-Molinst on the standard test set. From left to right: (1) Radar plot and GT-TMscore distribution assessing functional alignment and structural accuracy, respectively; (2) ESMFold PAE and pLDDT distributions evaluating structural foldability; (3) Bar and star plots illustrating the foldability and textual alignment (ProTrek score), respectively, conditioned on short text prompts. Across all generative metrics, Pinal consistently outperforms ProDVa, with distributions indicating more reliable generation quality across samples.

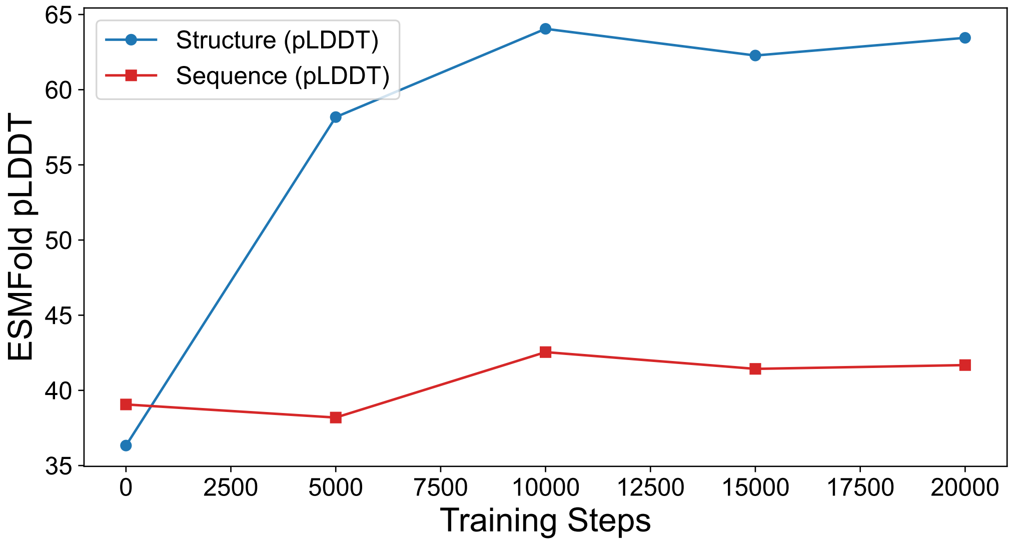


**Supplementary Fig. 2**: **Assessment of foldability in preliminary ablation experiments.** Comparison of ESMFold pLDDT scores for proteins generated by the text-to-structure model (blue circles) and the direct text-to-sequence model (red squares) over 20,000 training steps. For the text-to-structure model, generated Foldseek 3Di tokens were first decoded into amino acid sequences using SaProt, followed by structure prediction with ESMFold and assessment using ESMFold pLDDT, as pLDDT cannot be computed directly from 3Di token sequences.

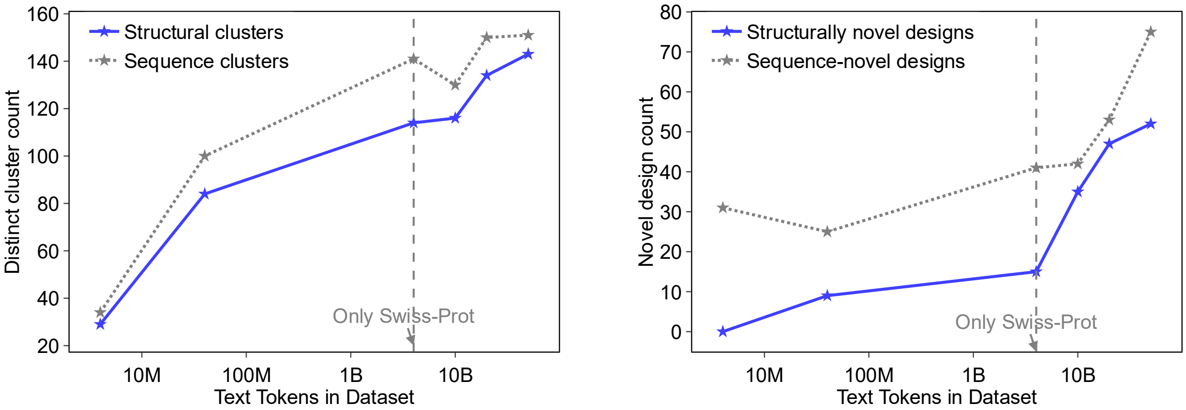


**Supplementary Fig. 3**: **Scaling of training data increases the yield of diverse and novel quality-filtered designs.** To ensure rigorous assessment, all generated candidates across different training scales were pre-filtered for high structural confidence (ESMFold pLDDT > 70) and functional alignment (ProTrek score > 10) prior to downstream evaluation. **Left**, Diversity analysis of the generated pools from each training scale, quantified by the total number of distinct clusters in structural (blue) and sequence (grey) spaces. **Right**, Novelty analysis of these quality-filtered proteins. Novelty is reported as the number of designed candidates classified as structurally novel relative to the PDB (blue solid line) or sequence-novel relative to UniRef100 (grey dotted line). The number of novel designs exhibits an overall increasing trend with training-data scale, suggesting that large-scale synthetic data augmentation is associated with an increased yield of novel, quality-filtered candidates and broader exploration of the accessible design space.

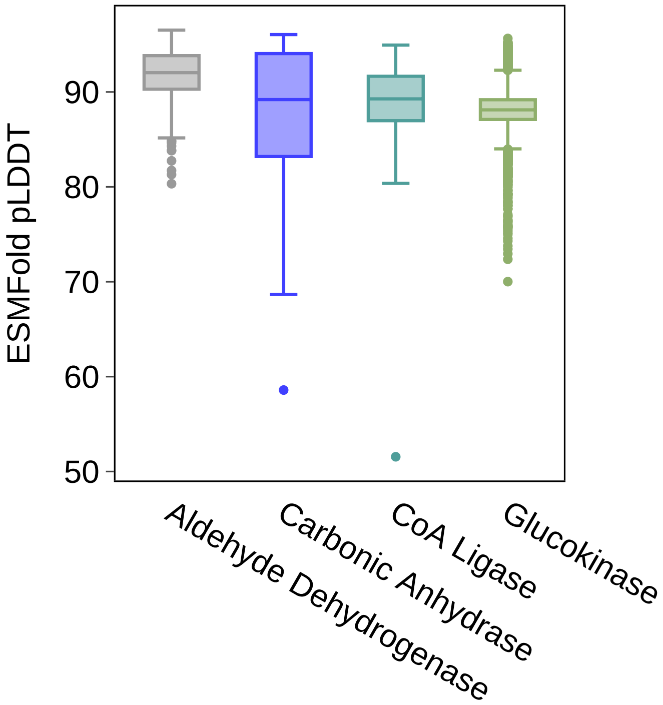


**Supplementary Fig. 4**: **Predicted structural confidence of Pinal-designed proteins used for phylogenetic analysis.** For each protein family, 5,000 sequences were generated by Pinal and filtered using a ProTrek score threshold of 15, retaining 2,382 aldehyde dehydrogenase, 843 carbonic anhydrase, 2,782 CoA ligase, and 4,547 glucokinase designs. The ESMFold pLDDT distributions of the retained sequences show high predicted structural confidence.

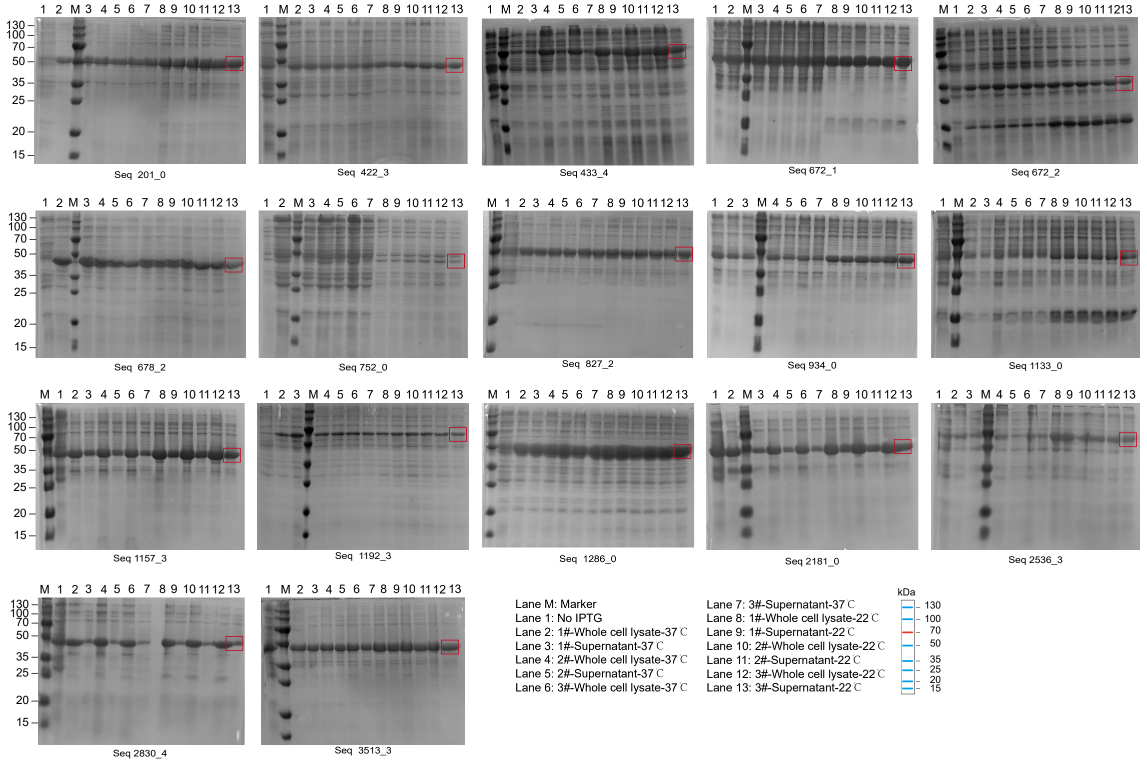


**Supplementary Fig. 5**: **Experimental validation of the soluble expression of novel Pinal-designed proteins.** SDS-PAGE analysis of the 17 successfully expressed novel proteins from a set of 20 computationally selected candidates. These proteins were designed by Pinal to have low sequence (<30%) and structural (<0.5 TM-score) similarity to any known proteins, demonstrating a high soluble expression rate of 85% for truly novel designs. For each designed protein (identified by its sequence ID), three independent colonies were tested for expression in *E. coli* BL21(DE3). Protein expression was induced with IPTG and conducted at two different temperatures (37°C and 22°C). The lanes for each gel are as follows: M, molecular weight marker (kDa); 1, uninduced whole-cell lysate; 2-7, whole-cell lysate and corresponding supernatant for three biological replicates induced at 37°C; 8-13, whole-cell lysate and corresponding supernatant for the same three replicates induced at 22°C. The red boxes highlight the target protein band in the supernatant fraction, confirming successful soluble expression. Recombinant proteins with N-terminal SUMO and Strep-tag II were expressed in *E. coli* BL21(DE3). Plasmids (5 ng) were transformed into 50 µL competent cells, heat-shocked at 42 °C for 60 s, recovered in LB, and plated on LB agar containing 50 µg mL$^{-1}$ kanamycin. Single colonies were grown overnight in 4 mL YP medium with kanamycin. Cultures were diluted 1:50 into fresh medium, grown to $OD_{600} \approx 0.6$, and induced with 0.1 mM IPTG at 22 °C or 37 °C for 20 h. Cells were harvested, lysed in PBS, and analyzed by SDS–PAGE. Uncropped and unprocessed SDS–PAGE image is provided in the Source Data file.

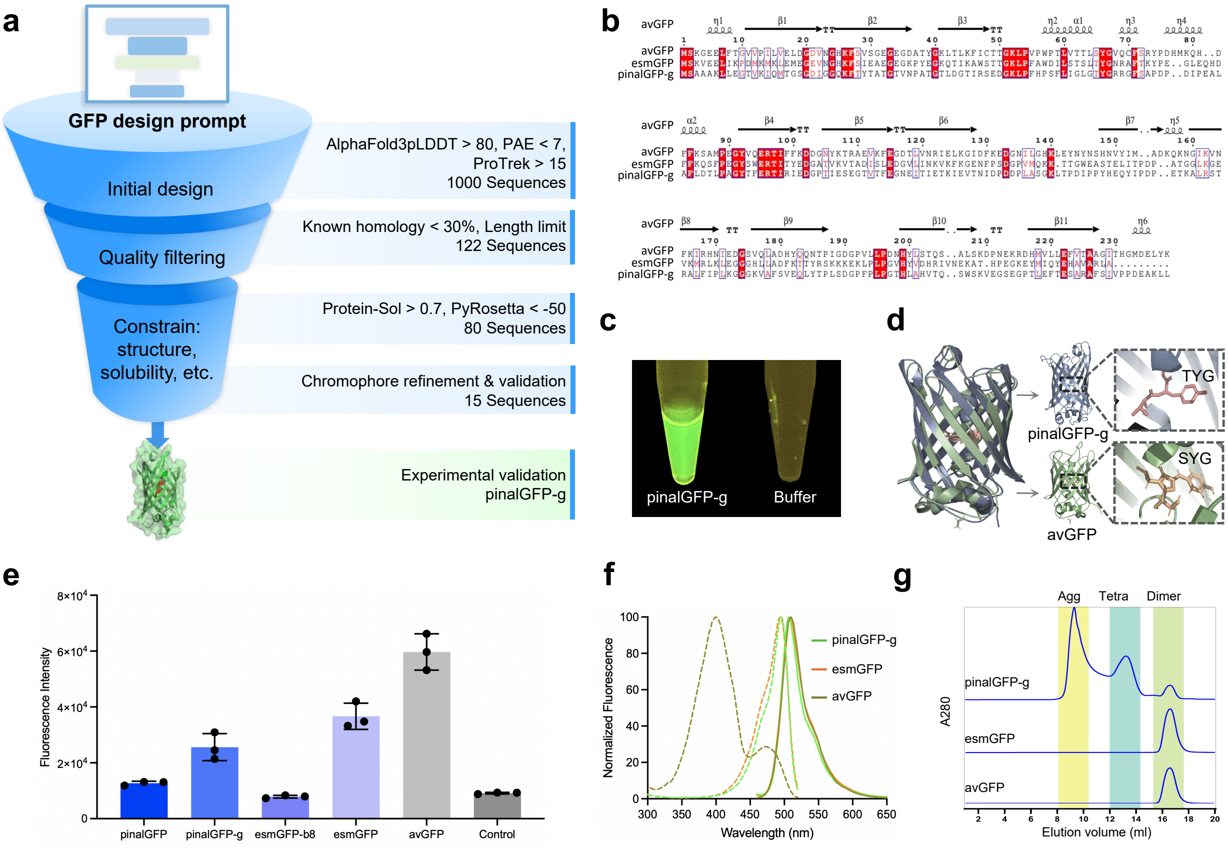


**Supplementary Fig. 6**: **Computational design, screening, and characterization of pinalGFP-g. a,** Flowchart illustrating the multi-tiered computational filtering strategy applied to the initial Pinal-generated sequences. The numbers indicate the count of candidate sequences remaining after each screening step. This pipeline culminated in the selection of 15 candidates for experimental validation. **b,** Multiple sequence alignment of pinalGFP-g, esmGFP, and avGFP. The alignment highlights the significant sequence divergence of pinalGFP-g from both the natural protein and the other synthetic variant. **c,** Visual evidence of purified pinalGFP-g (5 mg $mL^{-1}$) under 365 nm UV illumination, demonstrating the characteristic green fluorescence of the functional variant. **d,** Structural alignment of pinalGFP-g (blue) with avGFP (green, PDB: 2WUR) showing conservation of the $\beta$-barrel architecture. The engineered chromophore residues (T65–Y66–G67) are highlighted in the pinalGFP-g structure. **e,** Quantitative fluorescence analysis of GFP variants expressed in *E. coli* BL21(DE3) cells 24 h post-induction. Data are shown as mean ± s.d. (n = 3). **f,** Normalized fluorescence excitation (dashed lines, $\lambda_{em} = 510$ nm) and emission (solid lines, $\lambda_{ex} = 460$ nm) spectra. **g,** Size-exclusion chromatography profiles of the purified GFP variants on a Superdex 200 column. The elution volumes indicate their oligomeric states in solution: avGFP and esmGFP exist predominantly as dimers, whereas pinalGFP-g displays a more complex profile with fluorescence activity associated with the dimeric fraction.

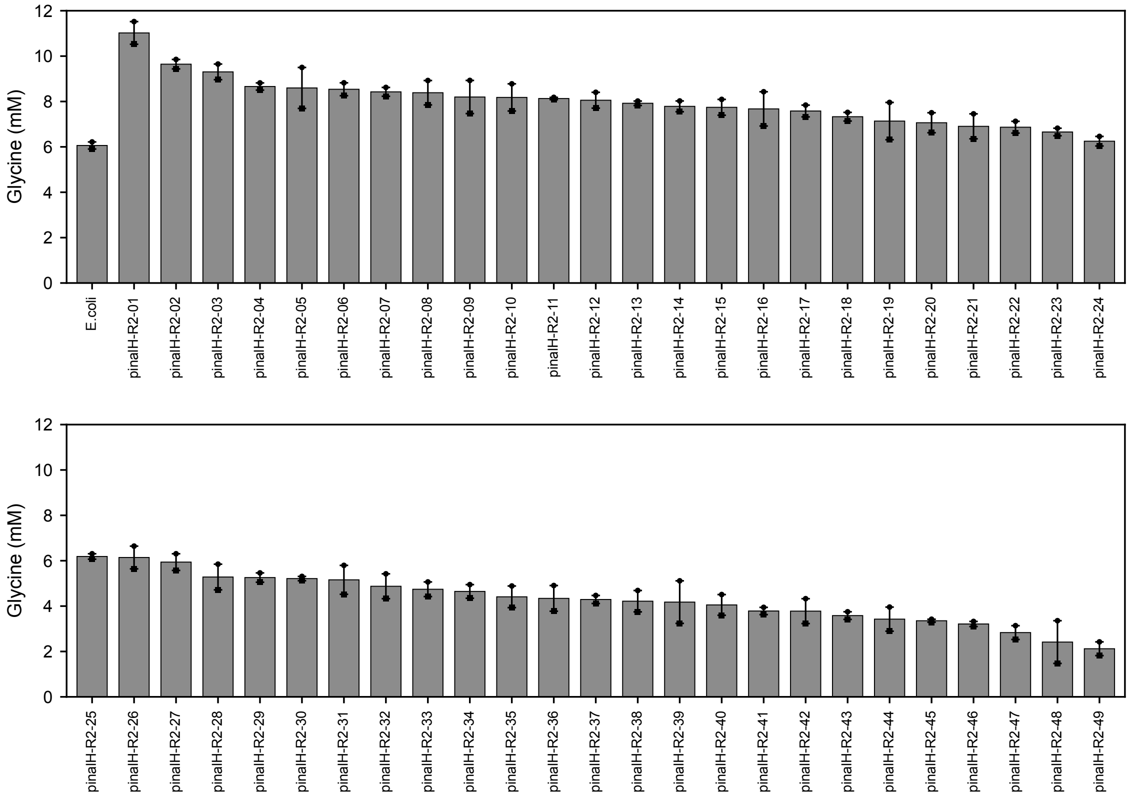


**Supplementary Fig. 7: Glycine production using Pinal-generated H proteins and the native E. coli H protein (ecH) in the second validation round.**

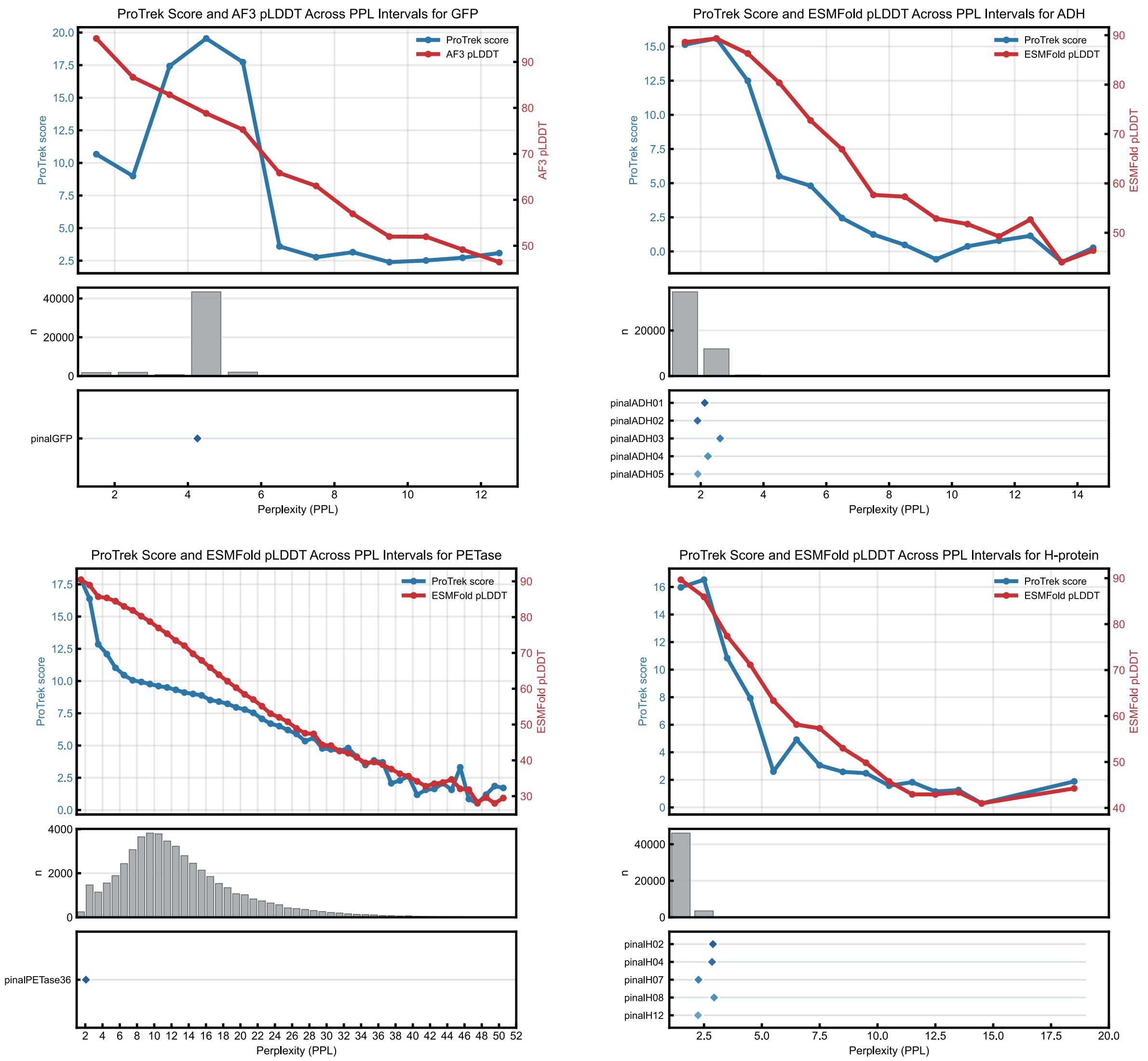


**Supplementary Fig. 8**: **Relationship between model perplexity and downstream ranking measures for designed protein candidate pools.** For each target, the upper panel shows the ProTrek score and predicted structural confidence across PPL intervals. The middle panel shows the distribution of designed sequences across PPL intervals. Diamonds in the lower panel indicate the PPL positions of experimentally validated designs for the corresponding target. For comparability across tasks, each reference distribution was constructed from 50,000 candidate sequences generated by Pinal.

**Supplementary Table 1**: A list of the 46 representative protein classes.

| | | |
|---|---|---|
| DNA/RNA polymerases | Intermediate filaments | GPCRs |
| Restriction enzymes | Growth factors | Receptor tyrosine kinases |
| Reverse transcriptases | Cytokines | Cell adhesion molecules |
| DNA repair enzymes | Chemokines | Heat shock proteins |
| Topoisomerases | Hormones | Molecular chaperones |
| Nucleases | Neurotransmitter receptors | Protein disulfide isomerases |
| Antibody | G-proteins | Peptidyl-prolyl isomerases |
| Fluorescent proteins | Second messengers | Luciferases |
| Actin | Transcription factors | $\beta$-galactosidase |
| Tubulin | Histones | Alkaline phosphatase |
| Collagen | RNA-binding proteins | Horseradish peroxidase |
| Keratin | DNA-binding proteins | Globular proteins |
| Fibronectin | Chromatin remodeling factors | Fibrous proteins |
| Laminin | Nuclear receptors | Motor proteins |
| Myosin | Ion channels | |
| Spectrin | Transporters | |

Note: Source databases for candidate retrieval were TrEMBL50 unless otherwise noted; FPbase was used for fluorescent proteins.

**Supplementary Table 2**: **Dataset statistics.** We have excluded proteins whose 3Di structures are not in Alphafold/UniProt database. Each dataset is assigned a specific sample weight for training.

| Dataset | Proteins | Protein-Text Pairs | Text Tokens | Sample Weight |
|---|---|---|---|---|
| SwissProt-Annot | 545K | 14M | 284M | 7 |
| SwissProt-Aug | 544K | 4M | 1.2B | 2 |
| SwissProt-LLM | 544K | 9M | 2.7B | 4 |
| TrEMBL50-ProTrek | 41M | 412M | 12B | 13 |
| TrEMBL50-ProLLM | 40M | 530M | 143B | 52 |
| InterPro | 113M | 791M | 4.4B | 15 |
| AFDB | 214M | – | – | 7 |

**Supplementary Table 3**: Hyperparameters for SaProt and T2struct models.

| Model name | Total Params | Encoder Params | Decoder Params | Text Length | Protein length | Learning rate | Total training steps |
|---|---|---|---|---|---|---|---|
| *SaProt* | | | | | | | |
| SaProt-T | 760M | – | – | 512 | 1024 | 5e-6 | 80,000 |
| SaProt-O | | | | 768 | 1536 | | 605,000 |
| *T2struct* | | | | | | | |
| T2struct-224M | 224M | 110M | 114M | 768 | 1024 | 1e-4 | 320,000 |
| T2struct-1.2B | 1.2B | 335M | 992M | | | | |
| T2struct-15.5B | 15.5B | 4.8B | 10.7B | | | 1e-5 | 740,000 |

#### Supplementary Table 4: Examples of descriptions from various data sources.

**Example Description (from SwissProt-Annot for *P95368*)**

The GO term related to this protein regarding molecular function envelopes shikimate 3-dehydrogenase (NADP+) activity. In this specific protein, position 1 to 269 is marked by a polypeptide chain designated as Shikimate dehydrogenase (NADP(+)). Through molecular interaction, the residue at 89 becomes linked with shikimate. Pertinent to biological process, the GO qualifier for this protein captures aromatic amino acid family biosynthetic process. The GO term of this protein covers amino acid biosynthetic process when considering biological process. In terms of molecular affinity, the shikimate is targeted by the residue at 64. Under the scientific classification, the source organism for this protein aligns with Neisseriaceae. In the interaction between molecules, the residue at 17 to 19 attaches to shikimate. A bond is formed between the residue at location 154 to 159 and the NADP(+). Member of the shikimate dehydrogenase family.

**Example Description (from SwissProt-Aug for *P95368*)**

This protein aligns with the shikimate dehydrogenase family. The sequence similarities suggest that it shares a common evolutionary history and structural or functional characteristics with other members of this family. The shikimate dehydrogenase family is known for its role in the metabolic pathways involving the shikimate pathway, which is critical for the biosynthesis of various compounds, including aromatic amino acids and other secondary metabolites.

**Example Description (from SwissProt-LLM for *P95368*)**

This protein is engaged in a critical enzymatic operation within the metabolic pathway of the organism it is associated with. Specifically, it functions as a shikimate dehydrogenase, which is a part of the shikimate dehydrogenase family. Its primary enzymatic operation is the catalysis of the reversible NADPH-linked reduction of 3-dehydroshikimate (DHSA) to produce shikimate (SA).

This reaction is a key step in the biosynthesis of aromatic amino acids, which are essential components in various biological processes. The protein's activity is linked to the chorismate biosynthetic process, which is a pivotal step in the production of aromatic amino acids, including tryptophan, phenylalanine, and tyrosine. The chorismate itself is a precursor in the synthesis of these amino acids, as well as in the production of other important molecules like the pigment chlorophyll.

The catalytic activity of this protein is also dependent on the binding of NADP+, which is a coenzyme that plays a crucial role in redox reactions. The reaction involves the conversion of NADP(+) to NADPH, which is then utilized in subsequent metabolic pathways. This operation is significant for maintaining the cellular redox balance, which is vital for the proper functioning of the cell.

In summary, the protein's enzymatic operation is the reversible reduction of 3-dehydroshikimate using NADPH, a process that is integral to the synthesis of aromatic amino acids and other key metabolites in the organism.

**Example Description (from TrEMBL50-ProTrek for *A0A1F3YBL3*)**

On the matter of biological process, the GO term specific to this protein involves bacteriocin immunity.

**Example Description (from TrEMBL50-ProLLM for *A0A1F3YBL3*)**

This protein exhibits a multifaceted identity within various biological contexts. Initially, it is referred to as B-type flagellar protein FliS, which suggests its involvement in the assembly and function of bacterial flagella, a crucial appendage for motility. The flagellum is a complex structure that allows bacteria to move through liquid environments, and the B-type flagellum is distinct from the A-type flagellum, which typically forms a cilium in eukaryotic cells.

Additionally, this protein has been identified under different names that point to its roles in various biological processes. It is recognized as Immunity protein HalI, which implies a potential role in bacterial defense mechanisms against phages or toxins. As Flagellar secretion chaperone FliS, it suggests a function in the proper folding and secretion of flagellar components.

In the context of the flagellar hook-basal body complex, the protein is termed Flagellar hook-basal body complex protein FliE. This suggests that the protein is part of a complex involved in the structural integrity and function of the bacterial flagellum. The protein is also associated with Uncharacterized protein YfhL and Putative protein RIG, which indicates that while its function is known to some extent, there are still aspects of its role that remain to be fully understood.

Furthermore, the protein is identified as Uncharacterized protein aq_aa15 and Cloacin immunity protein, which could mean that it is a part of a more specialized system within the Cloacin genus of bacteria or that its function is not fully elucidated in the broader scientific community.

The recurring mention of the polypeptide chain spanning from position 1 to 110 in various contexts suggests that this region is conserved and may be pivotal in the protein's structure and function, regardless of its various designations. This sequence is associated with different roles, from flagellar assembly to immune responses, indicating that this protein is a versatile player in bacterial physiology.

**Example Description (from UniProtKB for *A0A0A1H0A8*)**

Organic hydroperoxide resistance protein family.

**Supplementary Table 5**: **LLM prompt for data augmentation.**

| Prompt for generating protein descriptions for the SwissProt-Annot dataset |
|---|
| You are an expert biologist who is good at paraphrasing sentences from biological area while keeping their semantic meaning the same. Now given the sentence quoted by [], you have to try your best to create as much as possible variants of the sentence. Remind these sentences should have the same meaning. Here's the original sentence: [sentence]. Please give me as much as 10 variants of the sentence with the same meaning. Note that you should keep variants diverse as possible as you can. Here's the format:<br>1. paraphrased sentence 1<br>2. paraphrased sentence 2<br>... |
| **Prompt for generating protein descriptions for the SwissProt-Aug dataset** |
| You are an AI biology assistant capable of analyzing a single protein. You are provided with several sentences that describe the same protein you are examining. Using the information provided, you should analyze the protein and answer any related questions in detail. Rather than directly referencing the site information, use it to conduct a thorough analysis of the protein in natural language, discussing aspects such as the functions of specific sites, their relationships to other proteins, and their contributions to the overall functions of the protein. When incorporating details from the provided information, focus on analyzing the protein without explicitly stating the sources of your information. Always refer to the protein as "this protein" and maintain an analytical perspective throughout your response. If a question cannot be answered based on the available details, do not reference the source of the information. Here are the relevant sentences about the protein.<br>{Descriptions from SwissProt-Annot}<br>{Question} |

**Supplementary Table 6**: Sequence identity between experimentally validated active Pinal-designed proteins and their closest matches in Swiss-Prot and UniRef100.

| Sequence ID | Closest Swiss-Prot ID | Seq. Identity to Swiss-Prot | Closest UniRef100 ID | Seq. Identity to UniRef100 |
|---|---|---|---|---|
| pinalGFP | Q8ISF8 | 30.8% | UPI0021506B9F | 33.6% |
| pinalGFP-g | P42212 | 27.0% | UPI000FF8750E | 31.1% |
| pinalADH01 | P48814 | 54.5% | UPI0008115CC9 | 55.2% |
| pinalADH02 | F8DVL8 | 57.4% | A0A3T0D359 | 62.9% |
| pinalADH03 | P0DMP5 | 47.4% | A0A437RFF1 | 49.8% |
| pinalADH04 | Q96XE0 | 49.0% | A0A8T5A9C9 | 51.0% |
| pinalADH05 | F8DVL8 | 56.4% | A0A3T0D359 | 63.4% |
| pinalPETase36 | E9LVH9 | 60.2% | A0A2R7SV75 | 62.6% |
| pinalH02 | P39726 | 50.6% | G3B5R3 | 54.9% |
| pinalH04 | P39726 | 50.6% | P39726 | 55.8% |
| pinalH07 | O67573 | 55.8% | A0A3M2GYW8 | 58.2% |
| pinalH08 | Q89I87 | 54.4% | A0A931HCK5 | 55.5% |
| pinalH12 | O67573 | 55.4% | A0A7C5QLU9 | 57.9% |

Note: Closest Swiss-Prot homologs were identified by NCBI BLAST, whereas closest UniRef100 homologs were identified using MMseqs2 (GPU-enabled build, commit 1668032) against the UniRef100 database (release 25_01).

**Supplementary Table 7**: Protein amino acid sequences and associated descriptions.

| ID | Sequence | Text Prompt |
|---|---|---|
| index_1286_0 | MSDFFGPPEEELGGPKNPELKAAIAALNANPNEENEQALLEALKDARFLVPVTFDKELEEEDGKVVLDEDTKLSFLLLQNPDGEKYLPAFTSWEELKAWDPDGKYRPVVATFEDLAKLLEKDPSIAGIALNPFGDGFVLPREALEAILNGKPWVEEEEEVEIQVGEPKQPPEEFIEALREFLDQRPEIKAAYLKLMVQDGEESLLVVLDGEGDLQPLFEELGEAVKPAAGPLRLDILPADSPLAQQLTGNEEPFYKRKK | This protein is multifaceted, as indicated by various names associated with its function within the bacterial Type VI secretion system (T6SS). It is referred to as Protein SseB, which suggests its role in the secretion system's effector complex. The protein is also identified as TssB1, TssE1, TssA1, TssK1, TssC1, and TssF1, each of these names pointing to its involvement in different components of the T6SS. The T6SS is a complex machinery that bacteria use to deliver effectors into host cells, which can manipulate host cell functions to aid in the bacterial infection process. The common thread among these designations is the protein's contribution to the T6SS, a key virulence factor for many pathogenic bacteria. Specifically, TssB1 and SseB are related to the sheath component of the secretion system, which is crucial for the assembly and function of the secretion needle and baseplate. TssE1, TssA1, TssK1, TssC1, and TssF1 indicate different roles within the system, including the baseplate, effector translocation, and possibly other regulatory functions. The protein's primary function appears to be in the delivery of effector proteins into host cells, which can alter host cell physiology to promote bacterial survival. This role is essential for the bacteria's ability to establish infection and evade the immune response. The requirement of SseB for the correct localization of other components, such as SseC and SseD, underscores its importance in the assembly and functioning of the T6SS at the bacterial cell surface. |
| index_422_3 | MAKGKSLQIFLVDGSPDGIRTAEISNRTGKALVFPRSKLSKLKGREEFKQSGVYFLFGKDEDGEKLLYIGEAEAYKNGKGVLKRIRQHLKNDPELKGFDEVAIITSSDNNLNKASIKYLEYRFYELAKEAGRYKIRNGNLPKKPKLSEEDQEEMEEFLEDLKIILPGLGYDIFDPKEEEESPESEEEEEEEEE | In the context of the described protein, we can identify several modular domains, each with its specific location and category: 1. **GIY-YIG Domain**: This domain is located at amino acid positions 103 to 152. The GIY-YIG domain is a type of zinc-binding domain that is found in various nucleic acid binding proteins. It is known for its role in recognizing and binding to DNA or RNA. The protein in question contains this domain, which suggests a function related to nucleic acid manipulation or recognition. 2. **Uncharacterized Protein YBL059W Chain**: The polypeptide chain from position 1 to 193 is recognized as Uncharacterized protein YBL059W. This chain does not belong to a well-defined domain family and, as the name suggests, its function is not well-characterized. It may interact with other proteins or domains within the protein, or it could be part of a larger functional complex. 3. **Bacterial GIY-YIG-like Domain**: The protein also contains a bacterial GIY-YIG-like domain, which further emphasizes its potential role in nucleic acid interactions. This domain is typically found in proteins involved in DNA metabolism, such as those that participate in DNA repair or recombination. Regarding the category of each domain: - The GIY-YIG domain and the bacterial GIY-YIG-like domain are both zinc-binding domains. These domains are characterized by the presence of a zinc ion coordinated by specific amino acids, which is critical for their structure and function. The protein's domains are integral to its proposed function as a terminase, which is an enzyme involved in the replication of DNA and RNA viruses. The terminase is responsible for adding a poly(A) tail to the 3' end of the viral genome, a crucial step in the replication process. The presence of the GIY-YIG domain and the potential zinc-binding properties suggest that this protein may be involved in nucleic acid processing, possibly in a viral replication cycle, although the exact role and the nature of its interactions with other proteins would require further study. |

| ID | Sequence | Text Prompt |
|---|---|---|
| index_678_2 | MSSIIKPFRALLPAKGLASKVASPPYDVIDEEEREELLAKSPYNFLRIDLPELELEENPENFDPEENPPFPEPKENLEELIEEGILVRDPKPAFYVYQQERNGRTQTGIIGCVSVDAYDSGRIKKHEKTLPEKEEDRVELIEALEANFGPIFLTYPEDPEIDDLIDEITKGKPLIDFTDEDGVRHRLWRIDDPEEIEELQELFNKIPAIYIADGHHRAAAAANVRDERRAENPDYPEDADYDYFLAVLVPEEQLPPLYYYHH | This protein, designated as ORF187 and also referred to as the Immunity factor for TNT homolog and Toxin RelE, serves a critical role in the immunity response of the organism Borrelia bissettiae. It is an integral component of an LXG toxin-immunity module, which is a system designed to neutralize the toxic effects of specific toxins produced by the same strain. Primarily, the function of this protein is to counteract the RNase activity of a cognate toxin, BC_0920, thereby preventing damage to the bacterial cell. This neutralization activity is particularly significant in the context of bacterial biofilm formation, as LXG toxins can contribute to the disruption of biofilm structure and function. By neutralizing these toxins, the immunity protein helps to maintain the integrity and stability of the biofilm. The immunity protein is part of a larger system involving six LXG toxin-immunity modules in this strain. These modules are involved in promoting kin selection, mediating competition within biofilms, and driving spatial segregation of different bacterial strains. This suggests that LXG toxins may help to prevent internal conflict among strains, thereby avoiding a form of warfare within the biofilm community. This protein's function is also linked to intercellular competition. When the operon encoding the toxin is disrupted, it can be disadvantageous to the bacteria. However, overexpression of the cognate immunity protein can restore growth in competition with wild-type strains, indicating its role in competitive defense. The protein's domain, characterized as COMM and spanning positions 112 to 186, is crucial for its function. It is involved in mediating the interaction with the cognate toxin YobL, which is neutralized upon expression in E.coli. Importantly, this protein does not exhibit immunity protein activity against other LXG toxins, suggesting that its protective function is highly specific to the cognate toxin. Furthermore, the protein is likely to interact with the cognate toxin YwqJ, inhibiting its toxic activity. This interaction is key to the overall function of the LXG toxin-immunity module, as it helps to maintain the balance between toxin production and immunity within the bacterial population. In summary, this protein is a pivotal player in the immunity response of Borrelia bissettiae, providing protection against specific toxins and contributing to the bacterial competitive strategy in biofilm environments. |
| index_1192_3 | MSDKKEFAKEFYERLKEEADEEVEVSFDEEEYTLSIGDKTIGLSNLAEQLEEAPEEEHEEIIEQYLEMIEKAIGLEEEEEEFDLEKDIYPVIRSKSFVEEKEYISEPHAGDLVVTYAIDTEETYRFLTEEEFEKLPLTKEELKEIALENLKKKAPLEYEKQDIGEGETLYFLIGDGYAASRILLPEFLEEIEEEAGTELVVFVPHQDVLLVYGYKDEETLEEMAEMTLEMFEEGGIPITDLSFTYEDGELEPVFIFKKEKEEEE | This protein, which is a component of an LXG toxin-immunity module, serves a crucial role in the bacterial immune system by providing immunity against specific toxins. Its primary functions include: 1. **Neutralization of Cognate Toxins**: The protein effectively neutralizes the toxic effects of cognate toxins, such as YobL and YqcG, by directly interacting with them. This interaction is specific to the cognate toxin, meaning that the protein does not confer immunity against non-cognate toxins, which is a key feature of the immune system's specificity. 2. **Kin Selection and Biofilm Competition**: The protein contributes to kin selection, a process where bacteria preferentially compete with close relatives over more distant ones. This function is significant in biofilm formation, where it mediates competition between strains and promotes spatial segregation, which can help avoid intra-strain warfare and the resultant release of virulent toxins. 3. **Intercellular Competition**: The immunity protein is involved in intercellular competition during biofilm formation. The disruption of the operon encoding this protein can put the bacteria at a disadvantage in such competitions, while overexpression of the cognate immunity protein can restore growth in competition with wild-type strains. 4. **Immunity to Specific Toxins**: The protein imparts immunity to certain naturally sensitive host strains against toxins such as pediocin PA-1/ACH, suggesting its role in providing protection against a range of threats. 5. **Prevention of Early Activation of Toxins**: The protein also has a function in preventing the early activation of toxins like Tas1, which could otherwise lead to harmful effects. In summary, this protein is a critical part of the bacterial immune response, ensuring that the organism is protected against specific toxins, maintains competitive advantages within biofilm environments, and contributes to the overall health and survival of the bacterial community. |

| ID | Sequence | Text Prompt |
|---|---|---|
| index_827_2 | MKKKIPTYKINDFPGKEGNQDIEVSSLEEHLKLAPHIEEPHRHDFYLILYVTKGSGTHTIDFETYPIKPGSLCFVRPGQVHSWEFSDDLEGTVIIFTEDFLSKYFPNFSIEDFSFFNNSDSSPVLQIPEEDLPEFESLFEQLEEEYKSNDPYKDEILASLLYQLLLKINRLYSSQSNISNNSLELVRQFKQLVEENFKTEKQVSFYADKLNITVKTLNEITKKITGKTPSQLIQDRIILEAKRLLYYSDLSIAEIAYLLNFNDSSHFSKKFKKTGGMSPKERRKKLK | This protein exhibits several DNA-binding domains, which are crucial for its interaction with DNA and, by extension, its regulatory functions. The primary DNA-binding domain within this protein is characterized by the presence of the h-T-H motif, a common structural feature in many DNA-binding proteins. Specifically, the h-T-H motif is found in several overlapping regions throughout the protein, suggesting that the domain may be quite stable and possibly engage in multiple DNA-binding events simultaneously. The following are the placements of these h-T-H motif-containing domains: 1. From position 204 to 225, this domain is identified, which is consistent with the descriptions provided. 2. The domain is also noted to be present from position 203 to 224, which slightly overlaps with the previous span but maintains the core motif. 3. Another instance of the h-T-H motif is located from position 201 to 222, with a minor overlap in the amino acid indices. 4. At positions 251 to 274, another region of the protein is marked as having the h-T-H motif, which is again slightly overlapping with the previously mentioned domains. 5. The domain is also described from position 252 to 275, with a very slight shift in the end position. 6. At position 200 to 221, there's another mention of the h-T-H motif, which overlaps with the regions described earlier. 7. Lastly, the domain from position 249 to 272 is also noted, showing a broad consistency with the earlier mentioned domains. In addition to the h-T-H motif, the protein is characterized by a domain from position 187 to 285, described as being of the HTH araC/xylS-type. This type of domain is often found in regulatory proteins and is known for its ability to bind to specific DNA sequences. The similarity with the araC/xylS-type suggests that this protein might be involved in regulatory processes, such as gene expression control. The consistency in the description of the DNA-binding domains across the various spans indicates that these domains are likely to be functionally important, and their presence in multiple, overlapping regions could contribute to the protein's robust DNA-binding capabilities. This could be significant for processes such as transcriptional regulation, where precise and multiple interactions with DNA are often required. |
| index_3513_3 | MKSLKELLNLSEEELKETKVHIISDPENNPLLKAYLNGDLEKYEKLWKKRNAFGDKKYILVFLKEKGSNEYTFLGLYEIISSEKTENGYKYEVELLDEFEELEGRLIVEYKPPRRSTYRNLEKYKDKLIVEGILEEPIKLPEFPGYEKLVLSYDQLKEIIENNNESWKNALSSVKGIYLITDTKTGKQYIGSAYGGNGLWGRWKNYVKDGHGGNKKLKKLIEKNPEYKKNFRFSILEVAPDNATEEDILKREKEYYKLLLTKKKGLNLN | This protein, designated as ORF319, is a probable endonuclease, with a significant role in intron homing. To deduce the specific cofactor that aids this protein in its operational duties, we can consider the broader context of endonucleases and their functions. Endonucleases are enzymes that cleave DNA or RNA within the sequence of the molecule, which is a critical step in various cellular processes such as DNA repair, recombination, and gene regulation. Given the role of ORF319 in intron homing, which is a process involving the movement of introns within the genome, the cofactor likely plays a crucial role in facilitating this movement. For proteins involved in nucleic acid processing, a common cofactor is magnesium ions (Mg2+). Magnesium ions are known to stabilize the active site of many endonucleases and are essential for their catalytic activity. They can help in orienting the nucleotides correctly for cleavage and can also modulate the enzyme's affinity for its substrate. Another potential cofactor could be a metal ion like zinc (Zn2+), which is also known to be involved in the activity of certain endonucleases. Zinc ions can participate in stabilizing the active site of the enzyme and in the catalytic reaction itself. Since ORF319 is similar to endonucleases of group I introns of fungi and phage, it is possible that it also relies on a cofactor that is typical for these enzymes. For instance, some group I intron endonucleases require divalent cations, such as Mn2+ or Mg2+, to activate the enzyme. Given that the protein is encoded within intron 1 of COX1 and is involved in mobile group II introns, it's also plausible that the protein could have additional cofactors or cofactor-binding motifs that are specific to its intron-encoded nature or its role in intron movement. To definitively identify the particular cofactor that aids ORF319 in its operational duties, experimental analysis would be required to determine the conditions under which the protein exhibits its endonuclease activity. This might involve testing the activity in the absence or presence of various potential cofactors and monitoring the effects on the enzyme's function. |
| index_201_0 | EGWTVKTLILNSATYRQSSKVNPEKLEIDPENRLLARQNRRRLEAEMIRDQALAVSGLLSPKIGGPSVKPYQPEGIWENSNSKAKWKQSKGEDLYRRGLYTFWKRTFPYPSMLAFDAPDRNVCCVRRERTNTPLQALVLLNDPTFVEAARALAERILKEAKTDEERIEYAFRLALSRLPSAEELAVLLQSLQQQLAQYQANPEAAEALLAVGETKRPADLDPVELAAWTNVANVLLNLDEFITKE | The gene responsible for synthesizing the protein in question is identified by the ORF Name WH7805_09909. This gene is one of several that have been associated with the protein synthesis process, as indicated by the other gene names provided: EN45_078720, EN45_076310, NAS141_03721, NIES39_K04640, UT29_C0001G0139, NIES39_K04650, and DVH24_017674. However, the specific gene named WH7805_09909 is highlighted as the primary identifier for the gene involved in the synthesis of this protein. |

| ID | Sequence | Text Prompt |
|---|---|---|
| index_558_4 | MSELSPLERLLERLRREREEAEARGKEYKPTWGFVIYRTVYPEDDEKWEEFLELLKEETEEELRSSPGLEWLLDYLEWTVIEDPEKLDGASKEQVREHFREWVAEELNSPSGPPPSEGLDPDSSPRYRVCLQIDEECLDHFLSNLEDGKLKPDADPFVYAVEADWDEEEDENADEDEPYVGWMKVPIRDLFDELYLNLGFEDFEDLREDADEDGVYRPDGDD | The gene responsible for coding this protein is identified through various ORF (Open Reading Frame) names. These names include TRAVEDRAFT_139011, AA0117_g3624, VD0003_g7577, AA0117_g5322, BDW47DRAFT_120076, BDW47DRAFT_120077, and BDW47DRAFT_134043. These ORF names suggest that the gene could be found in different organisms and strains, as indicated by the associated organisms' names such as Galerina marginata (strain CBS 339.88), Ceriporiopsis subvermispora (strain B), and Ustilaginoidea virens. The diversity in the ORF names and associated organisms suggests that this gene may have been identified across different studies and genetic databases, potentially highlighting its importance in various biological contexts. |
| index_1157_3 | MKKKLLLAALLLLAASAAQAHEFWLEPEQFQVSASGSATLQLRVGEETEHQPFPLRPERIASFSVVGPDGKKTLPAEQLAGSIPLGQPGLHVVSLETTPRPITLEAEKFNEYLKEEGLTDVLAARKAAGKAAQPAKEQRYAKALVGVGDGKGADKPLGLPIEIVALANPYGADLSDGLTVQVLYDGKPLANVKVTVFEKSADGWVKVSTVKTDANGRATLPVEPGHRYLLSAVKMRPAADPAKADWLSYFASLTFFAPK | In the context of the mature form of the protein, the polypeptide chain corresponds to the region following the removal of the propeptide. Specifically, the mature protein's polypeptide chain is recognized as the Putative protein-disulfide oxidoreductase RP025, spanning from position 22 to 272. This region is of particular interest as it is where the protein's functional activities are likely centered, given its classification as a protein-disulfide oxidoreductase. This area is responsible for the protein's disulfide bond formation and reduction, which are critical in many biological processes, including protein folding and stability. The mature form does not include the propeptide, which is situated between positions 247 to 272, suggesting that this segment is not involved in the protein's mature function and is shed during the maturation process. |
| index_1133_0 | MWLLNTETLELKEFDDPESVPYAILSHTWGDEEVTFQDMINGTNKDSAGYKKIKKFCELARERYGIEWIWIDTCCIDKSSSAELSEAINSMFRWYQLAEVCIVYLSDVPAPSDPSDLDSPEFEEAFRNSRWFTRGWTLQELIAPRSVEFFDKDWQLIGTKRSLADLISEITGIDADVLRGEAPLSDFSVAERMSWAANRETTREEDRAYCLLGIFDVNMPLIYGEGDKAFRRLQEEIMKRIPDDSIFAW | The gene responsible for coding this protein has been identified through various ORF (Open Reading Frame) names. These names are VFPFJ_04709, VFPFJ_04705, VFPFJ_04707, VFPFJ_04702, VFPFJ_04704, and VFPFJ_04701. These ORF names suggest that the gene in question is likely part of a larger genomic context within the organism being studied. Each ORF name might correspond to a specific region or variant of the gene, potentially reflecting different alleles or isoforms of the protein. The use of these distinct ORF names may indicate that the gene has multiple transcripts or that it has been studied in various genetic backgrounds or conditions. |
| index_672_1 | MRLLNTRTLELEEFLDDDIPPYAILSHTWEDEEVSFQDIHPEEKRKKKKGYAKIKRCCELALKDGLEYVWVDTCCIDKTSSAELSEAINSMYRWYQKAEVCYAYLSDVSSDDDPETVLPKCRWFTRGWTLQELIAPSSVEFFNKDWEEIGTKDSLADLISEITGIPIDVLRHEKPLSSVSVAERMSWAANRETTRVEDRAYCLLGIFDVNMPTIYGEGENAFRRLQEEIIKRSDDQSIFAWGDPSSPSDLSSSPDDLLLL | This protein, which is part of the Heterokaryon incompatibility protein family, is associated with a complex biochemical process that is pivotal in the organism's biology. Specifically, the protein contains a domain that is annotated as a glycosyltransferase domain, which is characteristic of enzymes that catalyze the transfer of glycosyl groups from a donor to an acceptor molecule. The glycosyltransferase activity in this protein suggests that it may be involved in glycosylation processes, where sugars are added to other molecules, such as proteins or lipids. This modification can significantly impact the structure and function of the glycoprotein or glycolipid, potentially affecting its stability, solubility, and interaction with other proteins or cellular components. Given the context provided, this protein's catalytic function is likely related to the synthesis or modification of certain toxins. The mention of mutated toxins being expressed as transgenes in yeasts that cause lethality in lepidopteran larvae suggests that the glycosyltransferase activity may be part of a pathway that modifies or synthesizes these toxins. This modification could be crucial for the toxin's efficacy, allowing it to evade the host's immune system or enhance its ability to disrupt cellular processes in the larvae. Furthermore, the protein's impairment of pathogenicity on wheat lines with the Snn1 gene implies a role in interactions with the plant's defense mechanisms or perhaps in the protein-protein interactions that are essential for the pathogen's virulence. The glycosyltransferase domain may be involved in these interactions by modulating the protein's structure to facilitate recognition or binding to specific receptors or targets. The down-regulation of this protein's expression during fruiting body formation suggests that its function may be particularly relevant during certain stages of the organism's lifecycle, possibly indicating that glycosylation or toxin modification is a critical aspect of the organism's reproduction or environmental adaptation. In summary, the biochemical process that this protein likely catalyzes involves the glycosylation of molecules, potentially modifying toxins to enhance their biological activity and contributing to the organism's ability to interact with its environment, either as a pathogen or in other ecological roles. |

| ID | Sequence | Text Prompt |
|---|---|---|
| index_934_0 | MRLLNTETLELEEFLGDIPPYAILSHTWEDGEEVTFQDLQEGKGESKSGYAKIKRCCELARKDGYEYVWIDTCCIDKSSSAELSEAINSMFRWYQEAEVCYAYLSDVPSIPISEENEEELEAAFRKSRWFTRGWTLQELLAPRSVEFFSKDWQLLGTKSSLADLISEITGIPVEALRGSPLSSFSVAERMSWAASRETTRVEDMAYCLLGIFDVNMPLIYGEGENAFRRLQEEIMKSSDDHSIFAW | The protein in question originates from the organism Fusarium acaciae-mearnsii. This species is a fungal pathogen that has been identified in association with the protein sequence. |
| index_2830_4 | MLLLTKKELEELLEESNDLPDWQKEAFEWFKKKMTNKTNPFPCIFGKKAYKKDQLRYAFAEQGDPDAAEEIAEALPLLPEISPYSSLVVFFEDLGEDATIEDYEQLFWDTLKEVHALDPVPWPEDIPKDPEHPLWEFSFNGEPYFIIGMTPAHKKRKSRRFPYLMLVFNPRSQFEYLREKGRYEKTKKKIRKRLDKLDSVPLPPLLGSYGSKENSEWKQYFLPDEDKPCPFK | In analyzing this protein, several potential errors or areas of confusion can be identified: 1. **Protein Length Ambiguity**: The protein, referred to as Protein YecM, is described to have a polypeptide chain spanning positions 1 to 188. However, it is also noted that this protein may be missing up to 130 C-terminal residues compared to its orthologs. This discrepancy raises questions about the true length of the protein and how this affects its structure and function. It could lead to confusion regarding the complete sequence and the role of the truncated portion of the protein. 2. **Original Misclassification**: The protein was initially thought to encode for OrgA and OrgB, which suggests that its function and role within the cellular context were initially misunderstood. This misclassification could have implications for how the protein is studied and its current classification as a putative immunity protein component. 3. **Functionality of C-Terminal Region**: The protein's function is linked to blocking the toxic effects of the C-terminus of cognate toxin RhsC. The fact that this protein is missing up to 130 C-terminal residues could potentially affect its ability to perform this function, as the C-terminus is crucial for toxin interaction and neutralization. 4. **Sequence Similarities**: The protein shares sequence similarities with E.coli YfiI and P.aeruginosa RluD. This raises questions about the conservation of function across different organisms and the significance of these similarities in the context of the protein's role in a toxin-immunity protein module. 5. **Interactions with Other Proteins**: The protein interacts with YeeF but not with YobL, which might be due to the differences in their C-termini. The interaction with YeeF is thought to inhibit the toxic activity of the toxin. The stoichiometry of this interaction (2:2) suggests a precise requirement that could be difficult to achieve, leading to potential confusion in experimental designs and interpretations. 6. **Gene Name and Protein Synthesis**: The nucleotide sequence yubF corresponds to the synthesis of this protein. However, without a clear understanding of the complete protein sequence and its structure, the relationship between the gene sequence and the protein product might be difficult to interpret fully. 7. **No Obvious Phenotype on Deletion**: The deletion of the yeeF-yezG operon does not lead to a visible growth phenotype. This could be confusing, as it suggests that the function of this protein might not be essential for growth under certain conditions, or it could be that its role is masked by other cellular processes. Overall, these ambiguities and potential errors highlight the importance of further research to clarify the protein's structure, function, and interactions within its cellular context. |
| index_1157_2 | MKKLLLLLLLLLLLSSAAAQELNCQVQVNAPQVNADREVLRSLQKAITDFLNTTKWTDEEFEEEERIQCNILINLTEQPSENTFKGTLQVQSSRPVYGTNYSTPLLNFRDKNFEFEYIEFQPLEFNPNTFVSNLTSILAFYAYLILGLDYDSFSLNGAKPYFEKAKNLVNNAQNAGAPGWSDSNVKVNRYWLIENLLNPNFADVREAMYDYHREGLDRFSEDKEKAKKAIKEAIPKLKKAKRARPGSYLLKALLSSKEEELRRIGGKKNINN | In the context of the mature form of the protein, the polypeptide chain corresponds to the region following the removal of the propeptide. Specifically, the mature protein's polypeptide chain is recognized as the Putative protein-disulfide oxidoreductase RP025, spanning from position 22 to 272. This region is of particular interest as it is where the protein's functional activities are likely centered, given its classification as a protein-disulfide oxidoreductase. This area is responsible for the protein's disulfide bond formation and reduction, which are critical in many biological processes, including protein folding and stability. The mature form does not include the propeptide, which is situated between positions 247 to 272, suggesting that this segment is not involved in the protein's mature function and is shed during the maturation process. |

| ID | Sequence | Text Prompt |
| --- | --- | --- |
| index_2181_0 | MRLLNTRTLELEEFFGDDIPPYAILSHTWGDEEVTFQDMINGTAESKKGYAKIQECCELAREDGYEYVWVDTCCIDKSSSAELSEAINSMYRWYQKAEVCYAYLSDVPSDDDPRAEGSLFRKSRWFTRGWTLQELIAPSSVEFFSQEWEEIGTKRSLEQLISEITGIDIDVLRHERSLSDFSVAERMSWAANRETTRVEDRAYCLLGIFDVNMPLIYGEGDNAFRRLQEEIMKKSDDHSIFAW | The protein in question is associated with multiple gene names, which suggests that it may be encoded by more than one gene. The gene names MAJ provided include_08936, MAJ_07524, MAJ_07522, UVI_02036180, MAJ_09462, and MAJ_07523. Additionally, there are two similar gene names, MAJ_09464, which is likely a typographical variation of MAJ_09462. It is common for proteins, especially those involved in complex biological processes such as heterokaryon incompatibility, to be encoded by multiple genes, which might contribute to different variants or isoforms of the protein with distinct functions or roles within the cell. In the context of the information provided, it is not possible to determine a single specific gene name that uniquely codes for this protein without additional information about which gene is the primary or the most relevant for the protein's function. If we were to prioritize based on the number of mentions, MAJ_09462 (or its possible typographical variation MAJ_09464) might be considered a strong candidate, as it is mentioned twice along with its corresponding protein name, Heterokaryon incompatibility protein S. However, without further context or experimental data, we cannot definitively state that this is the gene that codes for the protein. |
| index_2536_3 | MSDPYEELREKLDAHPVPVPDTPELREILKLLFTPEEAALAAKLPFKPLTVEEAAKLTGLPEEEVEALLERMAEKGLVLRTEKDGERRYMLAPFVVGIFEFQFMMKRDLGERGKELAELFKEYLEDGEFAEFLLAGEVPMMRVVPVESAIPPDQEVLPYEKVKEHIERADKIAVAECFCRKQKELLGKDCGKPLETCMALNRGAEYYIEQGFAREITKEEALEILDEAEEAGLVHQVDNSQEELGFICNCCGCCCGLGKAALLLGAGGAMAASTFTADVDPEKCVCCGACVAKCVPGAIEFDEEKKKATVTDDCCLGCGCCVTVCPGFAITIERRDGEEEPPPPSEELLKKLIKAKRAAAA | In the final product of this protein, the polypeptide chain is formed by the sequence of amino acids that make up the entire protein structure. Specifically, the polypeptide chain extends from position 1 to 373, as indicated by the Chain description. This chain is recognized as an uncharacterized protein, which suggests that while its structure is known, its specific function or role within biological processes may not be fully understood yet. The sequence from position 1 to 373 is likely the primary structure of the protein, which then folds into a three-dimensional structure, potentially involving the domains and motifs described. For instance, the domain from position 287 to 318 is characterized as a 4Fe-4S ferredoxin-type domain, which is known to be involved in electron transfer reactions. This domain is repeated, with the motif at position 306 to 331 being classified as Cx9Cx9RCx2HK, indicating a pattern of repeats and possible functional significance. The region from position 1 to 373 also encompasses the binding site at position 320, where the residue forms a bond with the [4Fe-4S] cluster, suggesting that this site is crucial for the protein's function, possibly related to iron-sulfur cluster assembly and electron transfer, as indicated by the mutagenesis data. The mutation from cysteine to serine at position 321 affects iron-sulfur cluster assembly, which underscores the importance of this region in the protein's activity. The protein is part of the NapF family, and its gene sequence corresponds to the nucleotide sequence rnfC, which is indicative of its genetic origin and potential involvement in cellular processes related to iron-sulfur cluster metabolism or electron transport. In summary, the polypeptide chain from position 1 to 373 is the part of the protein that forms the backbone, upon which the protein's structure and function are based, including the iron-sulfur cluster interaction and its potential role in electron transfer. |
| index_672_2 | MRLLNTRTLELEEFIGDNIPPYAILSHTWEDEEVTFQDMQDPDRESKKGYAKIKRCCEQALKDGLEWVWIDTCCIDKTSSAELSEAINSMYRWYREAEVCYAYLSDVSGDGFSEEEFRKSRWFTRGWTLQELIAPRSVEFFSKDWGRIGTKRSLEEEIHEITGIPIEALRGASLSDFSVEERMSWAAKRETTREEDMAYCLLGIFDVNMPLIYGEGDKAFRRLQEEIMKSSDDDSLLSSLLELLSSSSSSSSSGPPPDDDSLLSSSSSSSS | This protein, which is part of the Heterokaryon incompatibility protein family, is associated with a complex biochemical process that is pivotal in the organism's biology. Specifically, the protein contains a domain that is annotated as a glycosyltransferase domain, which is characteristic of enzymes that catalyze the transfer of glycosyl groups from a donor to an acceptor molecule. The glycosyltransferase activity in this protein suggests that it may be involved in glycosylation processes, where sugars are added to other molecules, such as proteins or lipids. This modification can significantly impact the structure and function of the glycoprotein or glycolipid, potentially affecting its stability, solubility, and interaction with other proteins or cellular components. Given the context provided, this protein's catalytic function is likely related to the synthesis or modification of certain toxins. The mention of mutated toxins being expressed as transgenes in yeasts that cause lethality in lepidopteran larvae suggests that the glycosyltransferase activity may be part of a pathway that modifies or synthesizes these toxins. This modification could be crucial for the toxin's efficacy, allowing it to evade the host's immune system or enhance its ability to disrupt cellular processes in the larvae. Furthermore, the protein's impairment of pathogenicity on wheat lines with the Snn1 gene implies a role in interactions with the plant's defense mechanisms or perhaps in the protein-protein interactions that are essential for the pathogen's virulence. The glycosyltransferase domain may be involved in these interactions by modulating the protein's structure to facilitate recognition or binding to specific receptors or targets. The down-regulation of this protein's expression during fruiting body formation suggests that its function may be particularly relevant during certain stages of the organism's lifecycle, possibly indicating that glycosylation or toxin modification is a critical aspect of the organism's reproduction or environmental adaptation. In summary, the biochemical process that this protein likely catalyzes involves the glycosylation of molecules, potentially modifying toxins to enhance their biological activity and contributing to the organism's ability to interact with its environment, either as a pathogen or in other ecological roles. |

| ID | Sequence | Text Prompt |
|---|---|---|
| index_433_4 | MSSTSSPSPAGPVGAEIRPFRALRYNPEKIADLSDVVAPPYDVISPEEQERLYARHPYNIIRLILGKEEPDDSEENNRYTRAAELLEEWLEEGVLVQDPEPAIYVYRQEFTLDGKEVTRTGFIAAVKLEPFGTGAVLPHEQTLAKPKADRLKLIEATRANLSPIFLLYRDPERAVARILEEVTAGEPLIEVTDEDGVTHRLWRISDPADIAAIQELMADKSLLIADGHHRYETALNYREEHGDTSPGADYVLATLVNLDDPGLPIPPIHVVRLLGPEDDDELLLEELLAGLFEVEVAADEEELLAALAAQGPGAFALRFADSGTLLLLRLDDEAALAELLPDLPEALASVDDALLAALLLPLLLGDDAEELEEGGRLVYVRDEEELAALLAEGDAAFFLLNPPPIEVVVEAAGEGGPQQQKYTYFPPKRLTTLFFRPLL | The organism that serves as the source of this protein is Deinococcus erythromyxa. This organism is identified in the context of the protein sequence provided. |
| index_752_0 | MRLLNTRTLELEEFLGDPPPYAILSHTWGDEEVTFQDMQNGTYKGKAGYEKIKRCCEQALRDGLEYVWIDTCCIDKSSSAELSEAINSMYRWYRDAAVCYAYLADVPSDDDPETVLPKCRWFTRGWTLQELIAPRSVEFFNKDWELIGDKASLADLISEITGIDADVLNGEKPLSSASVAERMSWMANRETTRVEDIAYCLLGIFDVNMPLIYGEGDKAFRRLREI | This protein, which plays a crucial role in the expression of heterokaryon incompatibility and sexual functions, offers several intriguing insights regarding its structure, function, and regulation: 1. **Genetic Identity**: The protein is encoded by the gene ORF Name VFPFJ_04705, suggesting a specific genetic heritage within the organism. This genetic identity is essential for the proper expression of the protein's functions, which are integral to the organism's reproductive biology. 2. **Functional Significance**: As a component of the heterokaryon incompatibility system, this protein is involved in preventing the fusion of genetically distinct fungal hyphae. This is particularly important in fungi like the Oat take-all root rot fungus, which can lead to significant crop losses if not controlled. Its role in sexual functions suggests that it may also be involved in the mating process, which could have implications for the genetic diversity and adaptation of the fungus. 3. **Disruption Phenotype**: The disruption of this protein impairs the production of victorin or any derivative or intermediate. Victorin is a known antifungal compound, so the loss of this protein might result in increased susceptibility to fungal infections. This highlights the protein's role in defense mechanisms against pathogens. 4. **Expression Regulation**: The protein's expression is influenced by various environmental factors and regulatory proteins. It is repressed by glutamine and at alkaline pH, which could be a response to certain nutritional or environmental stresses. Conversely, it is highly induced under nitrogen starvation and acidic pH conditions, indicating its role in stress responses. The negative regulation by vel1 further suggests complex regulatory pathways that may be involved in fungal development and survival. 5. **Structural Analysis**: The protein's chain, spanning from position 1 to 680, is recognized as Heterokaryon incompatibility protein 6, OR allele. This implies that the protein has a defined structure that is crucial for its function. Analyzing this structural information could provide insights into how the protein interacts with other molecules and performs its functions. 6. **Biotechnological Implications**: The use of lichens containing usnic acid and the mention of chondramides, which are secondary metabolites with antifungal activity, suggest that the protein and its associated pathways might be of interest in biotechnology. These compounds could potentially be developed into antifungal agents, and understanding the protein's role could lead to more effective control strategies for fungal diseases. 7. **Potential for Toxicity**: The protein's link to usnic acid, which is toxic to the liver and can cause contact allergy, might suggest that the protein itself or its metabolites could have similar properties. This could be a concern in therapeutic applications or when using the protein as a model for studying fungal biology. In summary, this protein is a multifaceted molecule that has a significant impact on fungal biology, including sexual reproduction, defense against pathogens, and stress response. Its complex regulation and potential biotechnological applications underscore the importance of further study into its mechanisms of action and structure. |

| ID | Sequence | Text Prompt |
|---|---|---|
| index_3236_1 | MSAPTAPVGVRPFRGVRYDPAKVGDLSRVVAPPYDVIDPALQERLAARAPHNVVRLELGKEEPGDPPDDNRYTRAAALLAEWLADGVLVRDPEPALYVVYRQTFTVDGRTYTRRGFIAAVRLGDFAAGGAVLPHEHTLSGPKADRLALLRATRANFSPVFALYDDPGGAVAALLDELAAGPPDLEVTDDDGVRHRLWRVDDPADIAALAAAFAGRPLYIADGHHRYETALAYRDERRAETGAGADAPYDYVLAMLVNLPDPGLVLPTHHRVVRGDDDALAAAAAAAVFEVVEVADAAALAAALAAPGGGARRRALRLGDGRFVVLRLRRPGAAAAAALAGGDRPPLDALALHLGLLLALLGLSAEAAAAHLELVRDAAEAAAAAAAGGARAAAFPLPTEMDLAAAVAAAGAPLPKKKTYFYPKPLSGGLIRRL | The gene responsible for producing the described protein is identified by the ORF (Open Reading Frame) name GA0070610_0181. This gene encodes the protein in question, which is associated with a specific protein family and has a conserved domain structure. The ORF name serves as a unique identifier within the genomic context of the organism from which the protein is derived. |

**Supplementary Table 8**: Pinal prompt texts for different protein types.

| Protein Type | Prompt Text |
|---|---|
| ADH | Please design a protein that is an alcohol dehydrogenase. / alcohol dehydrogenase. / This protein is an alcohol dehydrogenase. / Generate alcohol dehydrogenases for me. / This is an alcohol dehydrogenase. |
| GFP | Green Fluorescent Protein (GFP) is a light-emitting protein originally isolated from the jellyfish Aequorea victoria. When exposed to blue or ultraviolet light, its unique structure—a beta-barrel encasing a self-formed chromophore—absorbs energy and re-emits it as bright green fluorescence. Unlike many fluorescent markers, GFP requires no external additives (e.g., substrates) to glow, making it ideal for non-invasive live-cell imaging. |
| PETase | PET hydrolases are a class of enzymes capable of catalyzing the degradation of polyethylene terephthalate (PET), a widely used plastic found in bottles and packaging. This enzyme class includes cutinases, lipases, and esterases, which hydrolyze the ester bonds in PET through a conserved catalytic triad (Ser–His–Asp/Glu). PETase is a well-characterized PET hydrolase that specifically targets and breaks down PET polymers. |
| H-protein | Design a H protein. H protein acts as an essential mobile shuttle within the glycine cleavage system, transferring the methylamine intermediate between the P and T proteins, thereby enabling glycine breakdown or reversed glycine synthesis. Simultaneously, it serves as a crucial carrier protein intermediate in the protein lipoylation pathway, facilitating the attachment of essential lipoic acid cofactors to other key metabolic enzymes. Its covalently bound lipoamide arm is central to both carrier functions. |

**Supplementary Table 9**: Amino acid sequences and structural token sequences of fluorescent proteins.

| ID | Protein Amino Acid Sequence | Protein Structure Token Sequence |
|---|---|---|
| pinalGFP | MSTKSASSSAALSAPVVKFKLTGKGSVNGIKFTATGTGEGDT VNGYAEVDLNIDGTGDLPLSPDLLALFGGTWGRLFAKVQPG TPPFLAALGPKGFTFKRTITFEDGSVVEVVQTVTFEGGTLVI EIDLKGSGFPEDSLFASGKLTKILPFFEHYFPADGKIKSRFTL AIPTIDGGYVYASVTQTYRFNEPGELPEPHVARVTLEITEKSE DRTYLVLSEKSKAIGPKDGPELLGL | DPDPPPPPVVLLADQKAKAKEWEWEDAARQTWIKIKIKMAG LAQFKMKMKMFIPPRPLDLWACQVVQCCPPVVQNSSAHEDP PFFPQQSQLPQQGKKKWKWKAWPQGKIKIKIWTWHDDPRYI YIYMYMYIYDDDCPDCTSVVFWAHWDKWKKKWADDPQWIK IWTKIWTAGNVGDTIIIIMIMIMDRPDRDDHGHIWMKIKDKDF PDADPSRSITMMMMGIYIDHPVNPCVVVVD |
| pinalGFP-g | MSAAAKLLEGTVKIQMTGSGDIGGQKFTYTATGTVNPATGT LDGTIRSEDGKLPFHPSFLIGLGTYGRRGFSAPDPDIPEALAF LDTLPAGYTFERTIRIEDGPTIESEGTVTFEGNEITIETKIEVT NIDPDDPLASGKLTPDIPPYHEQYIPDPETKALRSTRALFIPL KGGGKVAFSVEQLYTPLSDGPFPLPGTHLAHVTQSWSKVEG SEGPTLEFTESARAFSIVPPDEAKLL | DAPLLVLQADKFKEWEKEWEDAPNDTWIKIKIWIAHLQFQKT WIKMFTPVFADQWAVQVCVCVTPVLVSLSAHEDPVFVLARQ QSNLPQQGKKKWKWKDWPVAKIKTKIWTWHDDDRYIYIYM YIYIDRGDCCDCTSVVFFDNWFDKDKKKWAFDVVQLWIWIW DKGWTAGPPGDTIIMIMIMIIHRSDPDDGRGHHIWMKTKDKT WDDDPPDPGDMITIMMHIHIDDDDDPPVVVVD |
| esmGFP | MSKVEELIKPDMKMKLEMEGEVNGHKFSIEAEGEGKPYEGK QTIKAWSTTGKLPFAWDILSTSLTYGNRAFTKYPEGLEQHD FFKQSFPEGYSWERTITYEDGATVKVTADISLEDGVLINKVK FKGENFPSDGPVMQKKTTGWEASTELITPDPATGGLKGEVK MRLKLEGGGHLLADFKTTYRSKKKEKLPLPGVHYVDHRIVN EKATHPEGKEYMIQYEHAVARLA | N/A |
| esmGFP-b8 | MSKVEELIKPEMKMKLEMEGEVNGHKFSIEAEGEGKPYEGK QTIKAWSTTGKLPFAWDILSTSLTYGFRMFTKYPEGLEEHD YFKQSFPEGYSWERTITYEDGATVKVTSDISLEDGVLINKIKF KGTNFPSDGPVMQKKTTGWEPSTELITPDPATGGLKGEVK MRLKLEGGGHLLADFKTTYRSKKKEKLPLPGVHYVDHTIRN EKAPHPEGKEYVVQYETAVARLA | N/A |
| avGFP | MSKGEELFTGVVPILVELDGDVNGHKFSVSGEGEGDATYGK LTLKFICTTGKLPVPWPTLVTTLSYGVQCFSRYPDHMKQHD FFKSAMPEGYVQERTIFFKDDGNYKTRAEVKFEGDTLVNRI ELKGIDFKEDGNILGHKLEYNYNSHNVYIMADKQKNGIKVNF KIRHNIEDGSVQLADHYQQNTPIGDGPVLLPDNHYLSTQSALS KDPNEKRDHMVLLEFVTAAGITHGMDELYK | N/A |

**Supplementary Table 10**: Amino acid sequences and structural token sequences of active ADHs identified in this study.

| ID | Protein Amino Acid Sequence | Protein Structure Token Sequence |
|---|---|---|
| pinalADH01 | MSLKGKNVLFVGGLGGIGLATCRALVKKNLKNLAILDIVENP<br>EAVEELKALNPSVKVTFVKCDVTKPESIKAAFEEVKEKFGHL<br>DVLVNGAGILDDKNIEKTIAVNLTGLINTTLAALPLMDKRKK<br>GKGGVIVNVASVAGLEPFPAVPVYCASKHGVVGFTRSLGHP<br>FYYELTGVKVIAVCPGITETPLLKNLGKNPLFDEFKELVEKL<br>LKLKKQTPEECGKHIAKVIETAENGSIWKSDKGKLSELELPP<br>WWKPPKS | DQQAQFAEEEEQLPFLLSLLLLQLLLVRNHQEYEYEYCDDDVP<br>SQVVSCVSHVNYHYDYDNADLLDLVRLVVSVVVVCVPPVAGA<br>EYELDFAAQQQVCLPVRQSTLAVSLVSNCVSRLQRDFPVHRHD<br>AHEYEREAAVLLVDPDQQRRSSSVSSVNSLVVQLVCLDPVNCV<br>RGVYAYEYEHEYAEPGVRLVCSCPNTNDPVCNVVNVVSNPDD<br>HHYSNLSSNLVSVCRPDDDRSWYWYDGPNDIDTDDDDDPDDD<br>DDD |
| pinalADH02 | MNLNEFYVPGKNLFGAGSIKELAEELKGKGYKKALLVTDKF<br>LVKLGLADKLKALLAEAGIETVVFDGVEPNPKDTNVMEGLA<br>VFKENGCDFIIALGGGSPHDCAKAIALVATNGGSIRDYEGVD<br>KSKKPCLPLIAINTTAGTGSEMTRFAVITDEERHHKMVIVDK<br>HVTPKIAVNDPELMVGMPPGLTAATGMDVLTHAIEAYVSTG<br>ANPITDALALQAIRLIAKNLPKAVKNGSDLEAREQMHYAQML<br>AGMAFNNALLGVHHAAAHALGGFYNPPHGAGNVVVLVVVM<br>YYNAAVVPEKFAIIAKAAGLDVDGDSEEEAAEKAAEVVKDLL<br>KDVPIPLKLSGGGVKKEDIIPLAELAKKDACAATNPRPSSLEE<br>IIDIYEEAF | DPDFDFAWAQAEQAEAPSLLVVLVVCQPVPFAEEEEQAAPVC<br>VVVCVVVVSQVSNVVSNHHYDYDHHFDQQGALVSLVVSLVSC<br>VVRVGQAYEQAEAQRSLLSRLLNQQCNPQPDGSVCQFDEASG<br>DDAGRAYEYEYQALADLSRIAQWHWHQYPVVLATGIRRHSRS<br>GHNYYYQHLVSNLPDALVRLLLRLLNLLQLLLFLLQWPQHDPQ<br>LVVLSLLLLLLSLVQSLVCNVRSNPSVSSSSSSSSNSSVSRCSRR<br>PGALQQLLLSLCCSPVVDDSSLSSLLLVLLLLVVLCVPPLQSLLS<br>SQVSSVDPCPPPDSSVSSVVSSVSSLVSSVSSDRDNAVVVVVDD<br>LVCLLVSLVSSCVDPSHVTRNDRDDSVVSSVSNVVRD |
| pinalADH03 | MSLEGKTIVVTGASSGIGLATAKLLQERGAEVIGVDIVAPDFE<br>VAQFIQADLSTPEGVEAALAQLPEQIDGLVNNAGVGPSAPAE<br>LVLAVNLLALVALTEALLPRVPPGGSIVNTSSNAGRLWRDDP<br>EELEELLAAETPEELEAYLAANPIPKEEAYAFSKALVIALTRR<br>LALPLFRERGVRVNAVAPGLVDTPILDGFVEAMGEEAVAAL<br>LALQPRLAQPEDVANVIAFLASDESRWITGQVIFVDGGLSLLL<br>R | DACAPAEEEFEPCAWFLNVLLQVVNVVRRYQYEYEECDDHP<br>DDHNYYDHFDQLDPVRLVSSVVPDDLAGQEYELDDADFLVDD<br>LSSRCRRLALSSLSNCVVCLNRHAALGEYEHEAALLLVCLVVV<br>VPVLVVLLPDPHSVSSVVVCVVVPDDSLCSNSVSSVSLLNSQLV<br>VAPVSCVVGNYAYEYEHEYAAPIPCNVSVCVSCDVVSVVVLCV<br>VAVDHHYSNLRSVVVSVRRDPVCSVDHSYYHYSYRCSSVVVD |
| pinalADH04 | MKAAVLEEFGKPLEIVEVPKPEPKGEQVLIKVEAAGVCHSDV<br>HIWTGKYGGWDLEEDFGFKLPFTLGHEVAGVVEAVGPEVE<br>GWKVGDRVAVYPWIGCGDCRYCRSGEENLCENGRWIGLTV<br>DGGFAEYVLVPDARYVVKIENLSPEEAAPITCAGVTVYRAVK<br>EANPTSSDTVAILGAGGGLGTLAVQYAKALSDAKVIAIDIRDE<br>GLELAKEMGADVVINSATEDVVEEVKEITKGRGVDAVLDFV<br>GSPATWEKAPKLLAPGGTLTLVGVGGPPPPPPVLLAVGGGK<br>IIKGSYYGNRRDLEEVEEFVKGGVPPPPEIETIPLEEAAEGLE<br>KLLLGKIIGRYVLEP | DWFFWQADAPDGTDIDDDDQDDAAAQKFKFQFQKFWDDLQ<br>VLVVSNQDDQPDRCCPQQVADDTAGAGQLTWGFTCDYHPNH<br>PPDDGGFTKTFQQQAAPQPDPCNVVQFGSPHPNTRGDRRRH<br>HHDLGRMDMDPHPLRIDGFDQDDSHLRSVCSFLVLQLLVLLVV<br>LVAELVFEEEEELLLDQSNLSNLLNCVLPYNHQYEYEDQDPNS<br>QVSSVVSPHPHYDRVNPDVPLVVLCVVVVNQFGQEYEDQAQA<br>PVCLVRRVSRHGQNHEYEYEHSDYDDRDDDPVVCVVRVYYY<br>HYTGRGGSVSSVVSVVCDSVVSGGGKDEDEDASVCSSVQSVCN<br>VVVNDDIIYMHGD |
| pinalADH05 | MKINEFYVPGKNLFGAGSIKELAEELKGKGYKKALLVTDKF<br>LVKFGLADKLKALLAEAGIETVVFDEVEPNPKDTNVMEGLA<br>VFKENGCDFIIALGGGSPHDCAKAIALVATNGGSIRDYEGVN<br>KSKKPCLPLIAINTTAGTGSEMTRFAVITDEERHHKMVIVDK<br>HVTPTIAVVDPELMVGMPPGLTAATGMDVLTHAIEAYVSTG<br>ASPITDALALQAIRLIAKNLPKAVKNGSDLEAREQMHYAQML<br>AGMAFNNALLGVHHAAQHQLGGFYNPPHGAGNVVVLVVVM<br>YYNAAVVPEKFAIIAKAAGLDVDGDSEEEAAEKAAEVVKDLL<br>KDVPIPLKLSGGGVKKEDIIPLAENAKKDACAATNPRPSSLEE<br>IIDIYEEAY | DDDFDFAWAQAEQAEAPSLLVVLVVCQPVPFAEEEEQAAPVC<br>VVVCVVVVSQVSNVVSNHHYDYDHHFDQQGALVSLVVSLVSC<br>VVRVGQAYEQAEAQRSLLSRLLNQQCNPQPDGSVCQFDEASG<br>DDAGRAYEYEYQALADLSRIAQWHWHQYPVVLATGIRRHSRS<br>GHNYYYQHLVSNLPDALVRLLLRLLNLLQLLLFLLQWPQHDPQ<br>LVVLSLLLLLLSLVQSLVCNVRSNPSVSSSSSSSSNSSVSRCSRR<br>PGALQQLLLSLCCSPVVDDSSLSSLLLVLLLLVVLCVPPLQSLLS<br>SQVSSVDPCPPPDSSVSSVVSSVSSLVSSVSSDRDNAVVVVVDD<br>LVCLLVSLVSSCVDPSHVTRNDRDDSVVSSVSNVVRD |

**Supplementary Table 11**: Amino acid sequences and structural token sequences of active PETases identified in this study.

| ID | Protein Amino Acid Sequence | Protein Structure Token Sequence | T$_m$ ( °C) |
|---|---|---|---|
| pinalPETase36 | MAITTGAGPAPTAATLQASSGPYSVTSVSVSSTGAS<br>GFGGGTIYYPTASGSFGAVAISPGFTGSQSSIAWLAR<br>HLASHGFVVVAINTNSTLDSPQNRGNQLISALDYLT<br>NSAPAAVTAKIDTGRLGVMGHSMGGGGTLIAATQR<br>PDLKAAIPLTPWNLTTNFSGIKAPTLIIGAQNDTIAP<br>VALHSRPFYNSLPSDVPKAYIEITGATHFTPTFPNPT<br>ISRYSVAWLKRYVDGDARYDSYLCPGPTPGTNPNIS<br>RYKTNNPP | DDQPLDDDDQDDLVQLLDLAHQFDKDKDWFPCVL<br>QDLFHTFIKIAGPDQDAAAEEEEEEAQLDALLLQVS<br>LQNRLRRNGYTYTRTHGPDSHDALCSLLSSRVSVLV<br>CQCPPDDPSSNSRYDNQQYAYEYAASNLSSQLNNQL<br>VCVSYQEGERELYHDPDLASCSGQHQYEYEHECAE<br>PPSHCVNHVVSNQVNHDLPHWYKYWYFAPDYSSPS<br>SDDDSLCSSVVNLSSCCGSSVNCSSQVCVVVHDDAP<br>PHRGTNDMDTSPPD | 56.84 ± 0.29 |

**Supplementary Table 12**: Amino acid sequences and structural token sequences of active H-proteins identified in this study.

| ID | Protein Amino Acid Sequence | Protein Structure Token Sequence |
|---|---|---|
| pinalH02 | MLRLLLRLRRRPRSLSRVRLTSRSPNTPDSDVFPFSLLSDGPS<br>KVYFTKEHEWLAVDEDGTAVVGITEYAQEALGDVVFVDLPE<br>VGTKLSAGDELGAVESVKAASDLYMPVSGEVLEVNEALEDSP<br>GLVNEDPYGKGWLIKMKVAEGVDPENVEGLLDEEEYEKLLE<br>EEH | DDDDDDDPPDDDPPPPPPPPPDDDQQAADCVFPPQVVDPRG<br>FPDWWAFPLAKIWGAHPQQKIWIFGGLVVPVVQDQWDAWF<br>ADDAFDWFAFQDFGTWTHDPVDIDTDTDHHTWGFHDALPVC<br>HVPVRVCNVRRSHSRIGTITGGDPPDDSVPRPRIDDPVVNVVV<br>VVVVD |
| pinalH04 | MLRRLLLRFLRRRRLSSSSARRFSPDDPDPSKPPYSQFKNGPV<br>AVYFTKEHEWIAVDSDGKAVIGITEFAQEALGDVVFVELPEV<br>GTKLQAGDELGAVESVKAASDIYMPVDGEILEVNEALEDNPS<br>LVNESPYGKGWIVKMKVAEGVDPEKIEGLLSLEEYEKLLKEE<br>E | DDDDDDDDDPPDDPPPPPPPPPDDQQAADCVFPPQVVDPRG<br>FPDWWAFPLAKIWGAHPQQKIWIFGGLVVPVVQDQWDAWF<br>ADDAFDWFAFQDFGTWTHDPVDIDTDTDHHTWGFHDALPVC<br>HVPVRVCNVRRSHSRIGTITGGDPPDDSVPRPRIDDPVVNVVV<br>VVVVD |
| pinalH07 | MAEDMIEIKGCKFPKNLYYDVENHVWVRKEEDGTVTVGITD<br>VAQAELGDIVFVDIKEEGKKVKKGKSIAVVESVKWAEDIPSP<br>VSGEIVEVNEALEENPELVNESPYGEGWIVKIKPSDLEADLKE<br>LVSGEEAIDALKEEIEERGIDPEAIVEEHEEKEEE | DPDQWDADLNFIDGQQWWDDLVQQKIWHQDPVRKIWIFGGL<br>QNCLVQFQWDAKDADDFQDWDDAQAARMWTHDPVDIDGDG<br>DLFTFTFHDALPVCHVPVRCCNVPGPPSRTGTIGNGPDPVVSS<br>VVIDGGPVVVVSNNVSCVVVVGRSVVSSVVNCVPPPD |
| pinalH08 | MSKKITFKTEVDHCWIELEGDVVTVGLSPYAQEQLGDIVFIE<br>LTEKETVKAGDTLAVVESVKAASDVYAPVSGEIVERNEEVED<br>EPELLNSDDEEKNWIVKLTDVDQEELAALPKA | DDQDWPAWDDDLQWIWTDGPQKIWIFHHPVVCVVQDAWDA<br>KDADPDQWDAFQDFGMWTHDPRDIDTDTDLHTFGWPDAPP<br>VCRVVVRLCNDPDPVRGIGTMGGPTDPVSVVPGHGD |
| pinalH12 | MTENKIEIKGCKFPKDLYYDVERHIWVRKEEDGTVTVGITD<br>VAQAELGDIVFVELKKEGKKVKAGKSVAVVESVKAAEPIPSP<br>VSGEIVEVNEALEDNPELVNEDPYGEGWIVKIKPSDLEEDLK<br>KLVTGEEAIEAVKEEIEEEGIDAKALAEELEEEEEG | DPDQWDADLNFIDGQQWWDDLVQQKIWHQDPVRKIWIFGGL<br>QNCLVQFQWDAKDADDFQDWDDAQAARMWTHDPVDIDGDG<br>DLFTFTFHDALPVCHVPVRCCNVPGPPSRTGTIGNGPDPVVSS<br>VVIDGGPVVVVSNNVSCVVVVGRSVVSSVVNCVPPPD |

**Supplementary Table 13**: Inference hyperparameters used to generate candidates for wet-lab validation. Initial N denote the number of generated candidates before filtering.

| Task | T2Struct | | SaProt-T | | Initial N |
|---|---|---|---|---|---|
| | Sampling method | Temperature | Sampling method | Temperature | |
| GFP | multinomial | 1.0 | multinomial | 0.1 | 50,000 |
| GFP-g | multinomial | 1.0 | multinomial | 0.3 | 50,000 |
| PETase | multinomial | 1.2 | multinomial | 0.3 | 200,000 |
| ADH | multinomial | 0.8 | greedy | – | 2,500 |
| H protein | multinomial | 1.0 | multinomial | 0.3 | 50,000 |

**Supplementary Table 14**: Sequence identities of experimentally validated active Pinal-designed H-protein candidates from the second validation round to their closest matches in Swiss-Prot and UniRef100.

| Sequence ID | Closest Swiss-Prot ID | Seq. Identity to Swiss-Prot | Closest UniRef100 ID | Seq. Identity to UniRef100 |
|---|---|---|---|---|
| pinalH-R2-01 | Q7N198 | 56.9% | A0A7C4Z5T5 | 69.1% |
| pinalH-R2-02 | B7NHW5 | 67.2% | UPI0005CFBA93 | 67.2% |
| pinalH-R2-03 | B7NHW5 | 66.7% | A0A9X2J1U6 | 69.5% |
| pinalH-R2-04 | A9VNZ4 | 62.0% | C8WX88 | 66.9% |
| pinalH-R2-05 | A4WE56 | 68.0% | UPI0013314338 | 70.4% |
| pinalH-R2-06 | Q8CPW8 | 60.7% | A0A3C1A343 | 65.2% |
| pinalH-R2-07 | Q87I04 | 61.8% | A0A3A2I243 | 63.4% |
| pinalH-R2-08 | Q7MEH8 | 67.7% | UPI0019D4CC84 | 68.5% |
| pinalH-R2-09 | A7ZR13 | 66.1% | UPI0018EF130C | 67.5% |
| pinalH-R2-10 | Q0TDU8 | 68.3% | A0A6I6HFD8 | 68.0% |
| pinalH-R2-11 | Q13SR7 | 56.9% | A0A7V1RD64 | 61.7% |
| pinalH-R2-12 | A4WE56 | 64.3% | UPI001EF94A39 | 64.2% |
| pinalH-R2-13 | A0L104 | 64.3% | A0A2N7VZL8 | 67.5% |
| pinalH-R2-14 | B4RSJ6 | 53.5% | A0A5C6X105 | 53.9% |
| pinalH-R2-15 | A7MR83 | 73.8% | UPI000408B449 | 76.1% |
| pinalH-R2-16 | B7NHW5 | 72.7% | UPI0025AFA695 | 71.2% |
| pinalH-R2-17 | A3D084 | 66.1% | UPI00217DAEC5 | 67.7% |
| pinalH-R2-18 | B7NHW5 | 60.3% | UPI0026241FD0 | 63.6% |
| pinalH-R2-19 | B7NHW5 | 74.6% | A0A7U6GG83 | 70.5% |
| pinalH-R2-20 | Q13SR7 | 70.2% | UPI001D0FE8FE | 71.1% |
| pinalH-R2-21 | Q15PU5 | 64.6% | F7RY66 | 66.9% |
| pinalH-R2-22 | A7MR83 | 68.0% | A0A7C3JGB0 | 68.5% |
| pinalH-R2-23 | Q0TDU8 | 75.6% | UPI0003341EB3 | 75.5% |
| pinalH-R2-24 | Q1AR90 | 54.7% | A0A3M1HJL1 | 60.3% |
| pinalH-R2-25 | Q1AR90 | 57.1% | A0A2M8NU18 | 62.8% |
| pinalH-R2-26 | A1AF93 | 59.4% | A0A0N0VJG4 | 63.5% |
| pinalH-R2-27 | Q311A6 | 65.3% | UPI0018EF130C | 69.1% |
| pinalH-R2-28 | A7MR83 | 63.3% | UPI0013D043E0 | 65.3% |
| pinalH-R2-29 | A1S966 | 59.9% | UPI001F51B1B7 | 65.0% |
| pinalH-R2-30 | B0KD96 | 58.7% | UPI0029902583 | 62.2% |
| pinalH-R2-31 | A4WE56 | 65.2% | A0A2N3J0U8 | 65.8% |
| pinalH-R2-32 | Q7MEH8 | 68.3% | UPI0019D4CC84 | 69.6% |
| pinalH-R2-33 | A9MRH1 | 75.8% | A0A2S4RWI9 | 76.6% |
| pinalH-R2-34 | A1S966 | 67.4% | UPI000CF71096 | 73.5% |
| pinalH-R2-35 | C0PY27 | 70.1% | UPI000F98C1D0 | 73.1% |
| pinalH-R2-36 | A5IR11 | 69.4% | A0A0U1MGG1 | 70.9% |
| pinalH-R2-37 | A6TDR6 | 59.7% | UPI001CFD0610 | 61.6% |
| pinalH-R2-38 | Q1AR90 | 53.9% | A0A349HQD0 | 58.6% |
| pinalH-R2-39 | Q13SR7 | 53.0% | A0A971KXZ2 | 63.8% |
| pinalH-R2-40 | A1S966 | 64.6% | UPI0023DB7C7E | 69.6% |
| pinalH-R2-41 | A9MRH1 | 64.0% | A0A761QG38 | 61.6% |
| pinalH-R2-42 | A5IR11 | 60.5% | A0A7X9EAT0 | 68.1% |
| pinalH-R2-43 | B5XUD4 | 67.0% | UPI001CFD0610 | 65.8% |
| pinalH-R2-44 | A1S966 | 64.6% | A0A8J8E6N3 | 64.0% |
| pinalH-R2-45 | Q0TDU8 | 62.6% | A0A2T5PBH8 | 65.7% |
| pinalH-R2-46 | B7NHW5 | 64.3% | UPI0006817F32 | 61.4% |
| pinalH-R2-47 | Q9WY55 | 59.8% | A0A2G6JBT7 | 68.8% |
| pinalH-R2-48 | Q88CI8 | 58.2% | A0A1H7I0I4 | 62.0% |
| pinalH-R2-49 | A6U8Q4 | 55.1% | A0A931HCK5 | 48.4% |

Note: Closest Swiss-Prot homologs were identified by NCBI BLAST, whereas closest UniRef100 homologs were identified using MMseqs2 (GPU-enabled build, commit 1668032) against the UniRef100 database (release 25_01).

**Supplementary Table 15**: Amino-acid sequences and structure-token sequences of active Pinal-designed H-protein candidates identified in the second validation round.

| ID | Protein Amino Acid Sequence | Protein Structure Token Sequence |
|---|---|---|
| pinalH-R2-01 | MSAVTVPANLKYTESHEWVAPDGTVGITDFAQDLLGDIVFVEYPKGLEVGAEVSAGDEIAVVESVKAASDLYAPVSGKIVEINEAVGDEPELVNEDPYGKGWIFKLELAAGATDDLLTAEAYQALLAED | DPQDADDQQWFADLLQKIAHPQQFIFGDLLVCFVQDQWDAKAADPWPDAFDWFAFQDFGIWTHGPRDIDTDTDLHTFTWHFADVVCRVVVVRCCNVPVPPNVTGTGGPDDPPSCVPTHGSVRVSVVVVVD |
| pinalH-R2-02 | MTEEARRFTKEHEWLKLSEDGKTVTVGITDHAQEELGDIVFVELPEAGTALSADEDAAVVESVKAASDIYAPVSGEVVEVNEEVVDSPELVNEDPYGAWLVKIKLSDAAEVESLLDEEAYEALLAEEE | DQDFWWFAALQAKIWGQDPVRFKIWIFGDLVVCVVQDAWDAKAADDFFDWAAFQDFGIWIHGPVDIDTDGDNHTFTWHDADVVCRVPVRVCNVCRPVHTGTMGGHPDSVVRVVGHTPVRVVVSVVVVD |

Supplementary Table 15 – *Continued from previous page*

| ID | Protein Amino Acid Sequence | Protein Structure Token Sequence |
|---|---|---|
| pinalH-R2-03 | MSNSSELKFTKEHEWVKIEGNIAYVGISEYAQDQLGDIVFVELPEVGKTVKKGDDCAVVESVKAASDIYAPVSGEVVEVNEELEDNPELINEGSDIWIFKIKMSDPSELENLLDEAGYEALLEEEE | DDLPQDWWAALLAKIWGDDPQKTFIFGDLVRCVVQDAWDAKAAADAFDWDAFQDQGMWIHDPVDIDTDTDLATFGWHDADPVCHVCVCVCVVDGDDGGTMGRHPDPVSVVNIHDPVRVVVSVVVVD |
| pinalH-R2-04 | MANPEDLKYLKEHEWVKIDGDVAYVGITDHAQEELGDIVFVELPEVGAKIKAGEEFGVVESVKAVSDLYAPVSGEVVAVNEALSDSPELVNEAPYGAWLVKIKVDPASLEAGLSKLLDEEAYAALIEE | DQDDPDWWFAPLAKIWDDDPQKIFIFGDLVVCVVQDQWDDKFADDAQDWFAAQDFGTWIHDPVDIDTDGDLWTFGFHDAPPVCHVPVNCCNVPRPVDTGTMGRHDPVGHVNRVVRTDDPVVVVVVSVD |
| pinalH-R2-05 | MANALRFTKEHEWIKLESANTVRVGISEHAQDQLGDMVFIELKEVGTTVSAGDDIAVVESVKAASDIYAPVSGEVVAVNEALEDSPELVNEDPEGEGWIVRIKIADASELDSLLDEEAYAALLEEEE | DFFAWFADPLAKIWGDPDQFKIWIFGHPVNCVVQDAWDAKDADDFFDWDAFQDFGMWTDGPVGIDTDTDLAGFTFHDALCVCHVVVCCCNVPGPPSRTGTMTGHPGPVSRVPTHGPVRVVVSVVVVD |
| pinalH-R2-06 | MSDNNKSKKLSPFEIPEELLYSKEHEWVRLEDGLAVVGITEFAQEQLGDIVFVELPEVDSELKVGDGVAVVESVKAASDLYSPVSGEVVEVNEELEDEPELVNESPYGEGWLVKIKPSSDELPEDLMTAEEYEEYLEKEAEEEGLGE | DDPPPVPPPPDVWDFDQQWWAAPLQKTWHDDPQKTFIFHGLLVCVVQDAWDAKAADDAQDWFAFQDFGIWIHDPVDIDTDTDQATFGWHDALPVCHPCVRCCNVPGPPSRTGTMGNGPDPDGDPRIDGSVRVRVVVQVVCVVVVVRD |
| pinalH-R2-07 | MSKELRFTDSHEWLRPDGEDGTVTIGLSEYAASELGDIVFVDLPEVDTEVKAGDTLAVVESTKAVEDIYSPVSGEVIAVNEALEDNPELVNEDPYGEGWIVKLKLTDDTDEFEALMSEEEYLALLS | DDAFWFADPLQKIWGQPPPFQKIWIFHHLVNLVVLFDWDAKAADDAQDWAAAQDFHIWTDGPVGIDTRGDNATFGFHDALPVCHVPVCCCNVPGPPSRTGTMTHHPDDCPRVVVGHGPVVVVVVVD |
| pinalH-R2-08 | MSDGLKFSESHEWVKDNGDGTVTVGITEHAQEQLGDIVFVDLPEVGATLKAGETLAVVESVKAASDIYAPVSGEVVAVNEALSDSPELVNEDPYGDGWLVRVKPSSPPPAELIDEEEYLAELE | DQAFWWADPQQKIWGDPVPQKIWIFGDLVVCVVQDAWDAKAADDAQDWAAFQDFGIWIHGPVDIDTDTDQATFGWHDALPVCHVPVRVCNVCRPPSVIGTMGRGPDDGDPRIDGPVVVVVVVD |
| pinalH-R2-09 | MSAHNVPEGLKYTKEHEWIKIEGDVATVGITDYAQELLGDMVFVELPEVGATLSAGDEAGSVESVKAVSDLYAPAAGEVVAVNEALEDAPELVNEDAYGSWIIKIKLSSEPEDLLDAAAYAELIG | DPDFDQDAQWWADLLAKIWHDDPQKIFIFGGLVNCVVQDQWDDWFADDQQDWFAFQDFGTWTHDPVDIDTDTDQHTFGWHDADVVCRVVVRVCNVDRPVHTGTMTRHPDDGDPIHGRVRVSVSSD |
| pinalH-R2-10 | MAANIAELKFTKEHEWIKKEEDGVVTVGISEHAQEELGDLVFVELPEVGKAVTAGEAFGVVESVKAASDIYAPVSGEVVAVNEAVAESPESVNEDPYGAWIFKVKVSDESELEGLISGSEYEELTG | DFDDLQPWWAALLQKIWDDDPPQKIWIFGDLLVCVVQDQWDAKAAADFFDWAAAQDFRIWIHDPVGIDGDGDLATATWHDADVVCRVPVRVCNVCRPVHTGTMGRHPDPVSSVNIDGSNVVVVSRD |
| pinalH-R2-11 | MAVPTDLKYTKTHEWVRVEGDTATVGITDHAQEELGDIVYLEFLKKAGDALAAGDVFAVVESVKAVSDIYVNGDVVEVNEELAASPEIINDSPYGAWLVKVKPADLEAALAELLDADGYSALLEEE | DDQAQQWWAFLQAKIWHDDPQKIWIFGDLLVCVVQDQWDAKDFPDAQPDWAAFQDFGIWIHGPVDIDTDTARFTFHDAPPVCRVCVRCCNVCRPPDTGTMGRGPPPVVSRVNTHGSVRSVVSVVRD |
| pinalH-R2-12 | MSEIPANLKFLKTHEWVRKESDDTYLVGITDHAQEELGDIVFVELPEVGTTVKAGEVCGVIESVKAVSDVYAPVSGEVVAVNEALEDNPELINEDPYGDGWLVKIEVPADADLSHLIEGDDYAALLGE | DFDQDQFWFDDLLAKIWHDPDPFKIKIFGGLVHCVVADQWDAKAAADFFDWDAFQDFRIWTHGPVDIDTDTDLFTFTFHDALPVCHVPVRVCNVPGPPSNTGTMGMHTPPTDRVVTHTDPRVCVSVVD |
| pinalH-R2-13 | MANEKKFTKSHEWVKVEEDGTVTVGITDHAQEQLGDIVFVELPEVGDTITAGDSFGSVESVKTVSDLYAPVSGEVVAVNEALVDSPELVNEDAEGAWLFKVKPSGEPEDLLDEAAYEALIAEEEH | DAQAWWDAPLAKIWGQHPVRKIWIFGDLLVCVVQDQWDAWFAADQFDWAAFQDFGIWIHDPPDIDTDGDQATFGWHDADVVCRVPVRVCNVDRPPDTGTIGNGPDDTPPIHGPVVNVVSVVVVVD |
| pinalH-R2-14 | MSNGKRYTATHEWVEPEGGDGVVRVGLTDHAVEKLLEELGGPEIVFLELPEVGKELTAGDEIVTLETAKAVIEIYAPVSGEVVAVNDALVEDPELVDEDPYGDGWLLRLKLDDPADLEALLDEEEYEAFLEEGG | DDFAWFADPLQKIWGDPVQQQKIWIFHDVVNLVVLCVVLVAWAFPAKDADDFFDWAAFQDFGIWTDGPVDIDTDTDQATFTFHDALCVCRVPVVCCNVCVPPSNTGTITRGPDSCSSVVTHGPVRVVVCVVVTD |
| pinalH-R2-15 | MTDNSTINLSDLRFSESHEWLRKEADGTYTVGITDHAQEELGDVVFVDLPEVGATLEKGDDIGVVESVKAASDIYAPVSGEVVAVNEALSDSPELVNSSPYGEGWLIKLKPGADAEEELAALLDAEAYEALCEEE | DDPPPPPPQQQWWADLLQKIWGADPVRKIFIFGDLLVCVVQDQWDDKFADDQFDWFAFFDFGIWIHDPVDIDTDTDLHTFGWHDALPVCRVVVRCCNVPRPPSRTGTIGNGDPCSVVSRVPTHGPVRVVVSSVVD |
| pinalH-R2-16 | MSNIPAELKYSKEHEWVRIEGDVATVGITEHAQEELGDIVFVELPEVGAEIEAGETCGVVESVKAASDIYAPVSGEVVEVNEALSDSPELVNSDAYGAWLFKVKVTSDEQDEDDLLDADAYAALTE | DFPQDQQWWADLLAKIWHDDPQKTFIFGGLLVCVVQDAWDDKFAADQFDWADAQDFGIWIHDPVDIDTDGDQATFTWHFADVVCHVCVCVCVVPVPVDTGTMGGGDDPDGDPCNIDGSVRVVVSRD |

Supplementary Table 15 – *Continued from previous page*

| ID | Protein Amino Acid Sequence | Protein Structure Token Sequence |
|---|---|---|
| pinalH-R2-17 | MSDTGYTVPEGLKYSDEHEWVRQEEDGTYTVGITDFAQDLLGDMVFVELPEVGDDLKAGEPFGSAESVKTVSDLYAPVSGEVVAVNEALEEEPELVNSSPYGEGWLVKVKPDPESLKGLLDAEEYAKLIDGEEAEAIGDAS | DDPPVFDQDAQWWAAPLAKIWGADPVRKIWIFGGLVRCVVQDQWDAWFAADAFDWADAPDFGIWTHDPPDIDTDHDLATFGWHDALPVCRVVVRCCNGPGPGSRTGTIGRGDPVRSPRTDHSVQVVCRRLVNVCVVVVNPD |
| pinalH-R2-18 | MSTSASTGAETAYEVPEDLRYSRSHEWVDASGDALSGSGPVIVGITDHAQQLLGDIVFVDLPEVGDTLTVGEPFAVVESVKAASDLYAPISGKVVEVNEALSDSPELVNEDPYAEGWIARVELSDPSQLSALLDAEQYAELVEDDDDGEA | DDDDDDDDPPDDFDQDQQWWADLLQKIWRPPDCCVVDLAWTKIFGGLLVCVVQDAWDAKAADDQFDWADAQDFRIWIHDPVDIDTDGDLATFGFHFALPVCHVPVRCCNVPVPPSNTGTMGGHPDSVRSVVTDGSVRVSVSVVCVVVVVD |
| pinalH-R2-19 | MAQLKYTKEHEWVKLEDDGKTATVGITDHAQELLGDMVFVELPEVGPVSAGDDIAVAESVKAASDVYAPVSGEVVEVNEALSDSPELVNEDAYGAWIFKIKLSDPSDVDALLDEAAYAALIAE | DFFWFAALQAKIWTQDPVLFKIFIFGDLVNCVVQDAWDAKAAADFFWDAFFDQGMWTHDPVDIDTDTDLHTFTWHDADPVCHVCVCVCNVDRPVHTGTMTGHPDSCSRVVTHGPVRVVVSVVD |
| pinalH-R2-20 | MSDLSDLEFTEQHEWVLTEADGSKTIGITDHAQEELGDMVFVELPEVGTSVSEGETCAVVESVKAASDIYAPVSGEIVAVNEAVADAPELVNEDPYGSWLFKIK | DDPPVQWDADPQQKTWDQDPVRKTFIFGDLVNCVVQDQWDDKAAADFFDFDAFQDFGIWTHDPVDIDTDTDNHTFTWHDADVVCRVPVRVCNVCVPVHTGTIGD |
| pinalH-R2-21 | MTEEKIVEVKGGNTEFVVPEDRYYAKSHEWVKIEEDNTATIGITDYAQELLGDIVFVELPDLGKEVSAGDTVAVVESVKAASDIYAPLSGEIVAVNEELEDEPELINEDPYGAWLVKIKASDPEEDLENLLDAEEYAELLEEEAEE | DPPFDWDWDDDPDDIFIFGQQWWAALVQKTWHQDPQQKIWIFGGLVVDVVQDQWDAKDADDAFDWDAFQDFGMWTHHPVDIDTDTDLATFTFHDADPVCHVVVNCCNVCVPVDTGTMGRGDCRVVSRVNTHGRVRVRVSVNVVVVD |
| pinalH-R2-22 | MSAPEGLKYSKEHEWVRIEGDVATVGITEYAQEELGDIVFVELPEVGATIAAGDAFGVVESVKAVSDLYAPVSGEVVAVNDELEDEPELVNEDPYGTWIFKVKLSDVEADTADLLDAEAYDALIEE | DFDPDQWWADLQAKIWRDDPQKIFIFGGLVVCVVQPAWDDKFAADQQDWFAFQDFGTWIHDPVDIDTDTHLATFTWHDADVVCRVVVNVCNVCVPVDTGTMGRGPDPPVSRPNIHGRVRVRVSVVD |
| pinalH-R2-23 | MADIKGCEVPEDLKYHGEEHEWVRVEEDGTVTVGITEHAQEQLGDIVFVDLPEVDTTIEAGDTLAVAESVKTASDIYTPVSGEIVAVNEALEDSPELVNEDAYGAWIVKIKLSDESELEALLDAAAYEALLEEE | DDAQLNFDDDQQWWAQQVQQKIWHQDPVQKIFIFGGLSVCVVQDQWDAKAADDAQDWDDAQDFHIWTHDPVDIDTDTHLFTFTFHDADPVCHVPVRVCNVPRPVDTGTIGRGPDPVSSVVTDGRVRVVVSSVVD |
| pinalH-R2-24 | MSSESDLIELKIGEETIRIPKDRLYTESHEWVRIDGDEATVGITEFGQELLGDIVFVELPEVGTKLKAGDTLAVVESVKSVSDIYAPADLEILEVNEDVLENPELVNSDPYGKGWIVKVKLLGELDREGDGELLDAEEYYALLEKELNPEE | DDPPVQWDWDDDPPDIFTFGQQWFAAPLAKTWHDDPQKTFIFGDLVNDVVLDAWDAKDADDAQDWAAFQDFGMWIDHPVDIDTDTDQHGKGFHDALVVCRVPVVCCNPPVPPSNTGTMIGGDDDGPCVPPPSIDGSVRVVVSVVCVVPPDD |
| pinalH-R2-25 | MADLNLPKDLLYSESHEWVRIEGDTATVGITDLAQKLLGDIVFVDLPEVGDELKKGESFGSVESVKTVVDIYSPVSGTVVEVNEEVVENPELINSDPYGEGWIFKVKVKDLEEDKKELMDAEEYEKILEEEGEELAKIPHGH | DLPQPDDQQWWADPLAKIWHDDDQKIFIFGDLLVLLVQPDWDAWDAADAFDWADAQDFRTWTDGPQGIDTHGDLATFGFHFALVVCRVPVVCCNPPVPPSNTGTMGRGDCRVVSVVNIDGPVVVSVVSVVVVVVVSSPRDPDD |
| pinalH-R2-26 | MEVNGCEIPDDLLYNAEEHEWVRKEDDNTVSIGITDFAQELLGDIVFVEFEVEVGDTVKAGDSFAVVESVKTASDVYAPVSGEVVAINEELEESPELVNDSPYEAWIVKLKNVDEEDLSELVDAAEAEEELEEEAE | DAALNFDDDQQWWDQLVQQKIWHDPDPFKIFIFGGLLNCVVQDQWDAKDFPDDQQDFDAFQDFGMWIHDPVDIDTDTDLWTFGFHDADPVCRVVVRCCNVPVPVDTGTMGGPTDPVSSVNIDGPVVCRVSVVVSSD |
| pinalH-R2-27 | MSNNIPAELKYAESHEWVRIEGDVGTIGITDFAQEQLGDMVFVDLPEVGTEVTAGEPFGSVESVKAASDLYAPVSGEVVEVNEDLEDSPEKVNEDPYGVWFFKVKLSGDTDDLLDAAEYEKLVEGE | DPQPQDQQWWADLLQKIWHDDPQKTFIFGGLLNCVVQDAWDDWFAADFFDWAAAQDFRTWTHDPVDIDTDGDLATFTWHDADVVCRVVVRVCNVCVPVDTGTMGGGPDDPPSIHGSVRVCCSRVVD |
| pinalH-R2-28 | MSIPSELRYTKEHEWVRVESDGTVTIGITEHAVEELGDIVFVELPEVGTELKAGDAFGSVESVKTASDLYAPVSGEIVAVNAALEDSPELVNDKAESEDGWFFKVKPSEEAAGEDAAELLSAEAYEALIG | DDQDQQWFAALQQKIWHQDPVQKIFIFGDLVVCVVQDQWDDWFAADFFDWAAFQDFGIWTHDPVDIDTDGDNATFTFHDADPVCHVCVRVCNVARVDPPHTGTMGRHPDDDDGHDPVRGDGSVRVVVSVD |
| pinalH-R2-29 | MTETSTSTVINGHKIRAELRYDAEQHIWIRIEEDGTVTIGITDHAAEELGDLVFIELPEVGAEVEVGKSAAVVESVKAASDVYAPVSGEVVAVNEEVIDNPELINSDPYGAWIVRVKPEDEAEVEAALEELLDADGYANFLEEE | DDDQPPFDADLNFTDGQQWWDDLVQQKIWHQDPVRKIFIFGGLQNCQVQDQWDAKDADDAFDWFDFQDFRMWIHDPVDIDTDGDLATFGWHDADVVCRVPVNCCNVPRPVDTGTMGRHPDPVRVVVSRVVTHGRVRVSVSVVVD |
| pinalH-R2-30 | MSVEIGGCTIPDDRLYSPDDHEWVRIEGDEATVGITEYAQSLLGDIVFVDLPEVGTELEKGETAAVVESVKAASDLYSPVSGEVIEVNPVVENPELVNEDPYGEGWFAKVKVTDESELEGLLTAEEYLEYLIEETGVEAE | DWDDDLNWIQDQQWWADLVQQKTWHDDPQKTFIFGGLVVCSVQDAWDAKAADDFFDWADAQDFHIWTHDPVDIDGDGDLATFTFHFALPCRVPVRCCNVPVPPSRTGTMGGGDDPCSCPPIHGSVRSVVSVCVVVVTPDD |

Supplementary Table 15 – *Continued from previous page*

| ID | Protein Amino Acid Sequence | Protein Structure Token Sequence |
|---|---|---|
| pinalH-R2-31 | MAQDFAELKFTEEHEWIRIEEEDGKKYATVGITEFAAEALGDVVFVELPEVGKEVKAGEDCAVVESVKAASDVYAPVSGEIVEVNEALEDSPELVNSDPYEAWMFKIEMSEEPEGLLDAEEYAALTEDE | DFDDLQPWWAALLQKIWDWDDDPNWIKIFIFGDLLVLCVQPQWDAKDAADFFDWDAAPDFGMWIHDPVDIDTHGDQATATWHDADPVCHVPVRCCNVPVPVRTGTMGGHPDDGDDIHGSVRVSVSRVVD |
| pinalH-R2-32 | MSEESEYEVPEDLLYTEEHEWVRDEGDGVVTIGITEHAQELLGDMVFVDLPELNAEITAGEAFGTVESVKAVSELYSPVSGKVVEVNEALDDSPELVNEDPYGEGWIVRVKVSDLEEEKANLMDAEEYIELLEEE | DDDDPPFDFDQQWWAAPQAKTWHDPDPQKIWIFGGLVVCVVQDAWDAWDAADAQDWAAFQDFGTWTHGPVDIDTDGDNAGFTFHDALPVCHVCVCCCNVPRPPSRTGTMGRGDCSVVSVVNIHGRVVVSVVVVVD |
| pinalH-R2-33 | MSELKFTDEHEWIKLEDDGTVTVGISDHAQELLGDVVFVELPEAGTTVSAGDDCAVVESVKAASDIYSPVSGEIVAVNEALSDSPELVNSSPYDEGWIFKIKPSEADEVESLMDEEAYEALLEEEE | DFFWFAAPVQKIWTQDPVLKIWIFGDLVNCVVQDQWDDKAADDFFDWDAQQDQGMWIHDPVDIDTDGDNATFTWHDALPCCHVCVNCCNVPGPPSRTGTMGNGPDRCVRVPGHGPVRVVVSVVVVD |
| pinalH-R2-34 | MAEINGCQFPDELYYDVENHIWLRKEADGTYTVGITDHAQELLGDLVFVELPEVGTEVSAGEDCAVAESVKAVGDIYAPVDGEIVEINEALEDSPELVNSDPYGEGWLFKIKPSDESGFEDLIEGEEVIAALKEEA | DDADQNFDDDQQWWDPLVLQKIWHQDPVRKIWIFGGQVVCQVQDQWDAKDAADAQDWDAFQDFGMWTHDPPDTDTDGDLHTFGWHFALPVCHPCVRCCNQPRPPSRIGTMINHPDPPSNPNIDGHPVVVVVVSVVRD |
| pinalH-R2-35 | MSNLRYTKEHEWLRKEADGSVTVGITDHAQELLGDVVFVELPEAGKKVAAGEDFGVVESVKAASDIYAPVAGEVVAVNEALEDNPELVNSDAYGAWIFKIKPADASAASSELLDEAAYEALIG | DAPWFADLLAKIWHQDPVLKIWIFGDLVNCVVQDAWDAKAAADFFDWDAQQDFGIWIHDPRDIDTDGDNHTFTWHDADPVCHVPVCCCNVPVPVDTGTITRHPDSCPRVPPGHDPVRVVVSVD |
| pinalH-R2-36 | MAETYTIGAGEVPDELKYSKSHEWVRVEGDVATVGITDYAQELLGDMVFVELPEVGDTITAGEPFGSVESVKAVSDLYAPVSGEVIEVNEALEDSPELVNEDPYGDAWLVKVKLSNPEELEELLSAEEYAEAIGEA | DFDWDDQDQFDDDQQWWADLVAKIWDDDPQKIFIFGGLNRCVVQDAWDDWFAADQFDWAAAQDFRIWTHGPHDIDTDGDLATFGFHDALPVCVVCVSCCNTPRSPSGTGTMGRGPCCVSSVVTHGSVVNCVSVVRD |
| pinalH-R2-37 | MATINGFELNEDLLYSEDKHEHVWVKPLDETEVTVGITDYAQELLGDLVFVELKKVGKTVEAGEPFATVESVKAVEDIYSPVSGEIVAVNEELEDSPELVNEDPYGEGWIVKVKLDEPVDVDALKASLLDAEAYVALVEENS | DDADLNFDDDQQWFWDDDPQQIKTWRDDDQFKTWIFGGLSVCQVQDQWPAKDADDFFDWADAQDFGMWTHDPRDTDTDGDLATATWHDADPVCHVCVRCCNVPRPPSRTGTMGGGPGTDPVVVSRVPTDGRVRVSVVSVVRD |
| pinalH-R2-38 | MEIKGHEIPDELLYLVDDHIWVKVEGNTATIGLTEHAQEKLGDIVFVELKEVGTEVEAGEDVAVVESVKTASDVYTPVSGEIVEVNEALEENPELVNEDPYEAWIVKIKPTNLEEEKPKLVTAEEAEDILEEEAEEL | DAALNWDDDQQWWALLVQQKIWHDDPQKIWIFHGQLRCVVQDQWDAKDADDFFDWDDAQDFGMWIHDPVDIDTDGDQFTATFHDADPVCHVPVRCCNVPRPVHTGTMGRGDCCVVRVVSTDGCVVCRVSVNVVVVVD |
| pinalH-R2-39 | MEIGNYEVPTDLLYDVEDHVWVKEEEGNVYTVGITDLAQEALGDIVFVELKEPGTKVKAGKSLAVVESVKTASDVPSPVSGEIVEVNEELEDSPELVNEDPYEAWIVKIKVDEEELEKLVEGGEELLAEVEGE | DDQQPWDDDQQWWADLVQQKIWHDDDDQKIFIFHGLSVDVSQDQWDAKAADDFQDWDAAQDARIWTHDPVDIDGDGDLATFTFHDADPVCHVPVRVCNVPVPVDTGTIGRGDPVSSVVTDGIDCVSVCVNVVD |
| pinalH-R2-40 | MSEEKEKITGEIPADRKYSESHEWVKIEGDVGTVGITDYAQELLGDDVVFVELPEVGKTVSAGEDFGVVESVKAASDIYAPVSGEIVEVNEALEDNPELVNSDPYGEGWLFKMKITDEDELENLIKGEEYYEAVGEEHEVEGA | DDPDPWDKDFDFDQQWFAALLQKTWHDDPQKTFIFGGLVVCVQQDQAWFAKDAFDAQDWDAAQDFGIWIHDPVDIRTHGDLATFGWHFALPVCRVPVCCCNVPRPPSNIGTMTRGDDVVSVVNTDHYPRVVVSVPDHGDRHRD |
| pinalH-R2-41 | MSTETIKKYTEEHEWIKLEGDTVTVGITDHAQEQLGDVVFVELPEVGTELSSAGDSVAVVESVKAASDIYSPIKGEVTAVNEALEDSPELVNSDPEGDGWIFKMKLTGDGDDEDDALDSLLDEDAYEALIE | DDQDWDWWADLQAKIWIDRPQKIWIFGDLVVCVVQDQWDDKAAADFFDFFDFQFDFGMWIHDPHDIDTDTDLAGWGWHFALPVCHVPVRCCNVPRPPSRTGTMTGHDDPPDPRVVSVVPTHDPVRVVVSVD |
| pinalH-R2-42 | MSGPEGLKFTKSHEWVKVDGDVATVGITDHAQEELGDIVFVELPDVGKKLSAGDTFGVVESVKAASDIYAPVTGEVVEVNEAVVDSPELVNEDPYGAWLVKVEVSDLSELDALMDADGYEALVE | DQDPPAWWADLQQKIWDDDPQKIWIFGHLVVCVVQDQWDAKDADDFFDWAAFQDFGIKIHGPVGIDTDGDLWTFTFHDADVVCRVPVSCCNVCRPVHTGTMGRHPDCVSRVPTDHSVVVVVSVD |
| pinalH-R2-43 | MSSDFKYLKGHEWIRKEDDDTVRVGITKFAQDLLGDIVFVDLPEVGTEVKAGEPFAVVESVKAVSDLYAPVSGEIIAVNEELDDNPELVNSDPYGEGWIFKMKVSDPSELEAALKDLYTDLEAAEAWDEKHPDEAEEIKEELGLA | DAQQWWDDQLQKIWGDPDQFKIWIFGDLLVDVVQDQWDDKAADDFFDWAAAQDFRIWTHGPVDIDTDGDLHTFTWHFALPCCRVPVVCCNLPGPPSRTGTMTGGPHSVSVVVSVVVTHSDDVSSNVSCVVVVCVSVVVCVVSVVD |
| pinalH-R2-44 | MAAPTNVPEGLRYDKEDHIWVRLEADGTVTIGITNHAQELLGDIVFVELPEVGTEVSKGETAGVVESVKTASDVYAPVSGEIVAVNEDLEDSPETVNDDPYGAWIARVKPSDLDEELSKLLDAEAYQKVLDEE | DQDDQDQDAQWWADLVQLKIWHQDPVQKIWIFGDLVVCVVQDQWDDKAADDFFDWFAFQDFRIWIHDPVDIDTDGDLATFTWHFADVVCRVVVSVCNVDVPVGTGTMGRGPDDVPSSVVTHGRVVVSVSSVVD |

Supplementary Table 15 – *Continued from previous page*

| ID | Protein Amino Acid Sequence | Protein Structure Token Sequence |
|---|---|---|
| pinalH-R2-45 | MSLTAEDLLFLKEHEWVRKEEDGTYTVGISDYAQEELGDIVFVELPEVGDELEVGESCAVVESVKAASDVYAPVNGEVIAVNDALSDDPELVNEDPYGAWIFKVEPSEEEEDAEELAAKFGPLDAEEYEASIAE | DFDAQQFWFAFLVAKIWHQDPVRKIFIFGGLVVCVVQDQWDDKAAADAFDWFDAQDFGIWIHDPVDIDTDGDFAGFTWHDADVVCHVPVNVCNVCRPVHTGTMTRHPTDTDGPVRSCVRRVTDGSVRNVVSVVD |
| pinalH-R2-46 | MAEVKGCEIPDELYYDPEEHIWVRVEDDGTASVGITDIAQKLLGDMVFVELPEEGTKVEAGDDIAVVESVKTASDVIVPVSGEVVEVNEELEDSPELVNSDPYEAWIVKIKLSDESELEELKSGDAAVAALEEYAEELGLSA | DDAALNFDDDQQWWDQLVLQKIWHQDPVQKIFIFGGLLNCVVQDQWDAKDADDFQDWDDQQDFGMWTHDPVDIDTDGHRFTFTFHDAPPVCRVPVRCCNVPRPVDGGTMGRGPDPVSSVVIDGHPRVNVSNNVVCVVVVGHD |
| pinalH-R2-47 | MSLLILLLSSNGERFKFKDHEWIKVEGGTGTVGITDFAQDLLGDIVYVELPEVGSEVSKGEDIGSVESVKAASDLYAPISGEITEVNEALLSSPELVNEDPYGEGWFFKIEITDESELKELMDEEAYEKLIEED | DDLLPPPPVPQDWDWDDDPQKIWIDAVQKIWIFGRLNVCVSQDQWDDWDAADAFDWDAFQDFGTWIHDPVDIDTDGDLHTFTWHDALCVCRVVVCCCNVPGPPSRTGTMGGDPDPVSRVVTHDPVRVVVVSVVD |
| pinalH-R2-48 | MSLSELRYSESHEWVKIESDKTVTIGITDFAQKELGDMVFVELPEVGTEIEAGDSLGVVESVKAVSDLYTPVAGEVVAVNEEVLDNPESINEDPYDSWLVKIKTEEPVDASDLLDAEAYAASIKDDE | DDQQQWFADLLQKIWHDPDQFKIWIFGDLLVCVVQDQWDAWDADDAFDWAAFQDFGTWTHDPVDIDTDTHRHTFGWHDADVVCRVPVSCCNVPVPVDTGTMTGHPGGDDRVNTHGSVRVSVSCPVVD |
| pinalH-R2-49 | MAKEYWVKKEGDEVIVGLTEEAQEELGDIVYVEILEGDEVKKGDPIAVVESVKAVSDVYSPVDGEIVEKNEEVLEDPSLLNSDDEEKNWIVKLTGGAKAEDIEL | DPDQWDWDDDPQKIWIFGDLVNDVVQDQWDAKDADDDFWDAAQDFGMWTHDPVDIDTDGDLGGFTFPDAQVVCRVPVRLCNDPDPVRVTGTMTRGPIDPVSMGD |
